# Giant transposons promote strain heterogeneity in a major fungal pathogen

**DOI:** 10.1101/2024.06.28.601215

**Authors:** Emile Gluck-Thaler, Adrian Forsythe, Charles Puerner, Cecilia Gutierrez-Perez, Jason E. Stajich, Daniel Croll, Robert A. Cramer, Aaron A. Vogan

## Abstract

Fungal infections are difficult to prevent and treat in large part due to strain heterogeneity which confounds diagnostic predictability. Yet the genetic mechanisms driving strain-to-strain variation remain poorly understood. Here, we determined the extent to which *Starships*—giant transposons capable of mobilizing numerous fungal genes—generate genetic and phenotypic variability in the opportunistic human pathogen *Aspergillus fumigatus*. We analyzed 519 diverse strains, including 11 newly sequenced with long-read technology and multiple isolates of the same reference strain, to reveal 20 distinct *Starships* that are generating genomic heterogeneity over timescales relevant for experimental reproducibility. *Starship*-mobilized genes encode diverse functions, including known biofilm-related virulence factors and biosynthetic gene clusters, and many are differentially expressed during infection and antifungal exposure in a strain-specific manner. These findings support a new model of fungal evolution wherein *Starships* help generate variation in genome structure, gene content and expression among fungal strains. Together, our results demonstrate that *Starships* are a previously hidden mechanism generating genotypic and, in turn, phenotypic heterogeneity in a major human fungal pathogen.

**Importance:** No “one size fits all” option exists for treating fungal infections in large part due to genetic and phenotypic variability among strains. Accounting for strain heterogeneity is thus fundamental for developing efficacious treatments and strategies for safeguarding human health. Here, we report significant progress towards achieving this goal by uncovering a previously hidden mechanism generating heterogeneity in the human fungal pathogen *Aspergillus fumigatus*: giant transposons called *Starships* that span dozens of kilobases and mobilize fungal genes as cargo. By conducting a systematic investigation of these unusual transposons in a single fungal species, we demonstrate their contributions to population-level variation at the genome, pangenome and transcriptome levels. The *Starship* compendium we develop will not only help predict variation introduced by these elements in laboratory experiments but will serve as a foundational resource for determining how *Starships* impact clinically-relevant phenotypes, such as antifungal resistance and pathogenicity.

## Introduction

Infectious diseases caused by fungi pose a grave threat to human health and society. The World Health Organization recently coordinated a global effort to prioritize research among fungal pathogens based on unmet research needs and public health importance (1). Among the pathogens deemed most important for research include *Aspergillus fumigatus*, a globally distributed opportunistic human pathogen causing disease in an estimated 4 million people yearly (2, 3). Several infectious diseases are caused by *A. fumigatus*, including invasive pulmonary aspergillosis, which manifests primarily in immunocompromised individuals with mortality rates of up to 85% (2, 4). The treatment of *A. fumigatus* is complicated by its remarkable variation in virulence, resistance to antifungals, metabolism, and other infection-relevant traits (3, 5–8). Strain heterogeneity confounds “one size fits all” therapies and poses a significant challenge for developing efficacious disease management strategies (9). Recent evaluations of the *A. fumigatus* pangenome have revealed extensive genetic variability underlying variation in clinically-relevant traits such as antifungal resistance and virulence, yet in many cases, the origins of such variation remain unexplained (10–12). Determining the genetic drivers of strain heterogeneity will help accelerate the development of strain-specific diagnostics and targeted therapies.

Mobile genetic elements (MGEs) are ubiquitous among microbial genomes and their activities profoundly shape the distribution of phenotypic variation (13). MGE transposition generates structural variation that directly impacts gene regulation and function (14–16), and many elements also modulate genome content by acquiring genes as “cargo” and transposing them within and between genomes, facilitating gene gain and loss (17, 18). The ability to generate contiguous genome assemblies with long-read sequencing technologies has dramatically enhanced our ability to find new lineages of MGEs (13, 19). For example, we recently discovered the *Starships*, a new superfamily of MGEs found across hundreds of filamentous fungal taxa (20–22). *Starships* are fundamentally different from other fungal MGEs because they are typically 1-2 orders of magnitude larger (∼20-700 kb versus 1-8 kb) and carry dozens of protein-coding genes encoding fungal phenotypes (23). In addition to flanking short direct repeats, all *Starships* possess a “captain” tyrosine site-specific recombinase at their 5’ end that is both necessary and sufficient for transposition (21). Several *Starships* mediate the evolution of adaptive phenotypes: for example, *Horizon* and *Sanctuary* carry the ToxA virulence factor that facilitates the infection of wheat, and *Hephaestus* and *Mithridate* encode resistance against heavy metals and formaldehyde, respectively (24–27). *Starships* have been experimentally shown to horizontally transfer between fungal species (28), and an increasing number of comparative studies implicate them in the horizontal dissemination and repeated evolution of diverse traits (20, 21, 25–27). *Starships* may also fail to re-integrate into the genome during a transposition event, leading to the loss of the element, its cargo and its encoded phenotypes (21). Through horizontal transfer and failed re-integration events, *Starships* contribute directly to the generation of rapid gene gain and loss polymorphisms across individuals. However, we know little about how *Starships* drive genetic and phenotypic variation in fungal species relevant for human health and disease.

Here, we conduct the first systematic assessment of *Starship* activity and expression in a human fungal pathogen to test the hypothesis that these unusual transposons are a source of strain heterogeneity. We reveal that *Starships* are responsible in part for generating key instances of previously unexplainable variation in genome content and structure. Our interrogation of 508 diverse clinical and environmental strains of *A. fumigatus,* combined with highly contiguous assemblies of 11 newly sequenced strains, enabled the unprecedented quantification of *Starship* diversity within a single species and even within the same “reference” strain. We leveraged the wealth of functional data available for *A. fumigatus* to draw links between *Starship*-mediated genetic variation and phenotypic heterogeneity in secondary metabolite and biofilm production contributing to pathogen survival and virulence. We analyzed multiple transcriptomic studies and determined that variation in *Starship* cargo expression arises from strain- and treatment-specific effects. By revealing *Starships* as a previously unexamined mechanism generating phenotypic variation, our work sheds light on the origins of strain heterogeneity and establishes a predictive framework to decipher the intertwining impacts of transposons on fungal pathogenesis and human health.

## Results and Discussion

### Data curation and long-read sequencing

*Starships* are difficult (but not impossible) to detect in short-read assemblies due to their large size and frequent localization in regions with high repetitive content (20). The wealth of publicly available short-read genomes is still of high value, however, due to the breadth of sampling they provide combined with the still relevant possibility of finding *Starships*, especially in higher quality short-read assemblies such as those available for *A. fumigatus*. We therefore deployed a hybrid sampling strategy that leveraged both short- and long-read genome assemblies, combined with short-read mapping to genotype *Starship* presence/absence, to enable an accurate and precise accounting of how *Starships* impact strain heterogeneity. We downloaded 508 publicly available *A. fumigatus* assemblies (mostly assembled from short-read data) that span the known genetic diversity of this species isolated from environmental and clinical sources (51.7% clinical, 47.9% environmental, 0.4% unknown isolation source) and used Oxford Nanopore to sequence 11 additional strains with long-read technology (66.7% clinical, 25% environmental, 8.3% unknown), for a combined total of 519 assemblies (10, 11). The 11 isolates selected for long-read sequencing were chosen because they represent commonly utilized laboratory strains, clinical isolates and environmental strains from three major *A. fumigatus* clades (10). We performed additional short-read Illumina sequencing of the 11 strains to provide additional support and error-correct the long-read assemblies. The long-read assemblies are of reference quality, representing nearly full chromosome assemblies with an L50 range of 4-5 and a N50 range of 1.9-4.7 Mb, and are thus ideal complements to the available short-read assemblies for investigating the connection between *Starship*s and strain heterogeneity (Table S1).

### At least 20 active and distinct *Starships* vary among *A. fumigatus* strains

We began evaluating the impact of *Starships* on strain heterogeneity by systematically annotating them in the 519 assemblies using the starfish workflow (22). We identified a total of 787 individual *Starships* and validated these predictions by manually annotating a subset of 86 elements (Table S2, Table S3, Methods) (10, 11). As expected, more *Starships* were recovered in total from short-read assemblies, but on average more *Starships* were recovered from each reference-quality long-read assembly, highlighting the utility of sampling both short- and long-read assemblies (Fig. S1). Importantly, unlike many other fungal transposons, we found that with few exceptions, if a *Starship* is present in a strain, it is present at most in a single copy. Some other species appear to readily accumulate multiple copies of the same *Starship* (20), but this is not the case for *A. fumigatus*. In other fungi, genome defense mechanisms such as repeat-induced point mutations (RIP) are capable of degrading multi-copy *Starships* and may act to limit their copy number expansion (21); however, it remains unclear whether such mechanisms also explain the lack of copy number variation observed in *A. fumigatus*.

To determine how many different *Starships* actively generate variability in *A. fumigatus*, we grouped *Starships* into discrete types following a hierarchical framework that incorporates amino acid sequence comparisons of captain tyrosine recombinases and nucleotide sequence comparisons of the entire element (23). Following Gluck-Thaler and Vogan 2024, we assigned each *Starship* element to a family (based on similarity to a reference library of captain tyrosine recombinases), a *navis* (Latin for “ship”; based on orthology among the *A. fumigatus* captains) and a *haplotype* (based on k-mer similarity scores of the cargo sequences; Methods). For example, *Starship Osiris h4* belongs to *haplotype* 4 within the *Osiris navis*, which is part of the Enterprise-family. The most reasonable conclusion when observing the same transposon bounded by identical direct repeats at two different loci in two individuals is that this transposon is presently or recently active (21). Using a threshold that required observing the same *navis-haplotype* (hereafter *Starship*) at two or more sites, we identified 20 high-confidence *Starships* that met these criteria and 34 medium-confidence *Starships* that did not. These data reveal a phylogenetically and compositionally diverse set of *Starships* actively transposing within *A. fumigatus* that together make up this species’ *Starship* compartment (Methods, Table 1, Fig. 1, Table S4, Table S5) (23). For each of the 20 high-confidence elements, we aligned representative copies present in different genomic regions to highlight how precisely and repeatedly element boundaries are conserved across transposition events (Fig. S2).

**Figure 1:**
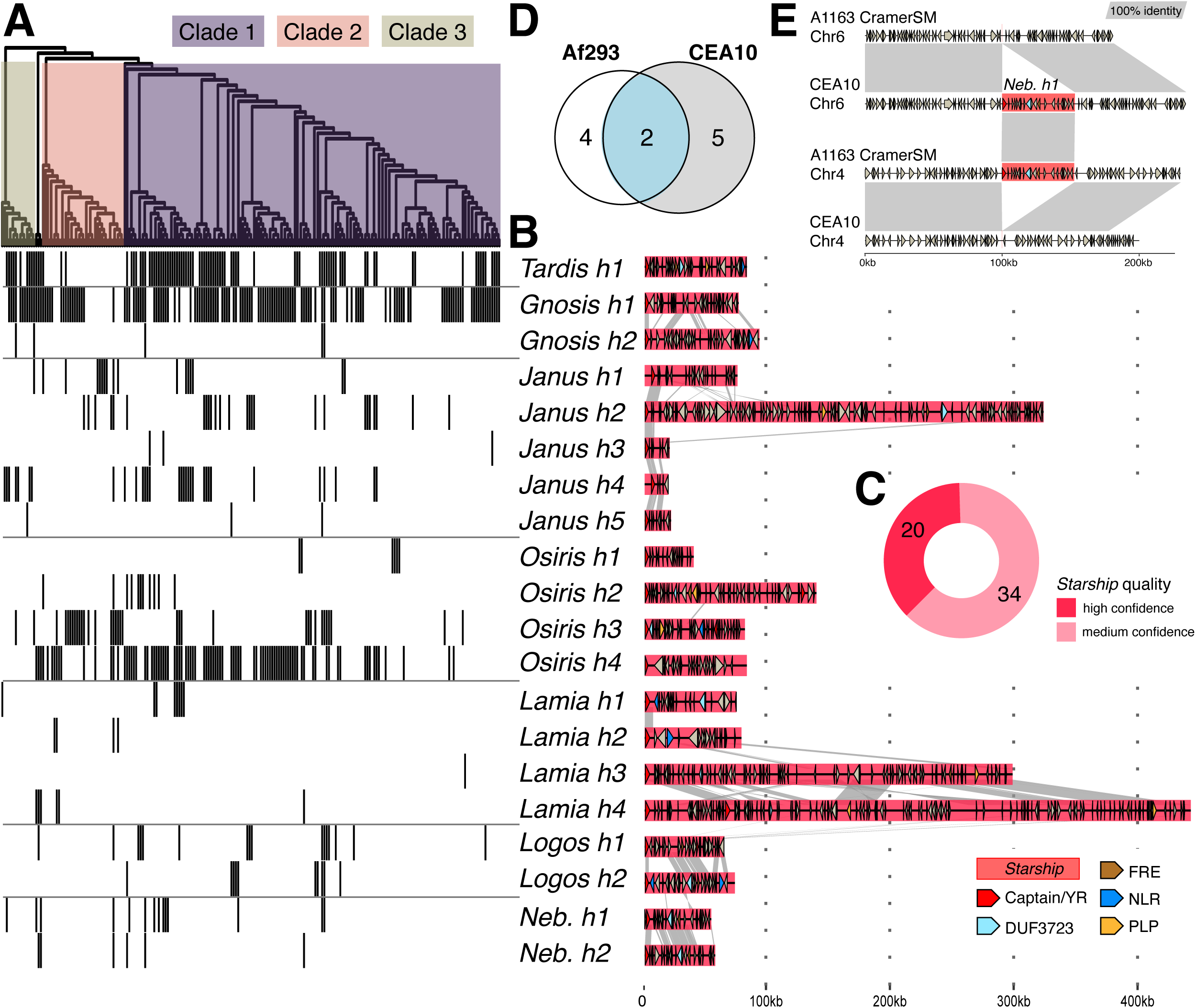
At least 20 distinct *Starships* carrying hundreds of protein-coding genes vary in their presence/ absence across *Aspergillus fumigatus* strains. A) Top: a SNP-based maximum likelihood tree of 220 *A. fumigatus* strains from Lofgren et al. 2022 (visualized as a cladogram for clarity), color-coded according to phylogenetic clade as defined by Lofgren et al. 2022. Bottom: A heatmap depicting the presence (gray) and absence (white) of 20 high-confidence *Starships* in each isolate. B) Schematics and sequence alignments of the type elements from 20 high-confidence *Starships*, where links between schematics represent alignable regions ≥500bp and ≥80% nucleotide sequence identity and arrows represent predicted coding sequences. C) A donut chart summarizing the number of distinct types of *Starships* by the quality of their prediction. D) A Venn diagram indicating the number of shared and unique high-confidence *Starships* in the reference strains Af293 and CEA10 (Table S7). E) Pairwise genome alignments between the CEA10 isolate from BioSample SAMN28487501 and the CEA10-derived A1163 isolate sequenced in this study demonstrating *Starship* movement. *Nebuchadnezzar h1* is present on Chromosome 6 in CEA10 (CP097568.1: 3645813-3698867) and absent at the corresponding locus in A1163 (NODE10: 188873-188877). Conversely, *Nebuchadnezzar h1* is present on Chromosome 4 in A1163 (NODE6: 77737-130777) but absent from the corresponding locus in CEA10 (CP097566.1:1344957-1344961).

After using all available genome assembly data to define the *Starship* compartment with automated methods, we explored a variety of other approaches for augmenting and validating our predictions, including manual annotation, BLAST-based detection, and short-read mapping (Methods). Certain detection methods were more appropriate for particular analyses, resulting in the partitioning of our findings into the high-confidence, the expanded, and the genotyping datasets. To analyze *Starship* features, like gene content and pangenome distributions (see below), we limited our analyses to the set of 459 high-confidence elements for which we have end-to-end boundaries and evidence of transposition (high-confidence dataset). For analyses that examine the presence/absence of elements across genomic regions, we used an expanded set of 1818 elements that consists of the 787 high and medium-confidence elements as well as 1031 elements whose presence was detected through BLAST searches, because these methods maximize our ability to correctly predict the presence/absence of an element in a given region (expanded dataset; Table S4, Table S5). For genotyping analyses that focus on detecting the presence/absence of an element regardless of where it is found in the genome, we took a conservative approach by limiting our analysis to 13 reference quality assemblies (reference strains CEA10 and Af293 in addition to the 11 newly sequenced long-read strains) plus 466 *A. fumigatus* strains with publicly available paired-end short-read Illumina data that were used to validate *Starship* presence/absence by short-read mapping to a reference *Starship* library (genotyping dataset; Table S21).

### *Starships* distributions are partially explained by strain relatedness but overall heterogeneous

All high-confidence *Starships* show polymorphic presence/absence variation, representing a previously unexamined source of genetic heterogeneity across *A. fumigatus* strains (Fig. 1A, Table S6). For example, the two commonly used reference strains Af293 and CEA10 carry 11 different high-confidence *Starships* but share only 4 in common (or only 2, using a more conservative threshold of not counting fragments or degraded elements; Fig. 1D, Table S7). The number of high-confidence *Starships* per long-read isolate ranges from 0-11 (median = 3, std. dev. = 3.39) (29, 30). To identify the underlying drivers of variation in *Starship* distributions, we investigated the relationship between *Starship* repertoires and strain relatedness by testing if phylogenetic signal underlies *Starship* distributions. We genotyped *Starship* presence/absence using short-read mapping in the best sampled clade of *A. fumigatus* with well resolved phylogenetic relationships (Clade 1 *sensu* Lofgren et al., 2022, n = 151) and found that isolate relatedness is weakly but significantly correlated with similarity in *Starship* repertoire, which is indicative of weak phylogenetic signal in *Starship* distributions (Fig. S3, Table S8, Mantel r = 0.163; *P* = 0.001, using the genotyping dataset). The relatively weak but significant correlation indicates that although closely related isolates tend to share similar *Starships*, relatedness alone cannot explain all observed variation in *Starship* repertoires. Thus, we conclude *Starships* are a source of genetic heterogeneity among even closely related strains.

While no sequence-based or phylogenetic method to our knowledge would enable us to unequivocally differentiate horizontal transfer events from incomplete lineage sorting, loss or sexual and parasexual recombination at the within species level, we hypothesize that horizontal transfer within *A. fumigatus* is at least partially responsible for generating the observed patchy distributions of *Starships*. *Starship* horizontal transfer has occurred repeatedly between individuals of different genera, and barriers to transfer are likely lower within species (25, 26). *Starship* horizontal transfer has recently been experimentally demonstrated under simple co-culture conditions between *A. fumigatus* and *Paecilomyces variotii*, which are separated by approximately ∼100 million years of evolutionary divergence (28). Surveys of the *Paecilomyces* fungi further estimate that ∼1/3 of *Starships* in this genus have evidence of horizontal transfer in natural populations (28). The dynamics of *Starship* horizontal transfer among *A. fumigatus* strains must be determined with future laboratory experiments.

### *Starships* actively transpose in isolates of the same laboratory strain

We examined the potential for *Starships* to introduce unwanted variation into laboratory experiments by comparing isolates of the same reference strain used by different research groups. First, we found that *Starship* insertions in the reference Af293 assembly used in our study coincide with 4/8 putative structural variants inferred in other Af293 strains using RNAseq expression data by Collabardini et al. 2022 (Table S9). This suggests that in addition to other mechanisms, *Starship* transposition and/or loss contributes to structural variation observed among isolates of what should otherwise be clones of the same strain (6).

We gathered additional evidence supporting our hypothesis by comparing independently collected DNA sequencing data from different isolates of the same or derived strains. We compared the locations of *Starships* between the long-read CEA10=CBS144.89 reference assembly (31) and 6 other CEA10 and CEA10-derived isolates from various labs, including our own, using short-read mapping and de-novo assembly comparisons (Methods). We found that *Starship Nebuchadnezzar h1* has jumped from chromosome 6 in the long-read reference CEA10 and A1160 assemblies and some A1163 strains to chromosome 4 in the A1163 isolate used by our lab (A1163-Cramer_SM), likely at some point after we acquired this isolate from its original culture collection stock (Fig. 1E). A1163 is a commonly used laboratory strain derived from CEA10; the only differences between A1163 compared with CEA10 should be that its native *pyrG* has a nonsense mutation and that it harbors an ectopic insertion of the *A. niger pyrG* (32). Similarly, we evaluated *Starship* heterogeneity in three additional strains, S02-30 (=AF100-9B), TP9 (=08-19-02-30) and ATCC46645, that met the criteria of having independently collected DNA sequencing datasets and long read assemblies. We found that ATCC46645 showed signs of *Starship* heterogeneity among isolates sequenced by different research groups. Specifically, a small number of reads supported precise excision events for *Nebuchadnezzar h1*, suggesting this *Starship* has transposed or has been lost in a subset of the nuclei within the short-read sequenced isolate (Fig. S4). This suggests that some *Starships* appear to be active under laboratory conditions, which is directly relevant for experimental design in this case as *Nebuchadnezzar h1* carries the HAC gene cluster known to impact biofilm-associated virulence (see below) and genes on *Starships* may be lost through failed transposition events (21, 33).

Together, *Starship*-mediated variation among isolates of the same “clonal” *A. fumigatus* strain suggests these transposons cause genomic instability over short-enough timescales to potentially impact routine laboratory work and experimental reproducibility. It is interesting to note that not all strains or closely related strains demonstrate intraspecific variation: for example, although we did not detect variation in *Starship* location or presence/absence between the long-read CEA10 and A1160 assemblies, which have been separated for decades (31), we did detect variation between this CEA10 strain and the A1163 isolate that we routinely use in our lab, indicating that much remains to be known about the signals and triggers promoting within-strain *Starship* variation.

### *Starships* mobilize upwards of 16% of the accessory genome that differs across strains

*A. fumigatus* strains differ extensively in the combinations of genes found in their genomes, resulting in a large and diverse pangenome (10, 11). To quantify how much pangenomic variation is attributable to actively transposing *Starships,* we estimated the total proportion of genomic content present in the 20 high-confidence *Starships* (Fig. 2, Methods). We examined these *Starships* in the 13 reference-quality assemblies and found that between 0-2.4% of genomic nucleotide sequence (and 0-2.1% of all genes per genome) is mobilized as *Starship* cargo (Fig. 2A). We then built a pangenome with all 519 strains and extracted all orthogroups (i.e., genes) found in the 13 reference-quality genomes to gain insight into the distribution of *Starship*-associated gene content (Table S10). Across the 13 isolate pangenome, 2.9% of all genes, which corresponds to 9.7% of all accessory and singleton orthogroups, have at least 1 member carried as *Starship* cargo (Fig. 2B). Accessory and singleton orthogroups are overrepresented >3 fold in *Starships*, with ∼92% of *Starship*-associated orthogroups being either accessory or singleton compared with 24.6% of non-*Starship* associated orthogroups. We determined the upper bounds of this conservative estimate by examining the 13 isolate pangenome in the context of all 54 high- and medium-confidence *Starships*, and found that 4.8% of all orthogroups, representing 16% of all accessory and singleton orthogroups, have at least 1 member carried as *Starship* cargo (Fig. S5, Table S11). Drawing parallels from decades of observations of bacterial MGEs (17) and recently published experimental data from *Starships* (21, 28), we hypothesize that localization on a *Starship* promotes a gene’s rate of gain and loss through *Starship-*mediated horizontal transfer and failed re-integration events (21). Together, these data reveal a previously hidden association between *Starships* and the making of *A. fumigatus’* accessory pangenome.

**Figure 2:**
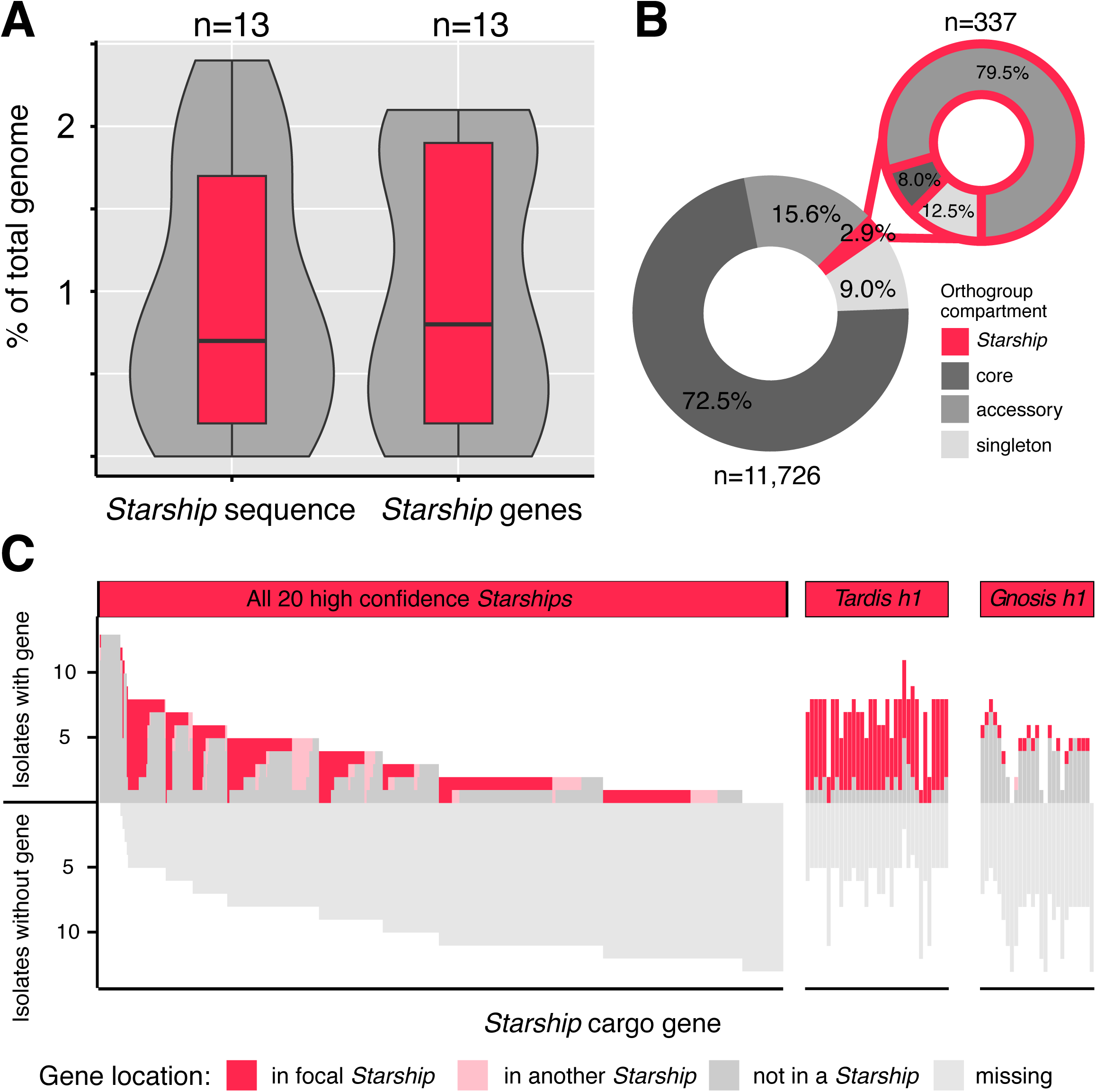
*Starships* are enriched in accessory genes whose presence/absence and genomic location vary across *Aspergillus fumigatus* strains. All panels visualize data derived from 13 reference-quality *A. fumigatus* assemblies and the 20 high-confidence *Starships* (n = 459 elements total). A) Box-and-whisker plots summarizing the total percentage of nucleotide sequence and predicted genes carried by *Starships* per genome. B) Donut charts summarizing the percentages of gene orthogroups in the core, accessory, singleton, and *Starship*-associated compartments of the *A. fumigatus* pangenome (Table S10). C) Iceberg plots summarizing the genomic locations of the single best BLASTp hits (≥90% identity, ≥33% query coverage) to the cargo genes from the type elements of all 20 high-confidence *Starships* (left) and two individual *Starships* (right; Table S12). Each column in the iceberg plot represents a cargo gene and is color-coded according to the genomic location of hits (full dataset in Fig. S5).

### *Starships* differ in their potential to mediate gain and loss of unique sequence

We further determined the potential for *Starships* to introduce variation into *A. fumigatus* strains by testing if *Starship* genes are uniquely found in those elements or found elsewhere in the genome. We calculated the degree of association between a given cargo gene and a *Starship* by identifying the genomic locations of best reciprocal BLAST hits to genes in the 20 high-confidence type elements (Fig. 2C, Fig. S6, Table S12, Table S13, Methods). Across the 13 reference-quality assemblies, we found that the vast majority of *Starship* genes display presence/absence variation between strains (i.e., are accessory or singleton) although several genes are present in conserved regions in these particular strains (while seemingly counter-intuitive, these genes do vary in their presence/absence and are *Starship-*associated when examining the larger 519 strain population, as expected). We found that many genes are almost always carried as cargo in active *Starships* when present in a given genome (e.g., the majority of cargo on *Tardis h1*), while others have weaker associations and are found in both *Starships* and non-*Starship* regions across different strains (e.g., *Gnosis h1*). Variation in the degree to which genes associate with active *Starships* suggests elements differ in capacity to generate accessory sequence variation among strains, highlighting the importance of investigating *Starships* at the individual element level.

### *Starship* activity generates structural variation at a genome-wide scale

We next asked where *Starship*-mediated variation occurs in the genome to better understand the implications of *Starship* activity for genome organization. We identified the genomic locations where *Starships* introduce structural variation by sorting all 787 high and medium-confidence elements, along with elements detected by BLASTn, into homologous genomic regions (Table S14, Table S15, Table S16; expanded dataset; Methods). This enabled us to genotype individuals in the 519 strain population for segregating *Starship* insertions that are polymorphic across strains (Fig. 3, Fig. S7; Methods). Across all strains and chromosomes, we found 79 regions that contain at least 1 segregating “empty” insertion site, for a total of 154 sites distributed across all eight chromosomes (a single region can have >1 insertion site if they are located close to each other). The average number of empty sites per genome is 44.5 (range = 9-63, std. dev. = 13.88), indicating that each strain harbors dozens of sites with structural variation introduced by *Starships*. For example, the Af293 reference strain has a total of 6 full-length *Starships* with annotated boundaries and 2 *Starship* fragments, and a total of 56 segregating empty sites (Fig. 3A). We found a significant linear relationship between the number of genomic regions a given *Starship* is present in (a proxy for transposition activity) and the total copy number of that element in the 519 strain population, suggesting active *Starships* contribute more to strain heterogeneity compared with less active elements (Fig. 3C; y = 14.7x - 1.24; *P =* 6.5e^-4^; R^2^ adj = 0.46; expanded dataset). Predicted *Starship* boundaries are precisely conserved across copies present in different genomic regions, suggesting transposition, and not translocation or segmental duplication, is the major mechanism responsible for generating variation in *Starship* location (Fig. 3D).

**Figure 3:**
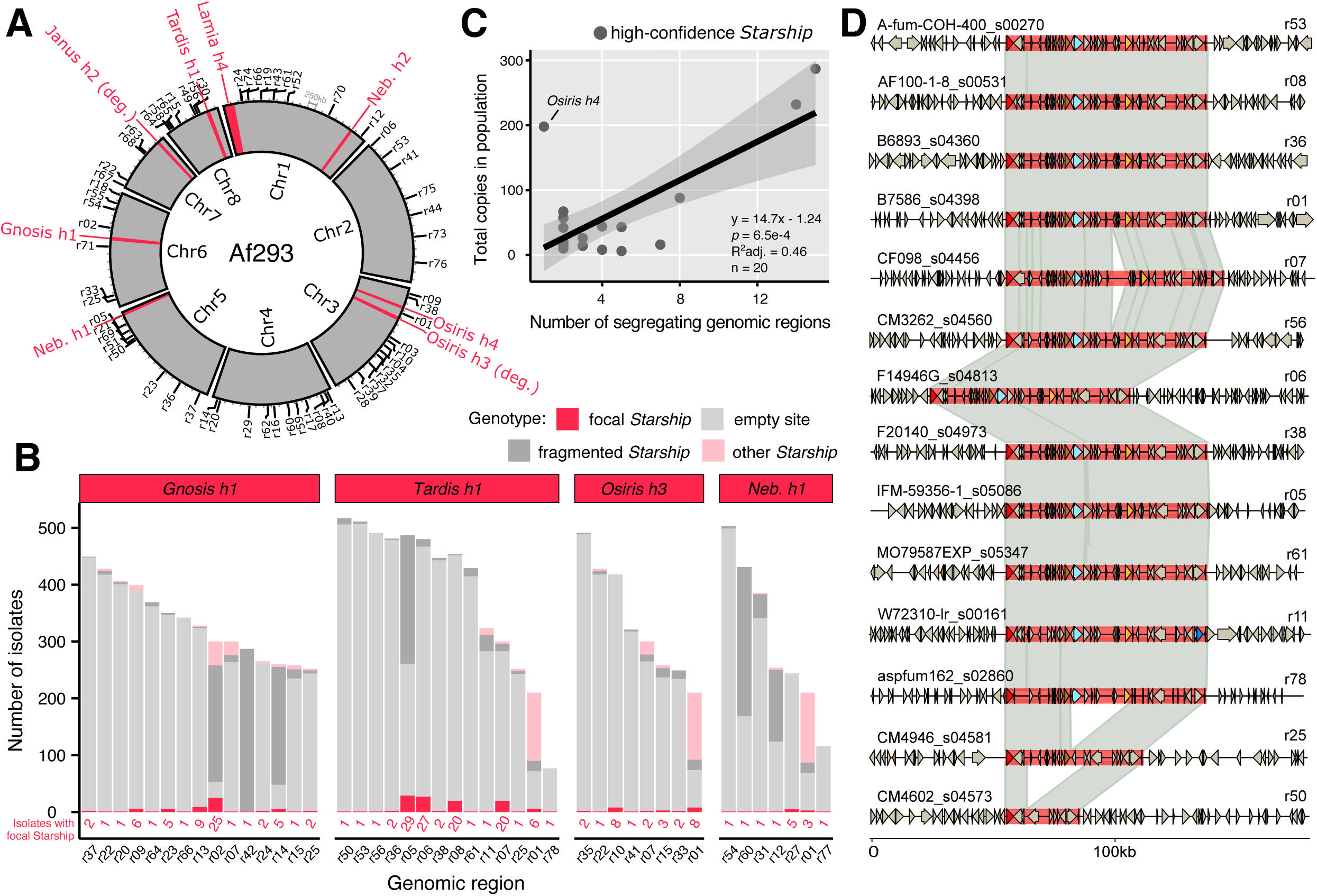
*Starships* and their insertion sites are distributed across all major chromosomes in the *Aspergillus fumigatus* genome. A) A Circos plot summarizing all inserted *Starships* (in red) and all genomic regions containing either an empty insertion site or a fragmented *Starship* (in black, along perimeter and labeled with the r prefix) in the 8 chromosomes of the *A. fumigatus* reference strain Af293 (Table S14). All genomic regions contain a *Starship* insertion in some other individual from the 519 strain population. B) Barcharts summarizing the genotypes of segregating genomic regions associated with the four most active high-confidence elements in the 519 strain population (Table S16; full dataset in Fig. S7). If an isolate did not have any *Starships* within a given region, it was assigned either an “empty” or “fragmented” genotype (Methods). C) A scatterplot summarizing the relationship between the number of genomic regions containing a given *Starship* and the total number of copies of that *Starship* in the 519 strain population, where each point represents one of the 20 high-confidence *Starships*. A line derived from a linear regression model is superimposed, with shaded 95% confidence intervals drawn in gray. D) Alignments of *Tardis h1* copies +/- 50kb of flanking sequence across 14 genomic regions. Links between schematics represent alignable regions ≥1000bp and ≥90% nucleotide sequence identity.

Given that *Starships* are mobilized by a tyrosine site-specific recombinase (21), we attempted to predict each *Starship*’s target site to gain insight into the genomic sequences that are susceptible to *Starship* insertions. We manually curated the sequence features of a large sample of high-confidence *Starships* and found 19/20 have identifiable direct repeats (DRs) ranging from 2-12 bp in length while 18/20 have terminal inverted repeats (TIRs; Table 1, Table S3). DRs typically reflect a portion of the element’s target site, while TIRs are predicted to facilitate transposition (34). While precise target site motifs must be confirmed experimentally, the target site TTACA(N_5_)AAT that we recovered for *Nebuchadnezzar* elements, which belong to the Hephaestus*-*family, resembles the canonical *Hephaestus* target site TTAC(N_7_)A, demonstrating the utility of our approach as a first step towards understanding what sequence features promote the gain and loss of *Starship*-mediated variation (21).

Generally, DRs >6 bp are associated with targeting of the 5S rDNA gene (e.g. *Osiris*), presumably because this is a relatively stable target site (22). In contrast to this, *Tardis* has a highly conserved 9 - 11 bp DR that does not correspond to the 5S gene. A k-mer analysis shows that the 10 bp consensus motif CTACGGAGTA is strongly overrepresented in the genome of Af293 (>99.99th percentile of k-mers; Fig. S8), indicating that this motif may represent some other type of highly conserved genomic sequence, such as a transcription factor binding site. This motif overlaps with predicted motifs associated with C6 zinc cluster factors in *A. nidulans* (https://jaspar.elixir.no/matrix/UN0291.2/) (35) and this DR sequence is further conserved among other *Eurotiomycetes* (22); however, the exact functions of these predicted motifs must be confirmed experimentally. This result highlights the fact that studying *Starship* elements in detail can reveal other important aspects of a species’ biology, and characterizing the *Tardis* motif is of interest for future research.

### *Starships* mediate variation at an idiomorphic biosynthetic gene cluster

Several genomic regions harbor multiple types of *Starships*, raising the possibility that strain heterogeneity arises in part from the formation of *Starship* insertion hotspots. We investigated the genomic region with the highest density of *Starship* insertions to define the upper bounds of *Starship*-mediated structural variation at a single locus (Fig. 4C, Methods). This region spans an average of 498.76 Kb (range = 158.07-781.54 kb; std. dev. = 83.02 kb) across the n = 210 strains for which we could detect it, and ranges from position 584,521-1,108,409 on Chromosome 3 in the Af293 reference assembly (representing 12.84% of the entire chromosome). By supplementing starfish’s automated genotyping with manual annotation of nine strains with distinct alleles at this region, we found this region contains at least 7 distinct *Starships* with identifiable DRs and 1 degraded *Starship* that range from 47.52-151.62 kb long and are inserted into 6 independent segregating sites (Fig. 4C). The majority (75%) of the *Starships* in this region are inserted into 5S rDNA coding sequences, which effectively fragments that copy of the 5S rDNA gene. Total 5S rDNA copy number varies between 28-33 in the 13 reference quality assemblies (median = 31, std. dev. = 1.4; Table S17), but between 6-8 intact copies are typically present at this single locus, representing a ∼10-fold enrichment of the 5S rDNA sequence relative to background expectations given the length of this region (using 5S rDNA frequencies in the Af293 genome). Thus, the enrichment of 5S rDNA at this locus potentiates *Starship*-mediated variation and the generation of a *Starship* hotspot.

**Figure 4:**
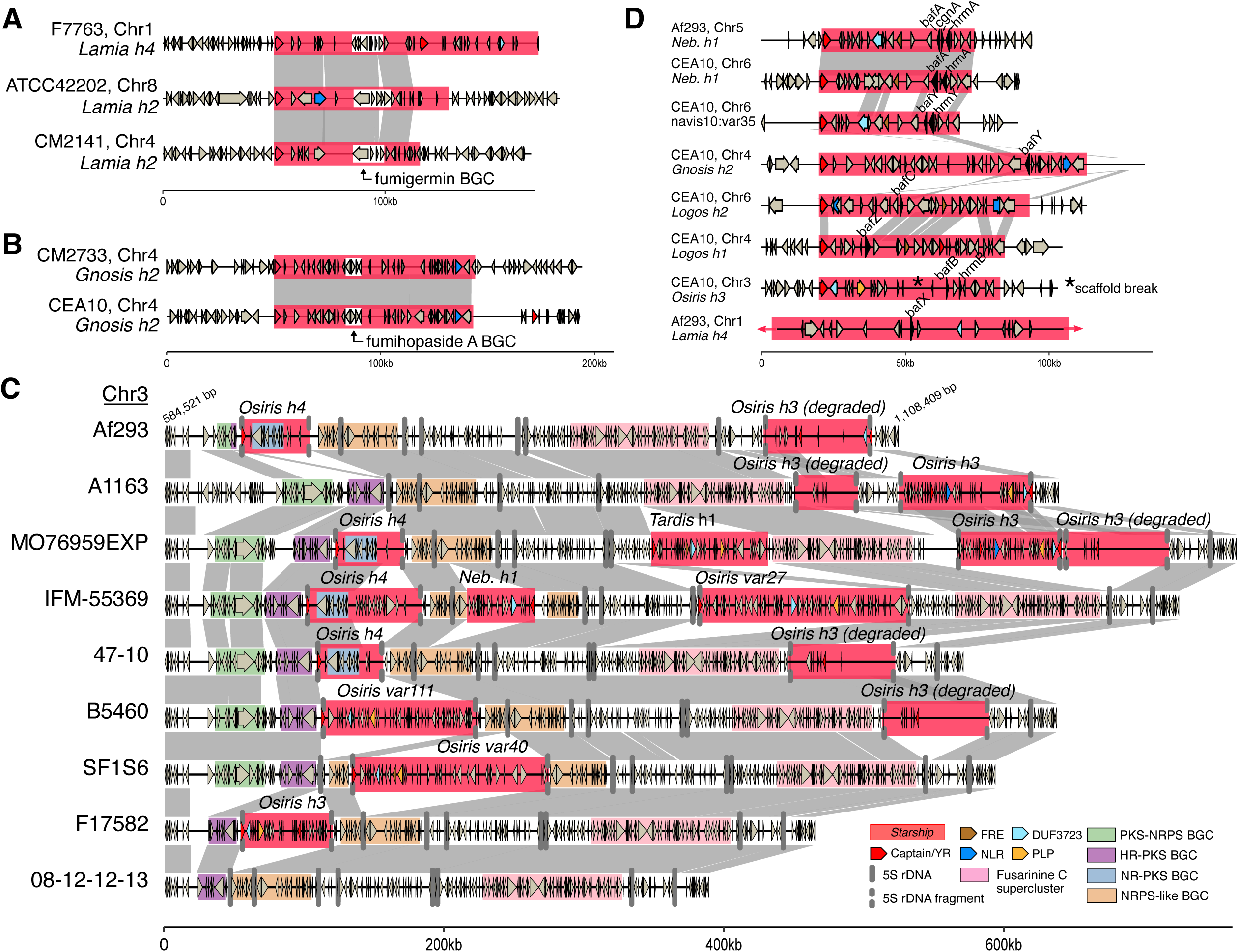
*Starships* mobilize adaptive traits and generate allelic diversity among *Aspergillus fumigatus* strains. Schematics and alignments of: A) *Starship Lamia h2* and *h4*, which carry the biosynthetic gene cluster (BGC) encoding the polyketide secondary metabolite fumigermin, shown inserted at 3 independent sites. B) Starship *Gnosis h2* carrying the BGC encoding the terpene secondary metabolite fumihopaside A, shown inserted at 2 independent sites. C) a large region on Chromosome 3 previously identified as an idiomorphic BGC containing multiple segregating *Starship* insertions and various combinations of putative BGCs, including a a non-reducing polyketide synthase (NR-PKS) BGC carried by *Starship Osiris h4*. *Starships* from the *Osiris* navis specifically insert in 5S rDNA sequence and are predicted to fragment it. D) Eight *Starships* in the reference strains Af293 and CEA10 that all carry homologs of biofilm architecture factor A (bafA). *Starship Nebuchadnezzar h1* (*Neb. h1*) carries bafA as part of the H_A_AC (hrmA-associated gene cluster), while *Starship navis10-var35* carries bafB as part of H_B_AC and *Starship Osiris h3* carries bafC as part of H_C_AC. Only a portion of *Starship Lamia h4* is visualized for figure legibility. All data for A-D was collected from the 519 *A. fumigatus* strain population (Table S5). Links between schematics represent alignable regions ≥5000bp and ≥95% nucleotide sequence identity and arrows represent predicted coding sequences. Abbreviations: polyketide synthase (PKS); non-ribosomal peptide synthetase (NRPS); highly reducing (HR).

We were surprised to find that in addition to multiple segregating *Starships*, the Chromosome 3 region contains an idiomorphic biosynthetic gene cluster (BGC) that was previously noted to be a recombination hotspot (Cluster 10 *sensu* Lind et al. 2017) (36, 37). The idiomorphic BGC locus is polymorphic for upwards of 4 smaller BGC modules and is upstream of a fifth BGC encoding the biosynthesis of fusarinine C (BGC8 *sensu* Bignell et al. 2016) that itself is embedded within a larger stretch of sequence containing NRPS and KR-PKS core genes. While the idiomorphic modules were not predicted by antiSMASH, manual inspection revealed that each contains a different core SM biosynthesis gene and numerous other genes involved in metabolic processes, suggesting they are part of a larger BGC or represent different cryptic BGCs. The NR-PKS BGC module at this locus is carried by *Starship Osiris h4,* and its presence/absence is directly associated with the presence/absence of the element, implicating *Starships* as a mechanism generating idiomorphic BGCs. Furthermore, some BGC modules are disrupted by the insertion of a *Starship* (e.g., the NRPS-like BGC module). Together, these findings support the assertion that *Starship* activity helps generate selectable variation in BGC genotypes. The implication of this variation for the expression and diversification of natural products warrants further investigation.

### *Starships* encode clinically-relevant phenotypes and are enriched in clinical strains

*Starships* encode genes with diverse functions typically associated with fungal fitness, including carbohydrate active enzymes, BGCs and metal detoxification genes (Table 1, Table S5). To investigate broad trends in how *Starships* might contribute to phenotypic heterogeneity, we compared the predicted cargo functions among the 20 high-confidence *Starships* (Table S18; high-confidence dataset; Methods). As expected, *Starship* types differ from each other in their predicted functional content, but functional diversity also varies to some extent within elements from the same type, indicating that both inter- and intra-specific *Starship* variation may contribute to functional heterogeneity among *A. fumigatus* strains. In this sense, different copies of the “same” *Starship* in different individuals have much greater potential to differ from each other at the sequence level compared with much smaller transposons. We found that 49.7% of COG-annotated genes in the high-confidence *Starships* are “Poorly Categorized”; 28.8% belong to the “Metabolism” category, 14.7% belong to “Cellular Processes and Signaling” and 6.8% belong to “Information Storage and Processing”.

Given the importance of metabolic processes for diverse fungal traits, we examined granular classifications within the Metabolism category and found that *Starship* types differ specifically in their contributions to metabolic heterogeneity (Fig. S9). For example, nearly all 75 copies of *Gnosis h1* carry at least 1 gene with a predicted role in “Coenzyme transport and metabolism,” and none carry genes with predicted roles in “Lipid transport and metabolism”, while the exact opposite is true for the 119 copies of *Tardis h1*. The most frequently mobilized COG metabolism categories are “Carbohydrate transport and metabolism” (present in 371 individual elements belonging to 13 high-confidence *Starships*), “Secondary metabolite transport and metabolism” (present in 329 elements belonging to 16 high-confidence *Starship* types) and “Energy production and conversion” (present in 281 elements belonging to 9 high-confidence *Starship* types), each of which has known associations with clinically-relevant pathogen phenotypes. We found no evidence that any metabolism-associated COG category was enriched in *Starships* compared with background frequencies in Af293, indicating it is likely the specific mobilized functions, as opposed to an enrichment of certain classes of functions, that defines *Starship* contributions to fungal phenotypes (data not shown).

While investigating associations between *Starships* and characterized pathogen phenotypes, we found three notable examples of cargo genes that encode known traits important for pathogen survival and virulence (Table S19, Methods) (10, 11, 36, 38–44). The BGC encoding fumihopaside A (AFUA_5G00100-AFUA_5G00135) is carried by *Starship Gnosis h2* (Fig. 4A). fumihopaside A is a triterpenoid glycoside that increases fungal spore survival under heat and UV stress exposure (38). Similarly, the BGC encoding the polyketide fumigermin (AFUA_1G00970-AFUA_1G01010), which inhibits bacterial spore germination, is carried by *Lamia h2* and *h4* (Fig. 4B) (45). Both the fumihopaside A and fumigermin BGCs were previously identified as “mobile gene clusters” based on their presence/absence at different chromosomal locations in *A. fumigatus* strains, and our results reveal that *Starships* are the mechanism underpinning their mobility (Cluster 1 and 33, respectively from Lind et al., 2017) (36).

Finally, a cluster of three genes (AFUA_5G14900/*hrmA*, AFUA_5G14910/*cgnA*, AFUA_5G14915/*bafA*, collectively referred to as the *<u>h</u>rmA*-<u>a</u>ssociated <u>c</u>luster or HAC) carried by *Nebuchadnezzar h1* increases virulence and low oxygen growth, and its expression modulates colony level morphological changes associated with biofilm development (Fig. 4D) (33, 46). The genes *bafB* and *bafC*, which are homologs of *bafA,* also mediate colony and submerged biofilm morphology and are each carried by up to 5 additional *Starships*, indicating a sustained association between this gene family and *Starships* (Fig. 4D, Fig. S10) (33). Mobilization of biosynthetic gene clusters and biofilm-related loci by *Starships* has potential to contribute to the rapid evolution of these traits. For example, *Starships* are known to mediate rapid gene gain and loss through horizontal *Starship* transfer, which has recently been experimentally demonstrated using *A. fumigatus* as a recipient (28), and *Starship* loss, which has been experimentally demonstrated to occur through failed re-integration events (21). Thus, it is reasonable to predict that *Starship*-mediated gain and loss effectively increases the capacity of BGC and biofilm-associated traits to rapidly increase or decrease in frequency within populations; however, this hypothesis remains to be tested experimentally. Minimally, we propose that by mobilizing genes contributing to biofilm formation, *Starships* help drive heterogeneity in clinically-relevant phenotypes at a population level (33).

To identify *Starships* that could be relevant in either environmental or clinical settings, we tested for an association between the high-confidence *Starships* and strain isolation source (genotyping dataset; Methods). We found that 5 high-confidence *Starships* are significantly enriched (*P*_adj_ <0.05) in strains from either clinical or environmental isolation sources across the 475/479 genotyped strains for which source data exists (Table S2, Table S20, Table S21, Fig. S11). *Lamia h3* (*P*_adj_ = 0.013) and *Janus h3* (*P*_adj_ = 0.034) are enriched in environmental strains. *Osiris h4* (*P*_adj_ = 0.022), *Nebuchadnezzar h1* (*P*_adj_ = 0.015; carries *bafA*, see above) and J*anus h2* (*P*_adj_ = 0.013) are enriched in clinical strains, providing complementary evidence to recently published analyses of *Starship* captain enrichment in clinical isolates across several other fungal species (47). Although clinical and environmental strains were sampled in roughly equal proportions, we can’t completely rule out biases in our sampling (e.g. phylogenetic and ecological biases) that would contribute to these enrichment patterns. Nevertheless, for each of the 5 enriched *Starships*, strains were often isolated from different countries (between 3-10 countries) and often belong to different phylogenetic groups (between 1-3 clades *sensu* Lofgren et al., 2022 and 1-6 phylogenetic clusters *sensu* Barber et al., 2021), suggesting these *Starships* are excellent candidates to experimentally test how giant transposons contribute to fitness in environmental and clinical settings.

### *Starship* expression contributes to heterogeneity in a strain-, treatment- and strain by treatment-dependent manner

The annotation of *A. fumigatus’ Starship* compartment next allowed us to test the hypothesis that *Starship* cargo is expressed under clinically-relevant conditions. We analyzed patterns of differential *Starship* cargo gene expression in 14 publicly available transcriptomic studies from three commonly used reference strains (Af293, CEA10, A1163). In total, 177 transcriptome samples were included in this meta-analysis. Samples were split into broad treatment categories and corrected for batch effects to allow for comparison of cargo gene expression between studies (Methods). Out of 596 *Starship* genes total (carried by the high-confidence *Starships* present in the three reference strains) and across all treatments, we identified 459 differentially expressed genes (DEGs) that included both captain tyrosine recombinases and cargo (Supplementary Data).

To identify experimental conditions that may impact *Starship* transposition and the subsequent generation of *Starship*-mediated heterogeneity, we first examined patterns of transcript abundance for the captain genes of the 12 high-confidence *Starships* in Af293, CEA10 and A1163 (Fig. 5A, Fig. S12). All captain genes have evidence for constitutive gene expression (a minimum median transcript coverage of 1 across the gene body for biological replicates) in at least one experimental condition (Fig. S13). Overall, 8 of the 12 captain genes from high-confidence *Starships* are differentially expressed in at least one study for one or more reference strains, revealing many opportunities for *Starship* transposition under lab conditions (Supplemental information). Differences in experimental conditions and/or strain backgrounds have no consistent positive or negative influence on captain gene expression, indicating that the intricacies of captain expression remain to be elucidated.

**Figure 5:**
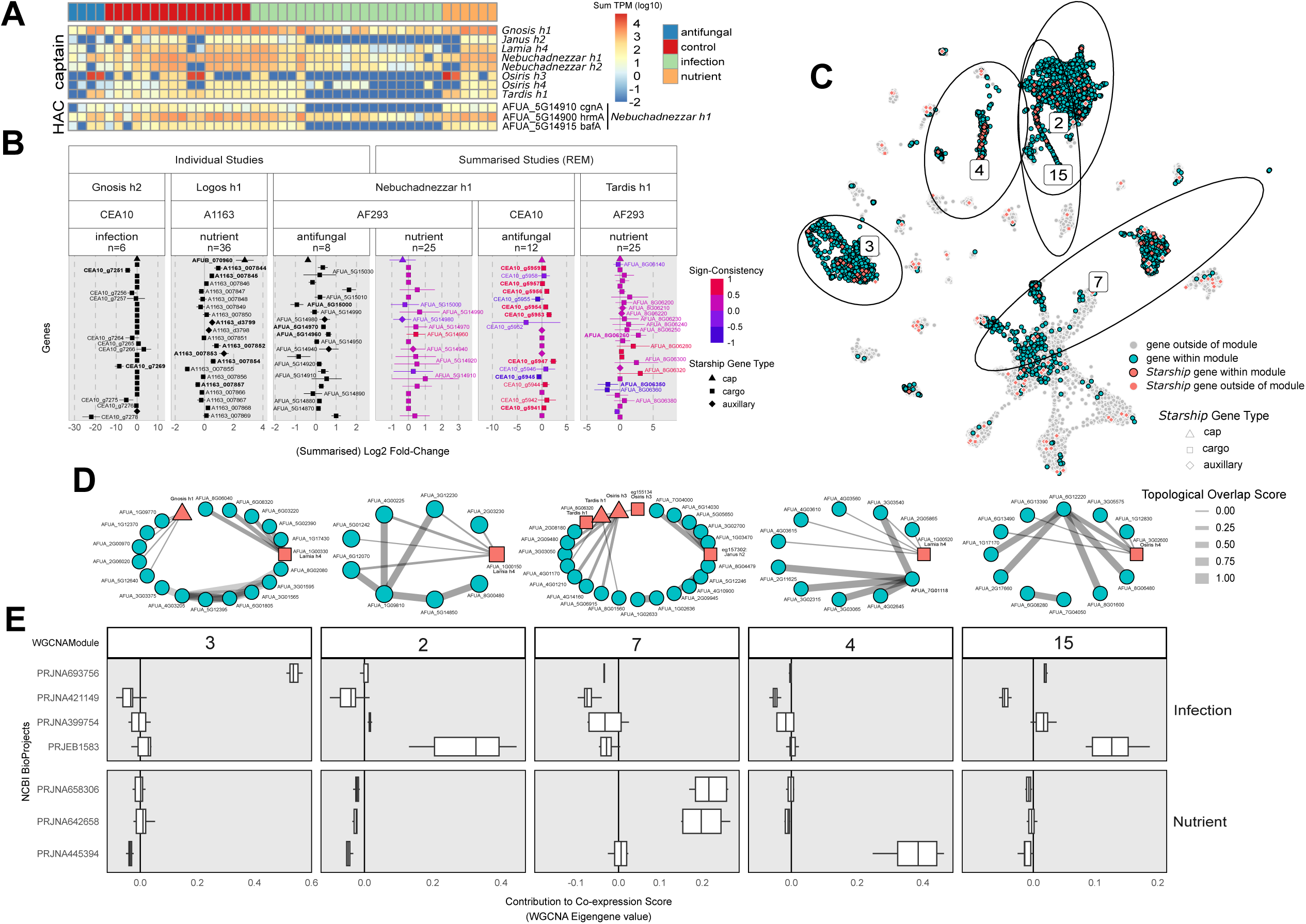
A) A heatmap of transcript abundances (log10TPM) of *Starship* captain tyrosine recombinase genes (top) and genes within the *hrmA*-associated cluster (HAC; bottom), collapsed across treatment replicates, for 11 RNAseq studies from *A. fumigatus* Af293. B) Results from differential expression tests for select *Starships*, treatment categories, and strain combinations. Differentially expressed genes (DEGs) based on a single study are shown as log2 fold-change (log2FC) in black with standard error bars, whereas DEGs identified across multiple studies are represented with summarized log2FC values from a random effects model (REM) and coloured by “sign-consistency”, the number of studies that reported DEGs in the positive (+1) or negative (-1) direction, centered around 0. Labels in bold font represent DEGs that are significantly differentially expressed in more than one study. C) The results of a weighted gene co-expression network analysis (WGCNA) constructed from 14 “antifungal”, “infection”, and “nutrient” RNAseq studies represented using UMAP clustering (90) based on the co-expression eigengene values for all genes in the *A. fumigatus* Af293 genome. Non-*Starship* genes present within modules of interest are shown in blue, while all *Starship* genes are shown in red, with those genes within modules of interest having a black outline. D) Genes within modules that were significantly associated with samples from specific treatment categories or studies were used to construct sub-networks which contained the top 10 edges in the network that were made between any pair of genes or any gene and a *Starship* captain. The connections between genes (edges) are based on the topological overlap matrix (TOM) for each module, and have been 0-1 scaled. E) Boxplots of eigengene values from WGCNA, akin to a weighted average expression profile, indicate the extent of co-expression (correlation) of the genes present within each module. Pairwise comparisons of module eigengene values determined if a module was significantly associated with samples from specific treatment categories or studies (Fig. S16).

In order to identify conditions where *Starships* have potential to impact phenotypes, we next examined patterns of differential expression for *Starship-*mobilized cargo genes. Overall, we did not find a significant enrichment of DEGs within *Starships* compared to the genomic background of any strain (Fisher’s Exact Test *P*-values > 0.05). However, the expression and significance of *Starship* DEGs vary across *Starship* naves, fungal strains, and experimental conditions, indicative of pervasive strain-specific, treatment-specific and strain by treatment interactions impacting *Starship* cargo expression (Fig. 5B, Fig. S14). For example, we compared transcript abundance patterns for genes within the *hrmA*- and *baf-*associated cluster (46) between treatment categories and found *cgnA* and *bafA* to be down-regulated in the majority of samples assigned to infection treatments but not in control or nutrient treatment samples (Fig. 5A, Fig. S12). Together, fungal strain identity, treatment condition, and non-additive interactions between fungal strains and treatments generate variation in *Starship* cargo expression.

To gain further insight into *Starship*-mediated heterogeneity in gene expression and regulation, we constructed weighted gene co-expression networks (WGCNA) which visualize correlations in gene expression. Genes are grouped into modules based on their level of shared co-expression patterns across all samples (Methods). We found five modules as being significantly associated with “antifungal”-, “infection”-, or “nutrient”-based studies (Fig. 5E). We identified two major trends in the transcriptional network properties of *Starship* cargo. First, multiple *Starship*-associated genes are integrated into modules containing many other genes from the genome, underscoring the possibility that phenotypes emerging from these networks are the product of regulatory interactions between *Starship* cargo and non-*Starship* genes (Fig. 5D, Fig. S15). In particular, connections involving captain genes hint at candidate loci involved in regulatory interactions between *Starships* and the *A. fumigatus* genome and are prime targets for future investigations into the regulation of *Starship* transposition (Fig. 5E). Second, we also identified modules composed of mainly *Starship* cargo, indicative of modular networks specific to particular *Starships* (Table S24). Together, our network analyses paint a nuanced picture of how *Starships* interact with the broader regulatory networks of the cell and implicate them in the generation of transcriptional variation.

### Relevance and Outlook

Phenotypic heterogeneity among fungal pathogen strains poses a major challenge for combating infectious diseases, yet we often know little about the genetic basis of this variation. We hypothesized that a newly discovered group of unusual transposons, the *Starships*, make important contributions to strain heterogeneity. We tested this hypothesis by systematically characterizing *Starship* presence/absence and expression in *A. fumigatus*, an important human fungal pathogen of critically high research priority (1, 48). Our work provides fundamental insight into the mode and tempo of fungal evolution by revealing that strain variation emerges not only from well-studied genetic mechanisms (e.g., single nucleotide polymorphisms, copy number variation, chromosomal aneuploidies, and other types of structural variants) (10, 11) but from the previously hidden activity of giant *Starship* transposons (Fig. 6).

**Figure 6:**
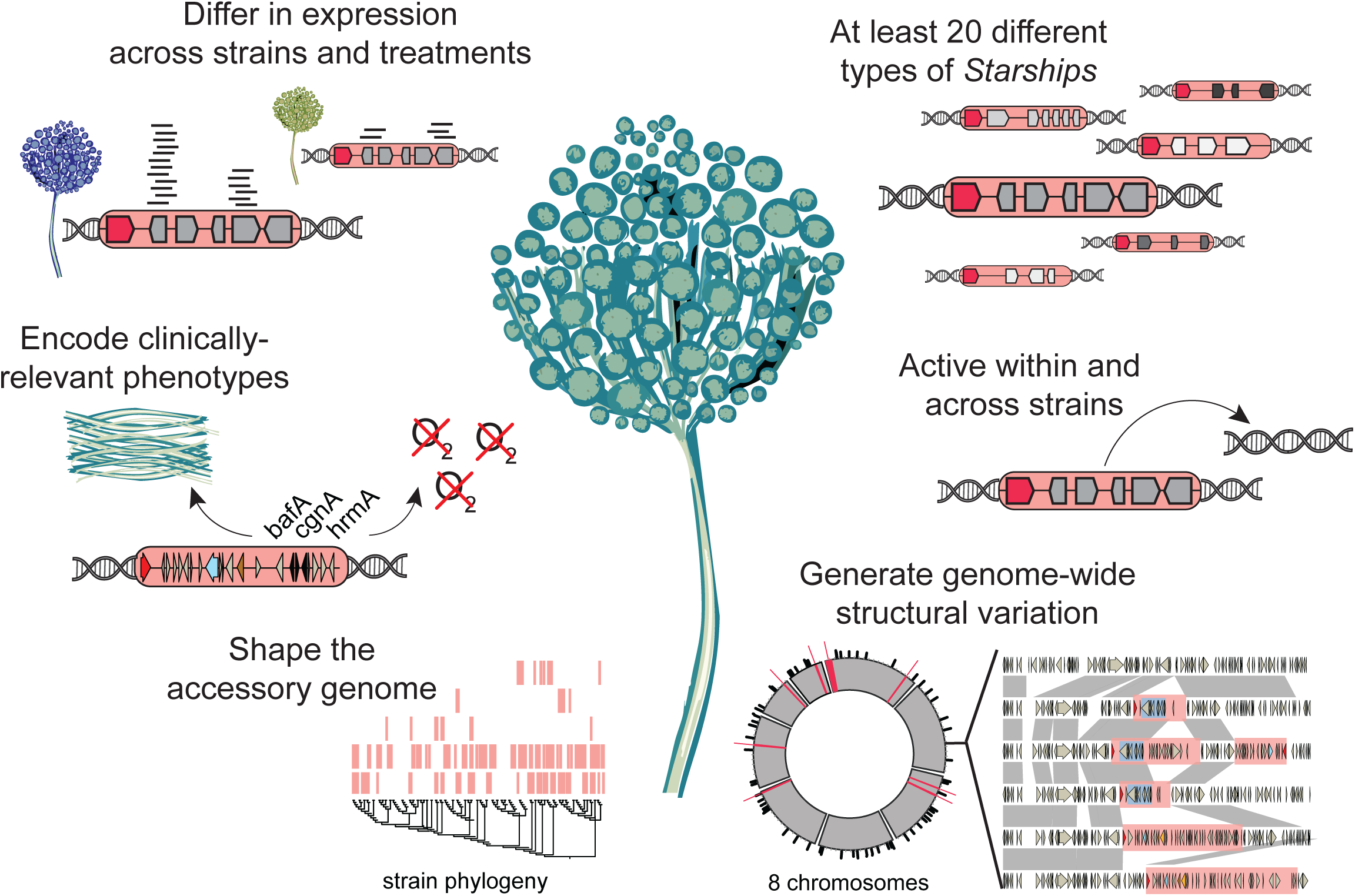
Contributions of *Starships* to strain heterogeneity in *Aspergillus fumigatus*.

Although much remains unknown about the fundamentals of *Starship* biology, *Starships* in *A. fumigatus* likely generate variation on a scale approaching MGE-mediated evolution in bacteria, which is a major mechanism driving bacterial adaptation. The median number of *Starship*-borne genes per *A. fumigatus* isolate is 0.8% (range = 0 - 2.1%), while between 2.9-4.8% of the pangenome (corresponding to between 9.7-16% of the accessory genome) is *Starship*- associated. These observations approach analogous measurements of bacterial plasmids in the Enterobacteriaceae, where the median number of plasmid-borne genes per genome is estimated at 3.3% (range = 0 - 16.5%) and where between 12.3 - 21.5% of the pangenome is plasmid-associated (49). Given our requirements that a high-confidence *Starship* be found in 2 different locations in 2 different strains, our estimates represent a relatively conservative assessment of the mobile fraction of *A. fumigatus’* genome. Mobility underpins prokaryotic genome dynamics, yet current models of fungal pathogen biology and evolution do not take gene mobility through transposition and horizontal transfer into account. These data highlight the need to understand the population level impacts of *Starships* across different species and to integrate these findings into a new predictive framework for fungal biology. Our *Starship* compendium thus provides practical insights for elucidating *A. fumigatus* biology and establishes a roadmap for deciphering the origins of strain heterogeneity across the fungal tree of life.

## Materials and Methods

### Long-read genome sequencing, assembly and annotation

For DNA extraction strains were grown at 37 °C for 24 hours in shaking liquid cultures using 1% glucose minimal media (50). Biomass was collected by gravity filtration through Miracloth (Millipore). High molecular weight DNA extraction followed the Fungal CTAB DNA extraction protocol (51). Biomass was ground in liquid nitrogen and incubated at 65 °C in a lysis buffer composed of 650 µl of Buffer A (0.35 M sorbitol, 0.1 M Tris-HCl pH 9, 5 mM EDTA pH 8), 650 µl Buffer B (0.2 M Tris-HCl pH 9, 0.05 M EDTA pH8, 2M NaCl, 2% CTAB), 260 µl of Buffer C (5% Sarkosyl), 0.01% PVP, 0.2 mg proteinase K for 30 minutes. Potassium acetate (280 µl of 5M solution) was added and incubated on ice for 5 minutes. DNA was extracted by phenol:chloroform:isoamyl alcohol extraction followed by a secondary extraction with chloroform:isoamyl alcohol. The supernatant was RNAse treated (2.5µg) and precipitated using sodium-acetate (0.3M) and isopropanol. DNA was finally purified using 70% ethanol, dried and rehydrated in TE (pH 9). To preserve long strands of DNA suitable for long-read sequencing samples were gently mixed by inversion and transferred using large bore pipette tips. DNA quality was assessed using NanoDrop and concentration quantified using Qubit dsDNA BR assay kit. Finally, DNA fragment size was assessed on 1% agarose gel, looking for large quantities of DNA to remain in the well after 1 hour of running at 100v. DNA was sent to SeqCoast for Oxford Nanopore Sequencing. We used previously sequenced Illumina short read data for polishing the assemblies with Pilon (10). Two strains not previously sequenced were sequenced with Illumina at SeqCoast.

The long-read genome assemblies were assembled with Canu v2.2 and polished with five rounds of Pilon v1.24 wrapped with AAFTF v0.3.0 (52–54). Summary statistics for each assembly were computed with AAFTF and BUSCO v5.7 using eurotiomycetes_odb10 marker set (55). Genome annotations (including for the publicly available CEA10 assembly) were performed using Funannotate v1.8.10 which trained gene predictors with PASA using RNA-Seq generated for each strain (Puerner et al., forthcoming), and predicted genes de novo in each assembly and produced a consensus gene prediction set for each strain (56, 57).

### Functional predictions

We annotated all predicted genes with Pfam domains, InterPro domains and GO terms using InterproScan v5.61-93.0 (-appl Pfam --iprlookup --goterms) (58). We annotated carbohydrate active enzymes using dbcan v3.0.4 (--tools all) and EggNOG orthogroups and COG categories using eggnog-mapper v2.1.7 (--sensmode very-sensitive --tax_scope Fungi)(59, 60). We annotated biosynthetic gene clusters using antiSMASH v6.0.1 (--taxon fungi --clusterhmmer -- tigrfam) and 5S rDNA using infernal v1.1.4 (--rfam -E 0.001)(61, 62). On average, 35.7% of genes within *Starships* have no known PFAM or Interpro domain, GO term, COG category or eggNOG orthology (range = 0-67.6%, std. dev. = 14.8%) but approximately half of all mobilized genes could be assigned to a COG (e.g., 50% of 11,900 genes total, across 459 high-confidence elements). To investigate associations between *Starships* and pathogen phenotypes, we searched for the presence of 798 genes known to be associated with virulence or infection-relevant phenotypes on *Starships* (Table S19, Methods)(10, 11, 36, 38–44).

### SNP analysis

We calculated pairwise kinship among strains by analyzing a previously generated VCF of all high-quality, filtered, biallelic single nucleotide polymorphisms (SNPs) from the Lofgren et al. 2022 population with plink v1.9 (--distance square ibs flat-missing --double-id --allow-extra-chr; Fig. S3) (63, 64). We calculated Jaccard similarity in *Starship* repertoire among all pairwise combinations of strains using genotyping data for the set of 20 high-confidence *Starships* and the following formula: *Starships* shared between strain 1 and 2 / (*Starships* unique to strain 1 + *Starships* unique to strain 2 + *Starships* shared between strain 1 and 2).

### Pangenome construction

We constructed a pangenome for the 13 reference-quality strains (our 11 long-read assemblies plus the Af293 and CEA10 references) combined with the 506 publicly available short-read assemblies (10, 11) using Orthofinder v2.5.4 (--only-groups -S diamond), and then extracted all orthogroups with a sequence from at least 1 of the 13 reference-quality strains (Fig. 2B)(65). Core orthologs are defined as being present in all 13 strains; accessory orthologs are present in between 2-12 strains; singleton orthologs are present in 1 isolate. We included all 519 strains in the Orthofinder analysis to ensure that ortholog groups are consistent across different population subsets.

### BLAST analysis

We determined the genomic locations of all best reciprocal blast hits to cargo genes carried by the 20 high-confidence *Starships* using BLASTp v2.13 (Fig. 1C) (66). For each cargo gene, we retrieved the highest scoring hit with ≥90% identity and ≥90% query coverage in each of the 13 reference-quality assemblies and determined whether that hit was found in the same *Starship*, a different *Starship*, or not in a *Starship* at all. If no hit was retrieved, that gene was marked as missing.

### Within strain analysis

To assess *Starship* transposition among isolates of the same strain, we compared 6 independently sequenced datasets of CEA10 and CEA10-derived strains to the long-read CEA10 reference assembly. Genome codes and NCBI accessions are as follows: CEA10-1: SRR7418934, CEA10-2: ERR232423, CEA10-MCH: ERR232426, A1160: SAMN28487500, A1163: SRR068950, A1163-Cramer_SM: SRX27799936. We evaluated three additional strains that also met the criteria of having a reference quality long-read assembly and at least 1 independently generated sequencing dataset generated by other research groups: ATCC46645, S02-30 (=AF100-9B), and TP9 (=08-19-02-30). Short-read sequence data were mapped to our long-read assemblies with bwa-mem2 and visually inspected in IGV to determine if any *Starships* were deleted/transposed (67). Short-read data from select isolates with evidence of transposition were *de novo* assembled with SPAdes and existing gene annotations were mapped from long-read assemblies using liftoff (68, 69).

### *Starship* annotation

We systematically annotated *Starships* in the 12 newly sequenced long-read genomes plus 507 publicly available *A. fumigatus* genomes by applying the starfish workflow v1.0.0 (default settings) in conjunction with metaeuk v14.7e284, Mummer v4.0, CNEFinder and BLASTn (Table S2) (22, 66, 70–72). Briefly, captain tyrosine recombinase genes were *de novo* predicted with starfish’s Gene Finder Module and full-length elements associated with captains were predicted with starfish’s Element Finder Module using pairwise BLAST alignments to find empty and occupied insertion sites. We filtered out all elements that were <15kb in length, which we have found to correspond to indels of captain genes but never to the transposition of a full-length *Starship* (22). We then manually examined alignments between each putative *Starship* and its corresponding insertion site and filtered out all “low confidence” poorly supported alignments indicative of a false positive insertion. Poor alignments were observed most often when *Starships* inserted into smaller transposons; when an inversion breakpoint occurred within a putative *Starship*, resulting in an over-estimation of *Starship* length; or when the insertion site was on a very small contig resulting in small flanking region alignments.

We verified and supplemented these automated *Starship* predictions with manual annotations of the 3 reference strains Af293, CEA10 and A1163, along with a set of 86 additional elements (Table S7, Table S3). Insertion sites and DRs were manually verified by generating alignments of the elements plus the 50kb flanks to a corresponding putative insertion site, as determined by starfish (Table S4). MAFFT v7.4 was used to generate alignments with default parameters (73). The DRs were visually assessed and the insertion regions were examined for the presence of TEs, which could lead to erroneous target site, DR and TIR determinations. For *Starships* in the three reference genomes, the set of reference *Starships* was used as a query with BLAST to verify their start and end coordinates, and insertion sites. Additionally, the starfish output was examined for additional elements which did not meet our initial strict cutoffs to attempt to capture the entire *Starship* repertoire of these strains.

### *Starship* classification

*Starships* were then grouped into *naves* (singular: *navis*, latin for “ship”) by clustering captain sequences in ortholog groups using mmseqs easy-cluster (--min-seq-id 0.5 -c 0.25 --alignment-mode 3 --cov-mode 0 --cluster-reassign)(74). *Starship* sequences were then grouped into *haplotypes* using the MCL clustering algorithm in conjunction with sourmash sketch (-p k=510 scaled=100 noabund) that calculated pairwise k-mer similarities over the entire sequence length, as implemented in the commands “starfish sim” and “starfish group”(75, 76). These automated and systematic predictions yielded a core set of 787 individual elements grouped into 54 distinct *navis-haplotype* combinations.

We identified segregating insertion sites associated with the 787 elements across all 519 strains using the command starfish dereplicate (--restrict --flanking 6 --mismatching 2) in conjunction with the Orthofinder orthogroups file (see above) that was filtered to contain groups absent in at most 517 strains and present in at most 8 copies per isolate. Starfish dereplicate enables the identification of independently segregating insertions by grouping *Starships* and their insertion sites into homologous genomic regions using conserved combinations of orthogroups between individuals. We genotyped the presence of each *navis-haplotype* combination within each region for each isolate. If an isolate did not have any *Starships* within a given region, it was assigned either an “empty” genotype (if the set of orthogroups that define the upstream region flank were adjacent to the set of orthogroups that define the downstream region flank) or a “fragmented” genotype (if additional orthogroups were in between the upstream and downstream region flanks).

To define high-confidence *Starships,* we used a threshold that required observing the same *navis-haplotype* in ≥2 independent segregating sites in different strains, which ensured an accurate and precise determination of each element’s boundaries and the sequences therein (22). We found 18 *navis-haplotype* combinations meeting these criteria. We considered 2 additional *navis-haplotype* combinations of interest (*Osiris h4* and *Lamia h4*) to be high-confidence after manually annotating their boundaries and finding evidence for flanking direct repeats that are signatures of *Starship* boundaries. Each high-confidence *Starship* is represented by a type element (similar in concept to a type specimen for defining a biological species) that constitutes the longest element assigned to that *navis-haplotype*. The type elements of the reference *Starships* range in size from 16,357-443,866 bp and collectively represent 459 individual elements with an average length of 71,515 bp (Table S4, Table S5). The remaining 328 elements have an average length of 50,249 bp and are represented by 34 “medium-confidence” *navis-haplotypes* that were only observed at a single segregating site. We predicted the target sites of each high-confidence *Starship* by manually aligning multiple empty insertion site sequences to each other and identifying columns within 25bp of the core motif that had conserved nucleotides in at least 80% of sequences (Table 1).

Each *navis* associated with a high-confidence element was assigned a charismatic name (e.g., *Tardis*) and a new, sequentially named *haplotype* (*h1, h2*, etc.) to distinguish it from the automatically generated *haplotype* codes (*var14, var09*, etc.). All other 34 *navis-haplotype* combinations with evidence of only a single segregating site were classified as “medium-confidence”. Medium-confidence *Starships* with captains not belonging to the high-confidence naves kept their automatically generated *navis* and *haplotype* codes (e.g., *navis01-var09*), while medium-confidence *Starships* with captains belonging to the high-confidence naves were assigned to that *navis* but kept their automatically assigned *haplotype* (e.g., *Osiris-var04*).

### *Starship* genotyping

Starfish requires a well-resolved empty insertion site in order to annotate the boundaries of a contiguous *Starship* element. It will therefore not annotate any element whose corresponding empty site is missing, or any element that is partially assembled or not assembled at all. We therefore supplemented starfish output using a combination of BLASTn and short-read mapping to the library of reference *Starship* sequences to decrease the false negative rate for presence/absence genotyping. First, we recovered full-length and partial elements using the set of high-confidence *Starships* as input to the command “starfish extend”. This command uses BLASTn to align full-length *Starship* elements to the sequence downstream from all tyrosine recombinase genes not affiliated with a full-length *Starship* element.

Separately, we downloaded publicly available paired-end short read Illumina sequencing data for all 466 strains for which these data were available (10, 11) and mapped them to the library of 20 high-confidence *Starships* using the command “starfish coverage” and the aligner strobealign (77), ensuring that we would correctly genotype the presence/absence of a *Starship* even if it is not present in an assembly or partially assembled. By mapping short reads directly to a library of known *Starship* elements, we overcame some of the drawbacks associated with working with short-read assemblies. We considered any *Starship* with a minimum of 5 mapped reads at each position across 95% of its entire length to be present. We considered a *Starship* to be present in a given individual if either the main starfish workflow, BLASTn extension, or short-read mapping identified it as present; otherwise we considered it absent.

### Meta-Analysis of *A. fumigatus* RNAseq Datasets

We collected transcriptomes from a total of 434 publicly available paired-end libraries of *A. fumigatus* strains Af293, A1163, and CEA10/CEA17 (Table S22). We surveyed the available metadata from each BioProject and binned the samples into general categories based on the type of treatment applied in each study. The categories that were the most well represented across multiple BioProjects include exposure to antifungals (“antifungal”), *in vivo* and *in vitro* infection experiments (“infection”), or supplemented growth media with specific nutrients (“nutrient”; Table S22, Table S23).

RNAseq reads were retrieved from the NCBI SRA using fasterq-dump from the SRA toolkit (last accessed June 26, 2024: https://github.com/ncbi/sra-tools) and were trimmed for quality and sequencing artifacts using TrimGalore (last accessed June 26, 2024: https://github.com/FelixKrueger/TrimGalore). Transcripts were quantified using salmon quant, which employs a reference-free pseudo-mapping based approach to quantify transcripts. In addition to the pseudo-BAM files created by salmon v1.10.1 (78), we created separate sets of BAM files to assess transcript coverage across Starship genes by mapping to the transcriptome with STAR v2.7.11a (79). We assessed transcript coverage across the core of Starship genes using the bamsignals R package (last accessed June 26, 2024: https://github.com/lamortenera/bamsignals. Genes with a median core-transcript coverage less than 1 were considered to have insufficient evidence of being expressed. We applied an abundance filter on transcript abundances, removing genes represented by fewer than 10 transcripts in at least 3 samples. We corrected for batch effects between BioProjects using CombatSeq (80) and excluded BioProjects with inconsistent/unclear metadata, or those that remained as outliers in the PCA after batch-correction. DESeq2 v1.45.0 (81) was used to perform a series of differential expression tests using the corrected transcript counts. These tests were performed individually for each BioProject, with each test comparing the differences in expression between all control and treatment samples. Differences in treatment level were included as a cofactor in the model, where applicable.

To summarize expression patterns across BioProjects which tested similar conditions, we employed a random effects model (REM) using the R package metaVolcanoR (82). This method identifies differentially expressed genes (DEGs) as genes that are significantly differentially expressed in all studies and accounts for the variation in the expression for each DEG observed in multiple BioProjects. The output from the meta-volcano REM are summarized log2 fold-change, a summary p-value, and confidence intervals for each DEG. In addition, a measure of “sign-consistency” is used to evaluate the consistency of DEG, which is expressed as a count of the number of BioProjects where a DEG was observed with the same directional change (+/-), centered around 0.

We performed weighted gene co-expression network analysis (WGCNA) using PyWGCNA v1.72-5 (83) to construct gene co-expression networks for *A. fumigatus* reference strain Af293. A single network was constructed using the batch-corrected TPM values from the collection of samples belonging to “antifungal”, “infection”, and “nutrient” treatment categories. PyWGCNA automatically estimates an appropriate soft-power threshold based on the lowest power for fitting scale-free topology and identifies modules of co-expressed genes through hierarchical clustering of the network and performing a dynamic tree cut based on 99% of the dendrogram height range. The topological overlap matrix is then computed and a correlation matrix is constructed to produce the final network. Distributions of TOM scores were generated based on edges in the network between genes within *Starships*, between *Starships*, between *Starships* and the rest of the genome, and between non-*Starship* genes in the genome. We compared these distributions using a Wilcoxon test and anova, adjusting p-values with the Holm–Bonferroni method. Modules in the network that were significantly associated with samples from a specific treatment category were identified using the pairwise distances between observations using pdist from the python module scipy (84). To highlight the genes within each module that have the most strongly correlated expression profiles, sub-networks were constructed using only the edges 10 highest TOM scores made between either *Starship* captain genes or any non-*Starship* gene that was present in the module. The collections of genes within these modules were also tested for functional enrichment using gprofiler2 (85).

### Statistical tests, alignments and data visualization

Enrichment tests were performed with either the Binomial test (binom.test; alternative = “greater”) or the Fisher’s exact test (fisher.test; alternative = “greater”) implemented in base R, using the p.adjust function (method = Benjamini-Hochberg) to correct for multiple comparisons. Mantel tests for matrix correlations were performed in R (method = “pearson”, permutations = 999). We conducted nucleotide alignments among *Starships* and their associated genomic regions (e.g. Fig. 1E, Fig. 3D) using Mummer 4.0 with the nucmer command with delta-filter options -m -l 5000 -i 95 (70). We visualized all alignments and insertion site distributions using circos and gggenomes (86, 87) with the help of wrapper script functions implemented in the function starfish locus-viz in the starfish package v1.1 (22). Step-by-step tutorials for *Starship* data visualization are maintained on the starfish github repository available at https://github.com/egluckthaler/starfish/wiki. All other figures were generated in R using ggplot2 (88).

## Data availability

Sequencing reads and genomic assemblies for the 11 isolates sequenced with Oxford Nanopore have been deposited and accessioned at NCBI (Table S1). Short-read sequence data for the A1163-Cramer_SM isolate are deposited at NCBI under Bioproject PRJNA1228067. We downloaded the Af293 and CEA10 reference assemblies (NCBI accessions: GCF_000002655.1 and SAMN28487501) as well as 252 assemblies and annotations from Barber et al. 2021 from NCBI, and 256 assemblies and annotations from Lofgren et al. 2022 from the associated Zenodo repository (64). We subsequently reformatted the contig and gene identifiers to include genome codes generated as part of this study (Table S2). All scripts (including raw data for figure generation), genomic data used for this study, and results from differential expression tests are available through the following Figshare repository DOI: 10.6084/m9.figshare.26049703.

## Acknowledgements

We thank Jacob Steenwyk for insightful feedback on an earlier draft of this manuscript.

## Supplemental figure legends

Figure S1: Counts of starfish-predicted *Starships* per *Aspergillus fumigatus* strain (n = 519), broken up by assembly project (either the 12 Oxford Nanopore Assemblies generated by study plus the AF293 reference genome, or from Lofgren et al 2022 or Barber et al 2021; Table S2). A) Counts of *Starships* with either ‘insert’ or ‘flank’ boundaries, which are derived directly from pairwise genome alignments against a putative insertion site. B) Counts of *Starships* with ‘insert’ or ‘flank’ boundaries plus those with ‘extend’ boundaries, which are derived from aligning genomic sequences to known *Starship* sequences. Insert and flank boundaries are associated with full-length *Starship* elements, while “extend” boundaries may be associated with either full-length *Starship* elements or element fragments.

Figure S2: Nucleotide alignments of representative *Starship* copies, +/- 100kb of flanking sequence, across all genomic regions where that *Starship* is found. Predicted functions of interest are annotated above the corresponding gene. Links between schematics represent alignable regions ≥1000bp and ≥90% nucleotide sequence identity. A) *Tardis*. B) *Gnosis*. C) *Janus*. D) *Osiris*. E) *Lamia*. F) *Logos*. G) *Nebuchadnezzar.*

Figure S3: A scatterplot depicting pairwise comparisons of single nucleotide polymorphism (SNP)-based Identity by State (IBS) and Jaccard similarity in high-confidence Starship presence/absence profiles between 151 Clade 1 strains from Lofgren et al. 2022 (Table S8).

Figure S4: Screenshots from the IGV genome browser showing deletions of *Nebuchadnezzar h1* among different isolates of the ATCC46645 strain sequenced by different research groups (regions depicted in red denote *Nebuchadnezzar h1*). Illumina short-reads of strain ATCC46645 sequenced by a different research group (accession: SRR7418935) mapped to the ATCC46645 genome sequenced with long read technology in this study. A zoom in of the genomic location of *Nebuchadnezzar h1* in the ATCC46645 long read assembly is shown (contig accession: ATCC46645-lr_scaffold15). Three short-read tracks indicate a deletion of the *Starship* (above). Note that more short reads are mapped than shown in the image. Track and color descriptions can be found in the IGV manual.

Figure S5: Donut charts summarizing the percentages of gene orthogroups in the core, accessory, singleton, and *Starship-*associated compartments of the *Aspergillus fumigatus* pangenome, derived from the 13 reference-quality *A. fumigatus* strains and the expanded set of 54 high and medium-confidence *Starships* (n = 1818 elements total; Table S11).

Figure S6: Iceberg plots summarizing the genomic locations of the single best BLASTp hits (≥90% identity, ≥33% query coverage) to the cargo genes from the type elements of the 20 high-confidence *Starships* in 519 *Aspergillus fumigatus* assemblies (Table S13). Each column represents a cargo gene. A) Results broken down by *Starship*, with columns arranged according to gene order within each *Starship*. B) Compiled results across all 20 *Starships*, with columns arranged according to the number of strains with BLASTp hits. Bars are colored according to the genomic location in which the BLASTp hits are found.

Figure S7: Barcharts summarizing the genotypes of segregating genomic regions associated with the 20 high-confidence elements in the *Aspergillus fumigatus* 519 strain population (Table S16). If an isolate did not have any Starships within a given region, it was assigned either an “empty” or “fragmented” genotype (Methods).

Figure S8: The putative target site of *Starship Tardis* (indicated with an arrow) occurs more often than you would expect by chance in the *Aspergillus fumigatus* genome. We estimated the copy numbers of all k-mers of length 10 in the Af293 reference genome and found that the k-mer of length 10 that corresponds to the putative target site of *Tardis* is present in high copy numbers that exceed the expected genome-wide frequency of k-mers of this length (>99.99th percentile).

Figure S9: *Starships* mobilize genes encoding diverse metabolic functions. Barcharts summarizing the presence of genes with metabolism-related COG (Clusters of Orthologous Groups) annotations in the 20 high-confidence *Starships* in the 519 *Aspergillus fumigatus* strain population (Table S18). The X-axis measures the percentage of copies of a given *Starship* that carry at least 1 gene with an annotation of interest.

Figure S10: The Biofilm Architecture Factor (*baf*) gene family is closely associated with diverse *Starships* in *Aspergillus fumigatus*. A maximum likelihood tree of *baf* sequences from the 519 *Aspergillus fumigatus* strain population. Branches with ≥80% SH-ALRT and ≥95% ultrafast bootstrap support are in bold. *Baf* sequences found in *Starships* have the corresponding *Starship* identification number appended to their right (Table S5). Sequences are color-coded according to 6 corresponding *baf* clades of interest, and a summary of all the *Starship* types found associated with each clade is printed on the right.

Figure S11: *Starships* are enriched in environmental and clinical strains. Barcharts summarizing the proportion of strains from 475/479 *Aspergillus fumigatus* genotyped strains with known isolation sources that have the 20 high-confidence *Starships*, broken up by isolation source. Fisher’s exact test P values for *Starship* enrichment across isolation source categories are shown above each bar (adjusted for multiple comparisons using the Benjamini Hochberg procedure; Table S20).

Figure S12: A heatmap of transcript abundances (log_10_ TPM) of *Starship* captain genes (A, C) and genes within the *hrmA*-associated cluster (HAC; B, D), collapsed across treatment replicates, for RNAseq studies from *A. fumigatus* CEA10 (A, B) and A1163 (C, D).

Figure S13: Heatmap of binary core transcript coverage across genes in *Starships* in strains Af293 (A), CEA10 (B) and A1163 (C). Genes that have a median core transcript coverage greater than 1 are shown here in blue, and values below 1 are shown in red.

Figure S14: Differentially expressed *Starship*-mobilized genes (DEGs) displayed in volcano plots across combinations of *A. fumigatus* strains and treatment categories. Differential expression based on singleton studies are shown as log_2_ fold-change (log_2_FC) in black with standard error bars, whereas DEGs identified across multiple studies are represented with summarized log_2_FC values from a random effects model (REM) and coloured by values “sign-consistency”, the number of studies that reported differential gene expression in the positive (+1) or negative (-1) direction, centered around 0.

Figure S15: Distributions of TOM scores from the WGCNA constructed from Af293 *A. fumigatus* studies. TOM scores were separated into distinct categories for edges that connected any two genes within a *Starships*, between different *Starships*, between *Starships* and the rest of the genome, or between any two non-*Starship* genes in the genome were compared. These distributions were compared using a Wilcoxon test (p > 0.05 = “ns”; p < 0.05 = “*”, p < 0.01* = “**”, p < 0.001 = “***”, p < 0.0001 = “****”).

Figure S16: Heatmap displaying the correlation values for eigengenes in each module and their association with samples from a specific study (labeled with BioProject IDs), or different levels of experimental treatment category across studies. Cells of the heatmap are coloured by the correlation value, as well as labeled in each cell. P-values from a pairwise comparison between observations (*pdist*) are shown in parentheses.

## Supplemental table legends

Table S1: Genome assembly statistics for the 12 strains sequenced with Oxford Nanopore long-read technology

Table S2: Metadata for publically available *Aspergillus fumigatus* strains Table S3: Manual annotation data of 86 Starship elements

Table S4: Metadata for all starfish-predicted Starships

Table S5: Sequence features of all starfish-predicted Starships in the 519 strain population

Table S6: Frequencies of the 20 high-confidence Starships in the 519 strain population using the expanded dataset

Table S7: Manual annotation of Starship elements in three *Aspergillus fumigatus* reference strains

Table S8: Pairwise comparisons between strains of SNP identity by state and Starship repertoire similarity

Table S9: Comparison of Starship coordinates with structural variants identified by Colabardini et al 2022 (doi: 10.1371/journal.pgen.1010001)

Table S10: Orthogroup frequencies in the 519 strain population Table S11: Orthogroup frequencies in the 13 reference-quality strains

Table S12: BLAST recovery of Starship cargo genes from the 13 reference-quality strains

Table S13: BLAST recovery of Starship cargo genes from the 519 strain population

Table S14: Genotyping of genomic regions with segregating Starship insertions

Table S15: Starships in genotyped genomic regions with segregating Starship insertions

Table S16: Summary of genotyping data for genomic regions with segregating Starship insertions

Table S17: Coordinates of predicted 5s rDNA sequences in the 519 strain population

Table S18: Proportion of elements carrying at least one gene annotated with various COG categories

Table S19: List of putative and published virulence and stress resistance genotypes in *Aspergillus fumigatus*

Table S20: Fisher’s exact test statistics for Starship enrichment by strain isolation source

Table S21: Presence/absence genotyping data for the 20 high-confidence Starships in the 519 strain population using the genotyping dataset

Table S22: Metadata from RNASeq studies collected from NCBI used in the meta-analyses of *Starship* gene expression.

Table S23: Summary of RNASeq studies collected from NCBI used in the meta-analyses of *Starship* gene expression.

Table S24: Summary of modules assigned, and genes within them, from the WGCNA constructed from Af293 samples.

Table S25: Significantly enriched functional terms from a series of enrichment tests (Fisher’s Exact Test) conducted on the genes within WGCNA modules.

## Supplemental information and results

### *Starship* frequencies

The population-level frequency of high-confidence *Starships* ranges from 1-60%, with 13 of the high-confidence *Starships* present in <10% of strains and no *Starship* present in more than 60% of strains (Table S6). Each high-confidence *Starship* is present across multiple individuals; however, any given *Starship* element is only present at most once in a genome, with only 3 exceptions that likely involve recently degraded copies, suggesting some mechanism for controlling their copy number (e.g., repeat-induced point mutation (21)). Individual elements typically segregate at low frequencies, with each segregating site containing on average an inserted element in just 0.4% of strains across the 20 high-confidence *Starships* (range = 0.2-52.3%, std. dev. = 6.5%, n = 154; Fig. 3B). However, multiple insertion sites often exist for a given element such that each high-confidence *Starship* is present on average in 4.35 different regions (range = 1-15, std. dev. = 3.92).

### Differential expression of *Starship* captains and cargo

Patterns of transcript coverage, transcript abundance, and differential expression of *Starship*-associated genes were generally heterogeneous, with various strain-, treatment- and strain by treatment interactions. The transcriptional profile for many *Starships* is patchy, and not all *Starship*-mobilized genes are expressed in response to experimental conditions. Only certain cargo genes have the minimum required evidence for constitutive expression: a median transcript coverage of 1 across the gene body. Nearly all genes in *A. fumigatus* CEA10 *Starships* surpass this threshold (Fig. S13B, across all samples from the various studies we included in our analyses. One notable exception being the captain gene in *Janus h2*, which is sparsely expressed across most studies. In comparison, genes present in *Starships* from Af293 (Fig. S13A) and A1163 (Fig. S13C) have fewer genes that surpass this threshold, with certain experimental conditions having more consistent expression of the *Starship* genes in these strains. For example, specific studies conducted on Af293 or CEA10, testing exposure to heatshock (PRJDB6203), hypoxia (PRJNA144647), *in vitro* infections (PRJEB1583, PRJNA399754, PRJNA399754). and various gene knock-out experiments (PRJNA390719, PRJNA396210, and PRJNA601094) have consistently reported expression of all genes across the *Starship* element.

Captain genes of multiple *Starships* from Af293 and CEA10 had low transcript abundance within “infection” studies. Furthermore, samples of a study performing an experimental infection using a murine model (PRJNA421149) had near-zero transcript abundances of captain genes from Af293 *Starships Janus h2*, *Osiris h3*, and *Gnosis h1*. However, this was not found within samples of a different study of an Af293 infection of human pneumocyte cell lines (PRJNA399754). Near-zero abundances were also found for a different set of captain genes, within CEA10 *Starships Tardis h1, Logos h2,* and *Gnosis h2*, from an *in vitro* infection study using human cell lines (Fig. S12), suggesting that the transcriptional response of *Starship* captains within the infection environment differs between strains.

We used tests of differential expression using existing *A. fumigatus* RNAseq studies to identify which environments or conditions influence the expression of genes within the *Starship* compartment (Fig. 5B, Fig. S14, Table S23). In particular, the change in expression of *Starship* captain genes within studies that applied similar types of treatments may have implications for what conditions may be more conducive for transposition. In general, multiple Af293 RNAseq “infection” studies have a prevailing negative impact on the expression of captain genes. Starships *Lamia h4*, *Tardis h1*, and *Gnosis h1* all had significantly lower expression within captain genes compared to their respective control samples (Figure 5B). We also observed a negative impact on captain expression within “antifungal” studies, which consisted of multiple studies conducted on Af293 and CEA10 with exposure to the antifungal caspofungin. Treatments of caspofungin were found to decrease the expression of Starship captains within Af293 Nebuchadnezzar h1 and CEA10 *Logos h2*. However, caspofungin was also found to be associated with increased expression of Af293 *Tardis h1* and *Gnosis h1* captain genes. Specific “nutrient” studies were also found to have a positive influence on captain gene expression. This includes a single study exposing A1163 to glucose/xylose media (PRJNA634102), which significantly increased the expression of captain genes within *Logos h1* and *Nebuchadnezzar h1*, compared to the control group grown on minimal media. Together, the differential expression results across these gene expression datasets demonstrate the context-dependent expression of captain genes, which we speculate would lead to variation in *Starship* transposition rates across different environments and strain backgrounds.

Similar to captain genes, the expression patterns of cargo genes within *Starships* are heterogeneous. The transcriptional profile of *Starships* is punctuated with expression of cargo genes (Fig. 5B, Fig. S14, Table S23), which can have sufficient transcript coverage even in the absence of captain expression, suggesting that the transcriptional regulation of these cargo genes are decoupled from transposition, and could be integrated into the existing genomic transcriptional networks. Of particular note are the genes that form biosynthetic gene clusters (BGC) which reside within *Starships*. In Af293, 63 BGC genes across seven Starships (*Gnosis h1, Lamia h4, Nebuchadnezzar h1, Nebuchadnezzar h2, Osiris h4, Tardis h1*, and *nav10 var35*) are differentially expressed in one or more of the treatment categories investigated in this study. At least one gene from the BGCs across these seven *Starships* are differentially expressed in response to caspofungin exposure, from a specific Af293 study (PRJNA472460). Furthermore, multiple genes present in Osiris h4 which are associated with another BGC (AFUA_3G02580, AFUA_3G02620, AFUA_3G02630, AFUA_3G02640) have significantly increased expression in nutrient studies compared to controls. Interestingly, generally fewer genes in BGCs are differentially expressed in Af293 “nutrient” studies.

As an opportunistic human pathogen, we were also interested in *A. fumigatus* genes that have specific relevance to virulence, such as genes within the *hrmA*-associated cluster (HAC). Two HAC genes present in *Nebuchadnezzar h1*, *cgnA* and *bafA*, have near-zero transcript abundances within samples from an Af293 study of an experimental infection of a murine model (PRJNA421149). However, genes within the HAC cluster were only differentially expressed within certain CEA10 antifungal studies and a single infection study: we found *bafA* in *Nebuchadnezzar h1* (CEA10_g5945) and *bafZ* in *Logos h1* (CEA10_g7202) to both have decreased expression in these studies, respectively (Fig. 5B, Fig. S14).

Certain cargo genes within Starships demonstrate patterns of differential expression in response to various experimental conditions, suggesting that these genes play a functional role in metabolic or enzymatic activity of the cell in response to specific stimuli. In Af293, we identified several Starship cargo genes with condition-specific expression patterns. Within *Nebuchadnezzar h1*, a proteoglycan 4 gene (AFUA_5G14930) showed increased expression during infection.

A group of cargo genes within Af293 *Lamia h4* displayed complex expression patterns across various treatment categories in this study. Within “infection” studies, multiple genes with diverse functions had decreased expression, including genes relevant to metabolic functions (AFUA_1G00800; flavin-containing amine oxidoreductase, AFUA_1G00340; thioesterase, and AFUA_1G00490; dihydrolipoamide acetyltransferase component of pyruvate dehydrogenase complex). One gene relevant to post-transcriptional modification also had decreased expression (AFUA_1G00730; DNAj domain protein). Few genes within Lamia h4 that serve potential roles in redox processes had increased expression within “infection” or “nutrient” treatments (AFUA_1G00440; DUF895 domain membrane protein and AFUA_1G00350; FAD-binding domains, respectively).

In CEA10, several genes showed increased expression with “antifungal” treatments, including genes containing NAD(P)-binding (CEA10_g9263; *Janus h2*) and PAS superfamily protein (CEA10_g5959; *Nebuchadnezzar h1*), and phosphotyrosine protein phosphatase (CEA10_g5954; *Nebuchadnezzar h1*). While several cargo genes were consistently downregulated in CEA10 *Logos h2*, including those encoding WD40 repeat proteins (CEA10_g5566), P-loop containing nucleoside triphosphate hydrolase (CEA10_g5567), homeodomain-like superfamily proteins (CEA10_g5587), and serine/threonine and tyrosine kinase (CEA10_g5577).

In response to “nutrient” conditions, several cargo genes within A1163 *Logos h1* were differentially expressed. with increased expression of serine threonine-protein (A1163_007845), DUF3723, MADS-box transcription factor (A1163_007852), and freB (A1163_007853). Conversely, a BTB domain-containing gene (A1163_007857), potentially involved in transcriptional regulation through chromatin structure control, showed decreased expression under the same conditions.

Additional cargo genes of interest may convey resistance to environmental stressors, including genes that provide resistance to environmental toxins. Genes incorporated in the arsenic detoxification pathway are present within *A. fumigatus* Starships *Nebuchadnezzar h1* within strains Af293 and CEA10. Arsenite methyltransferase genes were found to have significantly decreased expression in the Af293 copy (AFUA_5G15000) and significantly increased expression in the CEA10 copy (CEA10_g5956) with exposure to caspofungin (Fig. 5B). Collectively, these findings highlight the diverse functional roles of Starship cargo genes in responding to environmental stressors and nutrient availability, suggesting their importance in adaptability and survival of *A. fumigatus*.

### *Starship* co-expression networks

We performed weighted gene co-expression network analysis (WGCNA) using *PyWGCNA* to construct gene co-expression networks for *A. fumigatus* reference strain Af293. Overall, we found that genes within or between *Starships* are more strongly co-expressed together than with non-*Starship* genes, based on pairwise comparisons of the distribution of scores in the topological overlap matrices (TOM) (corrected p-values < 0.01 for all comparisons; Fig. S15). This suggests that a tighter transcriptional relationship exists between genes mobilized by *Starships* in Af293 compared to co-expression with other genes in the genome.

Genes in the *A. fumigatus* Af293 co-expression network were resolved into 27 modules (Table S24). The majority of genes found within each module are not contained within a *Starship*, yet almost all modules contain one or more *Starship* genes, including those previously identified as DEGs (Table S24). Modules were selected for further analyses based on the correlation of their gene expression profiles (WGCNA eigengenes) and their association with a treatment category (“antifungal”, “infection”, or “nutrient”) or specific study (Fig. 5E, Fig. S16).

To investigate functional compartmentalization of WGCNA modules, we performed enrichment tests and identified which GO/KEGG terms are significantly overrepresented within each module. Genes in module “2” are significantly enriched in functions including the production of antimicrobial secondary metabolites (fumagillin), the secretion of *A. fumigatus* mycotoxins (helvolic acid), proteins for heme binding, and synthesis of immunosuppressive compounds (endocrcoin) (Table S25).

To identify which genes are most strongly co-expressed with *Starship* cargo or captain genes, we subsetted the co-expression network to keep only the top 10 edges that were made between any pair of genes or any gene and a *Starship* captain. Two modules within the Af293 co-expression network (“2” and “15”) are significantly more commonly co-expressed within samples from a single infection study of Af293 (PRJEB1583; Fig. 5E). The connections with the highest TOM scores in module “2” include those between an IBR finger domain protein within the *Starship Lamia h4* (AFUA_1G00150) and genes involved in RNA binding pathways (AFUA_6G12070), as well as genes for alpha-amylase (AFUA_2G03230) and anthrone oxygenase (AFUA_4G00225) (Fig. 5D). Genes within the module “15” include those tightly connected to the expression of F-box domain protein within *Lamia h4*. Genes within module “15” are enriched in glycerophospholipid metabolism (Table S25). These co-expression relationships provide insight into how *Starships*-mobilized genes integrate into the existing transcriptional network in *A. fumigatus* strains.

Module “3” is significantly associated with increased expression within a single study of an experimental infection in mice (PRJNA693756) (Fig. 5E, S16). Genes within module “3” are enriched for genes in ribosome biogenesis in eukaryotes, carbon metabolism, and pyruvate metabolism (Table S25). The connections with the highest TOM scores in module “3” include genes co-expressed with *Gnosis h1* captain gene: genes with predicted catalytic activity (AFUA_1G12370), zinc-containing alcohol dehydrogenase (AFUA_2G00970), mitochondrial respiration (AFUA_2G06020), and exonuclease activity/DNA-directed DNA polymerase activity/role in mitochondrial DNA replication and mitochondrion localization (AFUA_5G12640). Module “7” is significantly associated with two studies testing supplementation with 5,8-dihydroxyoctadecadienoic acid (PRJNA658306) and lipo-chitooligosaccharides (PRJNA642658) in Af293 (Fig. 5E, S16). The focus of both of these studies is to understand how these supplementations impact the regulation of fungal growth and development. The connections with the highest TOM scores in the module “7”, include genes co-expressed with captain genes of *Tardis h1* and *Osiris h3. Osiris h3* captain co-expressed with predicted RNA binding, ribonuclease III activity and role in RNA processing (AFUA_3G03050). The connections made in this module with *Starship* genes present themselves as good candidates for future research to understand which genes are expressed along with transposition.

## References

1. 2022. WHO fungal priority pathogens list to guide research, development and public health action. World Health Organization, Geneva.

2. Denning DW. 2024. Global incidence and mortality of severe fungal disease. Lancet Infect Dis S1473–3099(23)00692–8.

3. Latgé J-P, Chamilos G. 2019. Aspergillus fumigatus and Aspergillosis in 2019. Clin Microbiol Rev 33:10.1128/cmr.00140-18.

4. Lin S-J, Schranz J, Teutsch SM. 2001. Aspergillosis Case-Fatality Rate: Systematic Review of the Literature. Clin Infect Dis 32:358–366.

5. Kowalski CH, Beattie SR, Fuller KK, McGurk EA, Tang Y-W, Hohl TM, Obar JJ, Cramer RA. 2016. Heterogeneity among Isolates Reveals that Fitness in Low Oxygen Correlates with Aspergillus fumigatus Virulence. mBio 7:e01515–16.

6. Colabardini AC, Wang F, Dong Z, Pardeshi L, Rocha MC, Costa JH, dos Reis TF, Brown A, Jaber QZ, Fridman M, Fill T, Rokas A, Malavazi I, Wong KH, Goldman GH. 2022. Heterogeneity in the transcriptional response of the human pathogen Aspergillus fumigatus to the antifungal agent caspofungin. Genetics 220:iyab183.

7. Horta MAC, Steenwyk JL, Mead ME, dos Santos LHB, Zhao S, Gibbons JG, Marcet-Houben M, Gabaldón T, Rokas A, Goldman GH. 2022. Examination of Genome-Wide Ortholog Variation in Clinical and Environmental Isolates of the Fungal Pathogen Aspergillus fumigatus. mBio 13:e01519-22.

8. Keller NP. 2017. Heterogeneity Confounds Establishment of “a” Model Microbial Strain. mBio 8:10.1128/mbio.00135-17.

9. Steenwyk JL, Rokas A, Goldman GH. 2023. Know the enemy and know yourself: Addressing cryptic fungal pathogens of humans and beyond. PLoS Pathog 19:e1011704.

10. Lofgren LA, Ross BS, Cramer RA, Stajich JE. 2022. The pan-genome of Aspergillus fumigatus provides a high-resolution view of its population structure revealing high levels of lineage-specific diversity driven by recombination. PLOS Biol 20:e3001890.

11. Barber AE, Sae-Ong T, Kang K, Seelbinder B, Li J, Walther G, Panagiotou G, Kurzai O. 2021. Aspergillus fumigatus pan-genome analysis identifies genetic variants associated with human infection. 12. Nat Microbiol 6:1526–1536.

12. Rhodes J, Abdolrasouli A, Dunne K, Sewell TR, Zhang Y, Ballard E, Brackin AP, van Rhijn N, Chown H, Tsitsopoulou A, Posso RB, Chotirmall SH, McElvaney NG, Murphy PG, Talento AF, Renwick J, Dyer PS, Szekely A, Bowyer P, Bromley MJ, Johnson EM, Lewis White P, Warris A, Barton RC, Schelenz S, Rogers TR, Armstrong-James D, Fisher MC. 2022. Population genomics confirms acquisition of drug-resistant Aspergillus fumigatus infection by humans from the environment. Nat Microbiol 7:663–674.

13. Hall JPJ, Harrison E, Baltrus DA. 2022. Introduction: the secret lives of microbial mobile genetic elements. Philos Trans R Soc B Biol Sci 377:20200460.

14. Aubert D, Naas T, Héritier C, Poirel L, Nordmann P. 2006. Functional characterization of IS1999, an IS4 family element involved in mobilization and expression of beta-lactam resistance genes. J Bacteriol 188:6506–6514.

15. Krishnan P, Meile L, Plissonneau C, Ma X, Hartmann FE, Croll D, McDonald BA, Sánchez-Vallet A. 2018. Transposable element insertions shape gene regulation and melanin production in a fungal pathogen of wheat. BMC Biol 16:78.

16. Hottes AK, Freddolino PL, Khare A, Donnell ZN, Liu JC, Tavazoie S. 2013. Bacterial Adaptation through Loss of Function. PLOS Genet 9:e1003617.

17. Weisberg AJ, Chang JH. 2023. Mobile Genetic Element Flexibility as an Underlying Principle to Bacterial Evolution. Annu Rev Microbiol 77:null.

18. Baquero F. 2004. From pieces to patterns: evolutionary engineering in bacterial pathogens. Nat Rev Microbiol 2:510–518.

19. Benler S, Faure G, Altae-Tran H, Shmakov S, Zhang F, Koonin E. 2021. Cargo Genes of Tn7-Like Transposons Comprise an Enormous Diversity of Defense Systems, Mobile Genetic Elements, and Antibiotic Resistance Genes. mBio 12:e02938–21.

20. Gluck-Thaler E, Ralston T, Konkel Z, Ocampos CG, Ganeshan VD, Dorrance AE, Niblack TL, Wood CW, Slot JC, Lopez-Nicora HD, Vogan AA. 2022. Giant Starship Elements Mobilize Accessory Genes in Fungal Genomes. Mol Biol Evol 39:msac109.

21. Urquhart AS, Vogan AA, Gardiner DM, Idnurm A. 2023. Starships are active eukaryotic transposable elements mobilized by a new family of tyrosine recombinases. Proc Natl Acad Sci U S A 120:e2214521120.

22. Gluck-Thaler E, Vogan AA. 2024. Systematic identification of cargo-mobilizing genetic elements reveals new dimensions of eukaryotic diversity. Nucleic Acids Res gkae327.

23. Urquhart A, Vogan AA, Gluck-Thaler E. 2024. Starships: a new frontier for fungal biology. Trends Genet 40:1060–1073.

24. Gourlie R, McDonald M, Hafez M, Ortega-Polo R, Low KE, Abbott DW, Strelkov SE, Daayf F, Aboukhaddour R. 2022. The pangenome of the wheat pathogen Pyrenophora tritici-repentis reveals novel transposons associated with necrotrophic effectors ToxA and ToxB. BMC Biol 20:239.

25. Urquhart AS, Chong NF, Yang Y, Idnurm A. 2022. A large transposable element mediates metal resistance in the fungus Paecilomyces variotii. Curr Biol 32:937–950.e5.

26. Urquhart AS, Gluck-Thaler E, Vogan AA. 2023. Gene acquisition by giant transposons primes eukaryotes for rapid evolution via horizontal gene transfer. bioRxiv 10.1101/2023.11.22.568313.

27. Bucknell A, Wilson HM, Santos KCG do, Simpfendorfer S, Milgate A, Germain H, Solomon PS, Bentham A, McDonald MC. 2024. Sanctuary: A Starship transposon facilitating the movement of the virulence factor ToxA in fungal wheat pathogens. bioRxiv 10.1101/2024.03.04.583430.

28. Urquhart AS, O’Donnell S, Gluck-Thaler E, Vogan AA. 2025. A natural mechanism of eukaryotic horizontal gene transfer. bioRxiv 10.1101/2025.02.28.640899.

29. Pain A, Woodward J, Quail MA, Anderson MJ, Clark R, Collins M, Fosker N, Fraser A, Harris D, Larke N, Murphy L, Humphray S, O’Neil S, Pertea M, Price C, Rabbinowitsch E, Rajandream M-A, Salzberg S, Saunders D, Seeger K, Sharp S, Warren T, Denning DW, Barrell B, Hall N. 2004. Insight into the genome of Aspergillus fumigatus: analysis of a 922 kb region encompassing the nitrate assimilation gene cluster. Fungal Genet Biol FG B 41:443–453.

30. Monod M, Paris S, Sarfati J, Jaton-Ogay K, Ave P, Latgé JP. 1993. Virulence of alkaline protease-deficient mutants of Aspergillus fumigatus. FEMS Microbiol Lett 106:39–46.

31. Bowyer P, Currin A, Delneri D, Fraczek MG. 2022. Telomere-to-telomere genome sequence of the model mould pathogen Aspergillus fumigatus. Nat Commun 13:5394.

32. Bertuzzi M, van Rhijn N, Krappmann S, Bowyer P, Bromley MJ, Bignell EM. 2021. On the lineage of Aspergillus fumigatus isolates in common laboratory use. Med Mycol 59:7–13.

33. Kowalski CH, Morelli KA, Stajich JE, Nadell CD, Cramer RA. 2021. A Heterogeneously Expressed Gene Family Modulates the Biofilm Architecture and Hypoxic Growth of Aspergillus fumigatus. mBio 12:e03579–20.

34. Wicker T, Sabot F, Hua-Van A, Bennetzen JL, Capy P, Chalhoub B, Flavell A, Leroy P, Morgante M, Panaud O, Paux E, SanMiguel P, Schulman AH. 2007. A unified classification system for eukaryotic transposable elements. Nat Rev Genet 8:973–982.

35. Lambert SA, Yang AWH, Sasse A, Cowley G, Albu M, Caddick MX, Morris QD, Weirauch MT, Hughes TR. 2019. Similarity regression predicts evolution of transcription factor sequence specificity. Nat Genet 51:981–989.

36. Lind AL, Wisecaver JH, Lameiras C, Wiemann P, Palmer JM, Keller NP, Rodrigues F, Goldman GH, Rokas A. 2017. Drivers of genetic diversity in secondary metabolic gene clusters within a fungal species. PLoS Biol 15:e2003583.

37. Abdolrasouli A, Rhodes J, Beale MA, Hagen F, Rogers TR, Chowdhary A, Meis JF, Armstrong-James D, Fisher MC. 2015. Genomic Context of Azole Resistance Mutations in Aspergillus fumigatus Determined Using Whole-Genome Sequencing. mBio 6:e00536–15.

38. Ma K, Zhang P, Tao Q, Keller NP, Yang Y, Yin W-B, Liu H. 2019. Characterization and Biosynthesis of a Rare Fungal Hopane-Type Triterpenoid Glycoside Involved in the Antistress Property of Aspergillus fumigatus. Org Lett 21:3252–3256.

39. Brown NA, Goldman GH. 2016. The contribution of Aspergillus fumigatus stress responses to virulence and antifungal resistance. J Microbiol 54:243–253.

40. Steenwyk JL, Mead ME, de Castro PA, Valero C, Damasio A, dos Santos RAC, Labella AL, Li Y, Knowles SL, Raja HA, Oberlies NH, Zhou X, Cornely OA, Fuchs F, Koehler P, Goldman GH, Rokas A. 2021. Genomic and Phenotypic Analysis of COVID-19-Associated Pulmonary Aspergillosis Isolates of Aspergillus fumigatus. Microbiol Spectr 9:10.1128/spectrum.00010-21.

41. Raffa N, Osherov N, Keller NP. 2019. Copper Utilization, Regulation, and Acquisition by Aspergillus fumigatus. 8. Int J Mol Sci 20:1980.

42. Bignell E, Cairns TC, Throckmorton K, Nierman WC, Keller NP. 2016. Secondary metabolite arsenal of an opportunistic pathogenic fungus. Philos Trans R Soc B Biol Sci 371:20160023.

43. Abad A, Fernández-Molina JV, Bikandi J, Ramírez A, Margareto J, Sendino J, Hernando FL, Pontón J, Garaizar J, Rementeria A. 2010. What makes Aspergillus fumigatus a successful pathogen? Genes and molecules involved in invasive aspergillosis. Rev Iberoam Micol 27:155–182.

44. Auxier B, Zhang J, Marquez FR, Senden K, Heuvel J van den, Aanen DK, Snelders E, Debets AJM. 2022. Identification of heterokaryon incompatibility genes in Aspergillus fumigatus highlights a narrow footprint of ancient balancing selection. bioRxiv 10.1101/2022.11.25.517501.

45. Stroe MC, Netzker T, Scherlach K, Krüger T, Hertweck C, Valiante V, Brakhage AA. 2020. Targeted induction of a silent fungal gene cluster encoding the bacteria-specific germination inhibitor fumigermin. eLife 9:e52541.

46. Kowalski CH, Kerkaert JD, Liu K-W, Bond MC, Hartmann R, Nadell CD, Stajich JE, Cramer RA. 2019. Fungal biofilm morphology impacts hypoxia fitness and disease progression. Nat Microbiol 4:2430–2441.

47. O’Donnell S, Rezende G, Vernadet J-P, Snirc A, Ropars J. 2024. Harbouring Starships: The accumulation of large Horizontal Gene Transfers in Domesticated and Pathogenic Fungi. bioRxiv 10.1101/2024.07.03.601904.

48. Rokas A, Mead ME, Steenwyk JL, Oberlies NH, Goldman GH. 2020. Evolving moldy murderers: Aspergillus section Fumigati as a model for studying the repeated evolution of fungal pathogenicity. PLOS Pathog 16:e1008315.

49. Shaw LP, Chau KK, Kavanagh J, AbuOun M, Stubberfield E, Gweon HS, Barker L, Rodger G, Bowes MJ, Hubbard ATM, Pickford H, Swann J, Gilson D, Smith RP, Hoosdally SJ, Sebra R, Brett H, Peto TEA, Bailey MJ, Crook DW, Read DS, Anjum MF, Walker AS, Stoesser N, REHAB CONSORTIUM. 2021. Niche and local geography shape the pangenome of wastewater- and livestock-associated Enterobacteriaceae. Sci Adv 7:eabe3868.

50. Shimizu K, Keller NP. 2001. Genetic involvement of a cAMP-dependent protein kinase in a G protein signaling pathway regulating morphological and chemical transitions in Aspergillus nidulans. Genetics 157:591–600.

51. Carter-House D, Stajich J, Unruh S, Kurbessoian T. 2020. Fungal CTAB DNA Extraction.

52. Koren S, Walenz BP, Berlin K, Miller JR, Bergman NH, Phillippy AM. 2017. Canu: scalable and accurate long-read assembly via adaptive k-mer weighting and repeat separation. Genome Res 27:722–736.

53. Walker BJ, Abeel T, Shea T, Priest M, Abouelliel A, Sakthikumar S, Cuomo CA, Zeng Q, Wortman J, Young SK, Earl AM. 2014. Pilon: an integrated tool for comprehensive microbial variant detection and genome assembly improvement. PloS One 9:e112963.

54. Palmer JM, Stajich JE. 2022. Automatic assembly for the fungi (AAFTF): genome assembly pipeline (v0.4.1). Zenodo.

55. Manni M, Berkeley MR, Seppey M, Simão FA, Zdobnov EM. 2021. BUSCO Update: Novel and Streamlined Workflows along with Broader and Deeper Phylogenetic Coverage for Scoring of Eukaryotic, Prokaryotic, and Viral Genomes. Mol Biol Evol 38:4647–4654.

56. Palmer JM, Stajich J. 2020. Funannotate v1.8.1: Eukaryotic genome annotation (v1.8.1). Zenodo.

57. Haas BJ, Delcher AL, Mount SM, Wortman JR, Smith RK, Hannick LI, Maiti R, Ronning CM, Rusch DB, Town CD, Salzberg SL, White O. 2003. Improving the Arabidopsis genome annotation using maximal transcript alignment assemblies. Nucleic Acids Res 31:5654–5666.

58. Jones P, Binns D, Chang H-Y, Fraser M, Li W, McAnulla C, McWilliam H, Maslen J, Mitchell A, Nuka G, Pesseat S, Quinn AF, Sangrador-Vegas A, Scheremetjew M, Yong S- Y, Lopez R, Hunter S. 2014. InterProScan 5: genome-scale protein function classification. Bioinformatics 30:1236–1240.

59. Cantalapiedra CP, Hernández-Plaza A, Letunic I, Bork P, Huerta-Cepas J. 2021. eggNOG-mapper v2: Functional Annotation, Orthology Assignments, and Domain Prediction at the Metagenomic Scale. Mol Biol Evol 10.1093/molbev/msab293.

60. Zheng J, Ge Q, Yan Y, Zhang X, Huang L, Yin Y. 2023. dbCAN3: automated carbohydrate-active enzyme and substrate annotation. Nucleic Acids Res 51:W115–W121.

61. Blin K, Shaw S, Kloosterman AM, Charlop-Powers Z, van Wezel GP, Medema MH, Weber T. 2021. antiSMASH 6.0: improving cluster detection and comparison capabilities. Nucleic Acids Res 49:W29–W35.

62. Nawrocki EP, Eddy SR. 2013. Infernal 1.1: 100-fold faster RNA homology searches. Bioinformatics 29:2933–2935.

63. Purcell S, Neale B, Todd-Brown K, Thomas L, Ferreira MAR, Bender D, Maller J, Sklar P, de Bakker PIW, Daly MJ, Sham PC. 2007. PLINK: A Tool Set for Whole-Genome Association and Population-Based Linkage Analyses. Am J Hum Genet 81:559–575.

64. Lofgren LA, Ross B, Cramer Jr RA, Stajich JE. 2021. Aspergillus fumigatus pangenome dataset (v2021-12-12). Zenodo.

65. Emms DM, Kelly S. 2015. OrthoFinder: solving fundamental biases in whole genome comparisons dramatically improves orthogroup inference accuracy. Genome Biol 16:157.

66. Camacho C, Coulouris G, Avagyan V, Ma N, Papadopoulos J, Bealer K, Madden TL. 2009. BLAST+: architecture and applications. BMC Bioinformatics 10:421.

67. Vasimuddin Md, Misra S, Li H, Aluru S. 2019. Efficient Architecture-Aware Acceleration of BWA-MEM for Multicore Systems, p. 314–324. In 2019 IEEE International Parallel and Distributed Processing Symposium (IPDPS).

68. Prjibelski A, Antipov D, Meleshko D, Lapidus A, Korobeynikov A. 2020. Using SPAdes De Novo Assembler. Curr Protoc Bioinforma 70:e102.

69. Shumate A, Salzberg SL. 2021. Liftoff: accurate mapping of gene annotations. Bioinformatics 37:1639–1643.

70. Marçais G, Delcher AL, Phillippy AM, Coston R, Salzberg SL, Zimin A. 2018. MUMmer4: A fast and versatile genome alignment system. PLOS Comput Biol 14:e1005944.

71. Levy Karin E, Mirdita M, Söding J. 2020. MetaEuk—sensitive, high-throughput gene discovery, and annotation for large-scale eukaryotic metagenomics. Microbiome 8:48.

72. Ayad LAK, Pissis SP, Polychronopoulos D. 2018. CNEFinder: finding conserved non-coding elements in genomes. Bioinforma Oxf Engl 34:i743–i747.

73. Katoh K, Standley DM. 2013. MAFFT Multiple Sequence Alignment Software Version 7: Improvements in Performance and Usability. Mol Biol Evol 30:772–780.

74. Hauser M, Steinegger M, Söding J. 2016. MMseqs software suite for fast and deep clustering and searching of large protein sequence sets. Bioinformatics 32:1323–1330.

75. Brown CT, Irber L. 2016. sourmash: a library for MinHash sketching of DNA. J Open Source Softw 1:27.

76. Enright AJ, Van Dongen S, Ouzounis CA. 2002. An efficient algorithm for large-scale detection of protein families. Nucleic Acids Res 30:1575–1584.

77. Sahlin K. 2022. Strobealign: flexible seed size enables ultra-fast and accurate read alignment. Genome Biol 23:260.

78. Patro R, Duggal G, Love MI, Irizarry RA, Kingsford C. 2017. Salmon provides fast and bias-aware quantification of transcript expression. Nat Methods 14:417–419.

79. Dobin A, Davis CA, Schlesinger F, Drenkow J, Zaleski C, Jha S, Batut P, Chaisson M, Gingeras TR. 2013. STAR: ultrafast universal RNA-seq aligner. Bioinformatics 29:15–21.

80. Zhang Y, Parmigiani G, Johnson WE. 2020. ComBat-seq: batch effect adjustment for RNA-seq count data. NAR Genomics Bioinforma 2:lqaa078.

81. Love MI, Huber W, Anders S. 2014. Moderated estimation of fold change and dispersion for RNA-seq data with DESeq2. Genome Biol 15:550.

82. MetaVolcanoR. Bioconductor. http://bioconductor.org/packages/MetaVolcanoR/. Retrieved 26 June 2024.

83. Rezaie N, Reese F, Mortazavi A. 2023. PyWGCNA: a Python package for weighted gene co-expression network analysis. Bioinformatics 39:btad415.

84. Virtanen P, Gommers R, Oliphant TE, Haberland M, Reddy T, Cournapeau D, Burovski E, Peterson P, Weckesser W, Bright J, van der Walt SJ, Brett M, Wilson J, Millman KJ, Mayorov N, Nelson ARJ, Jones E, Kern R, Larson E, Carey CJ, Polat İ, Feng Y, Moore EW, VanderPlas J, Laxalde D, Perktold J, Cimrman R, Henriksen I, Quintero EA, Harris CR, Archibald AM, Ribeiro AH, Pedregosa F, van Mulbregt P. 2020. SciPy 1.0: fundamental algorithms for scientific computing in Python. Nat Methods 17:261–272.

85. Kolberg L, Raudvere U, Kuzmin I, Vilo J, Peterson H. 2020. gprofiler2 -- an R package for gene list functional enrichment analysis and namespace conversion toolset g:Profiler. F1000Research 9:ELIXIR-709.

86. Krzywinski MI, Schein JE, Birol I, Connors J, Gascoyne R, Horsman D, Jones SJ, Marra MA. 2009. Circos: An information aesthetic for comparative genomics. Genome Res 10.1101/gr.092759.109.

87. Hackl T, Ankenbrand M, van Adrichem B. 2023. gggenomes: A Grammar of Graphics for Comparative Genomics (R package version 0.9.9.9000).

88. Wickham H. 2016. ggplot2: Elegant Graphics for Data Analysis, 3rd ed. Springer International Publishing, Cham. http://link.springer.com/10.1007/978-3-319-24277-4. Retrieved 10 October 2023.

89. McInnes L, Healy J, Saul N, Großberger L. 2018. UMAP: Uniform Manifold Approximation and Projection. J Open Source Softw 3:861.

