## Supplementary material for "Giant transposons promote strain heterogeneity in a major fungal pathogen": Table 1

| **Family** | **Name** | **Predicted target site** | **Mean length in bp (SD)** | **Type element** | **Functions of interest in type element** | **Ref. strains** |
|---|---|---|---|---|---|---|
| Tardis | *Tardis h1* | TACGGAGTAG | 81,743 (9,347) | W72310-lr_s00161 | GH71, SAH | Af293 |
| Prometheus | *Gnosis h1* | A($N_3$)CTA($N_{17}$)T | 82,313 (23,570) | A-fum-AFUG-100413-0667_s00261 | GH18, lysM | Af293, CEA10 |
| Prometheus | *Gnosis h2* | - | 93,292 (60) | CM2733_s04518 | bafY, fumihopaside A BGC | CEA10 |
| Phoenix | *Janus h1* | - | 62,365 (19,076) | F14513G_s04800 | CHROMO | - |
| Phoenix | *Janus h2* | A($N_3$)TTACTA ($N_2$)A($N_{15}$)CT | 217,821 (102,982) | 47-4_s00005 | CHROMO, DHDPS, GH18, GH71 | CEA10 |
| Phoenix | *Janus h3* | A($N_3$)TTACTA ($N_2$)A($N_{15}$)CT | 42,070 (37,412) | SF1S6_s05434 | CHROMO | - |
| Phoenix | *Janus h4* | - | 19,097 (5,047) | CM6458_s04616 | CHROMO | - |
| Phoenix | *Janus h5* | - | 34,546 (18,524) | 47-10_s00011 | CHROMO | - |
| Enterprise | *Osiris h1* | 5S rDNA | 37,633 (2,148) | A-fum-AFIS-13708-CDC-14_s00209 | putative BGC | - |
| Enterprise | *Osiris h2* | 5S rDNA | 138,260 (2,424) | B5269_s04148 | ARS, B-LAC, COP | - |
| Enterprise | *Osiris h3* | 5S rDNA | 70,476 (28,148) | Afu-218-E11_s00395 | bafB, hrmB | CEA10 |
| Enterprise | *Osiris h4* | 5S rDNA | 51,342 (13,227) | Af293_s00032 | putative BGC | Af293 |
| Galactica | *Lamia h1* | A(N)TAGT | 69,899 (24,972) | S02-30_s00148 | GH18, lysM | - |
| Galactica | *Lamia h2* | A(N)TAGT | 65,414 (6,136) | ATCC42202_s00060 | fumigermin BGC | - |
| Galactica | *Lamia h3* | A(N)TAGT | 236,680 (74,695) | ISSF-21_s05253 | GH18, GH71 | - |
| Galactica | *Lamia h4* | A(N)TAGT | 291,189 (215,918) | F7763_s04695 | bafX, fumigermin BGC, GH3, GH18, GH71, GT8, GT31 | Af293 |
| Prometheus | *Logos h1* | A($N_{10}$)TACTTAT TA(N)A($N_6$)A | 69,008 (5,313) | CEA10_s00024 | bafZ | CEA10 |
| Prometheus | *Logos h2* | - | 56,189 (31,755) | CEA10_s00031 | bafC | CEA10 |
| Hephaestus | *Neb h1* | TTACA($N_5$)AAT | 48,346 (5,585) | AF293_s00037 | ARS, bafA, cgnA, hrmA | Af293, CEA10 |
| Hephaestus | *Neb h2* | TTACA($N_5$)AAT | 54,751 (7,226) | AF293_s00031 | ARS | Af293 |

Table 1: High Confidence *Starships* in *Aspergillus fumigatus*. Abbreviations: GH: glycosyl hydrolase; GT: glycosyl transferase; SAH: salicylate hydroxylase; DHDPS: dihydrodipicolinate synthetase; CHROMO: chromatin modification domain; ARS: arsenic resistance cluster; COP: copper resistance; B-LAC: Beta-lactamase
