## Supplemental Figures S1 to S16 for "Giant transposons promote strain heterogeneity in a major fungal pathogen"

Figure S16: Heatmap displaying the correlation values for eigengenes in each module and their association with samples from a specific study (labeled with BioProject IDs), or different levels of experimental treatment category across studies. Cells of the heatmap are coloured by the correlation value, as well as labeled in each cell. P-values from a pairwise comparison between observations (*pdist*) are shown in parentheses.

Figure S1

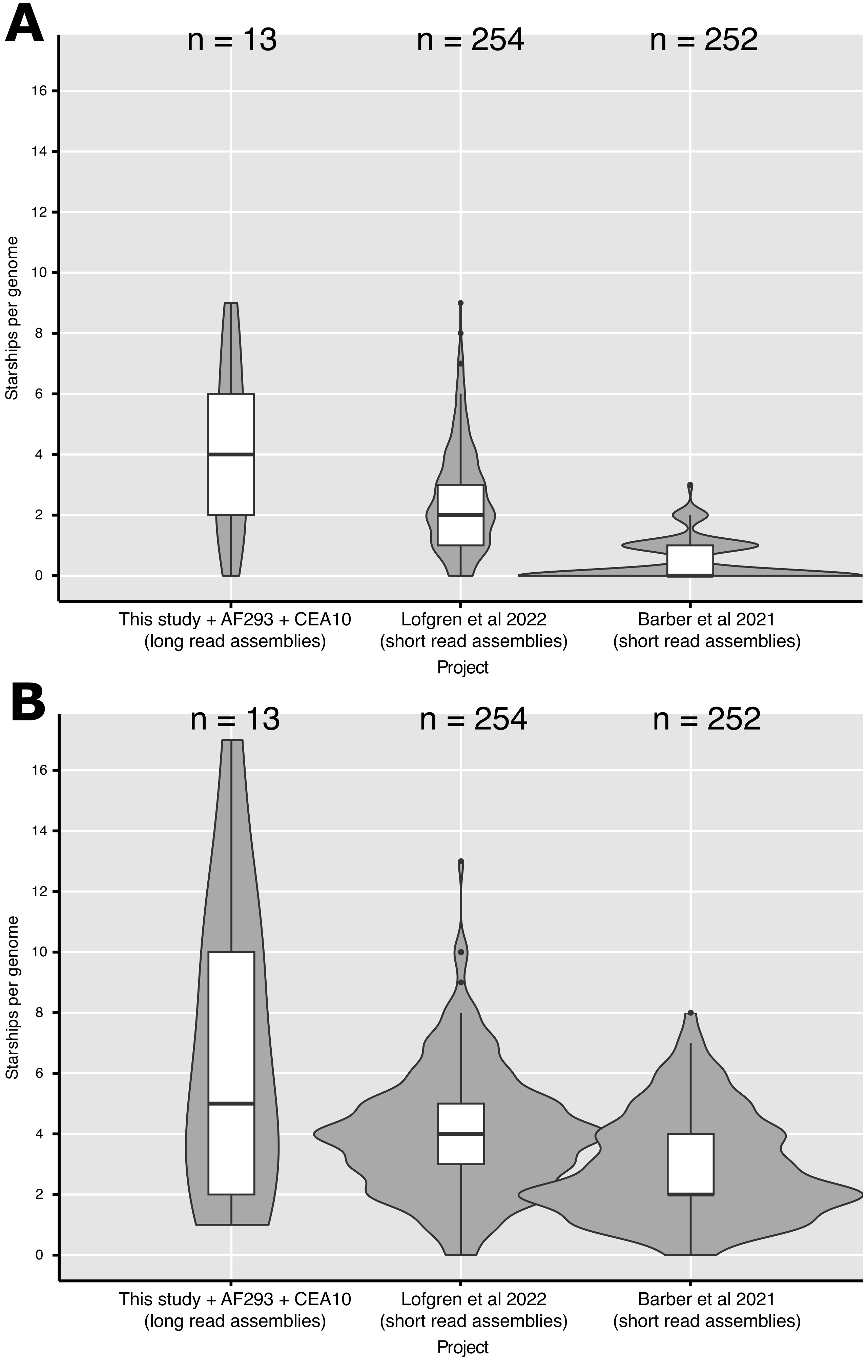
A
n = 13
n = 254
n = 252
Starships per genome
0
2
4
6
8
10
12
14
16
This study + AF293 + CEA10
(long read assemblies)
Lofgren et al 2022
(short read assemblies)
Barber et al 2021
(short read assemblies)
Project
B
n = 13
n = 254
n = 252
Starships per genome
0
2
4
6
8
10
12
14
16
This study + AF293 + CEA10
(long read assemblies)
Lofgren et al 2022
(short read assemblies)
Barber et al 2021
(short read assemblies)
Project

### Figure S2A

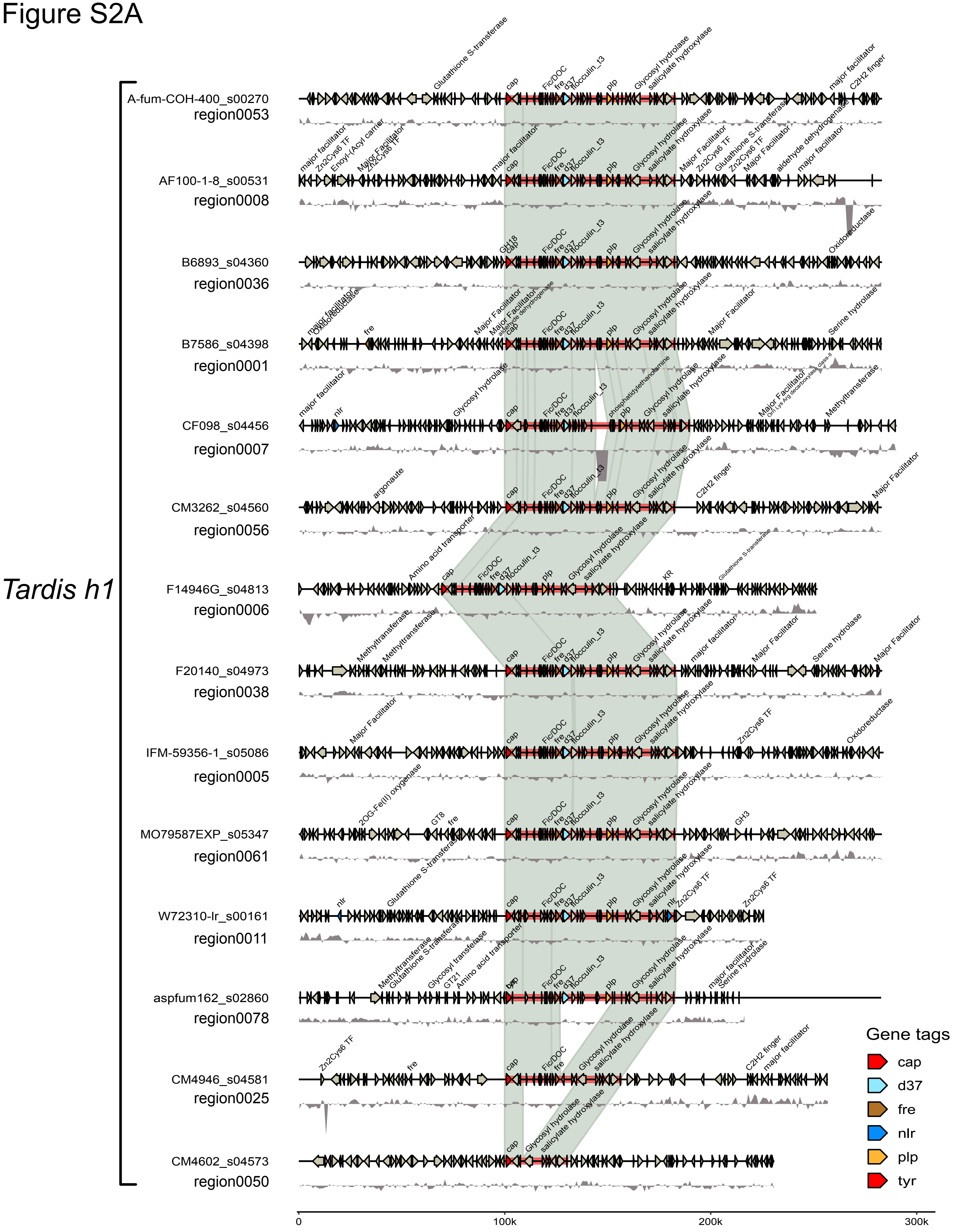
Tardis h1
A-fum-COH-400_s00270
region0053
AF100-1-8_s00531
region0008
B6893_s04360
region0036
B7586_s04398
region0001
CF098_s04456
region0007
CM3262_s04560
region0056
F14946G_s04813
region0006
F20140_s04973
region0038
IFM-59356-1_s05086
region0005
MO79587EXP_s05347
region0061
W72310-Ir_s00161
region0011
aspfum162_s02860
region0078
CM4946_s04581
region0025
CM4602_s04573
region0050
0
100k
200k
300k
Gene tags
cap
d37
fre
nlr
plp
tyr

Figure S2B

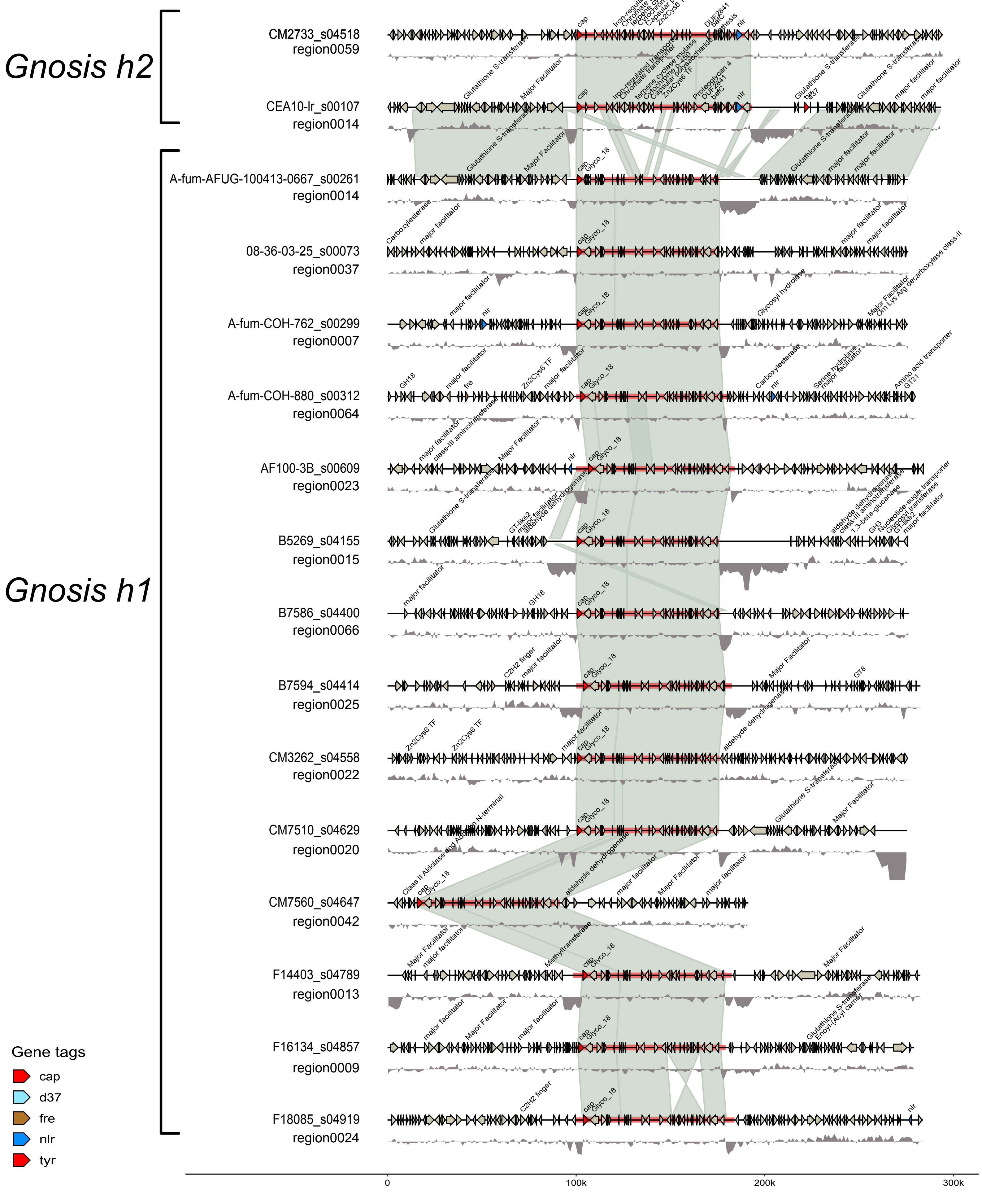
Gnosis h2
Gnosis h1
CM2733_s04518
region0059
CEA10-lr_s00107
region0014
A-fum-AFUG-100413-0667_s00261
region0014
08-36-03-25_s00073
region0037
A-fum-COH-762_s00299
region0007
A-fum-COH-880_s00312
region0064
AF100-3B_s00609
region0023
B5269_s04155
region0015
B7586_s04400
region0066
B7594_s04414
region0025
CM3262_s04558
region0022
CM7510_s04629
region0020
CM7560_s04647
region0042
F14403_s04789
region0013
F16134_s04857
region0009
F18085_s04919
region0024
Gene tags
cap
d37
fre
nlr
tyr
0
100k
200k
300k

Figure S2C

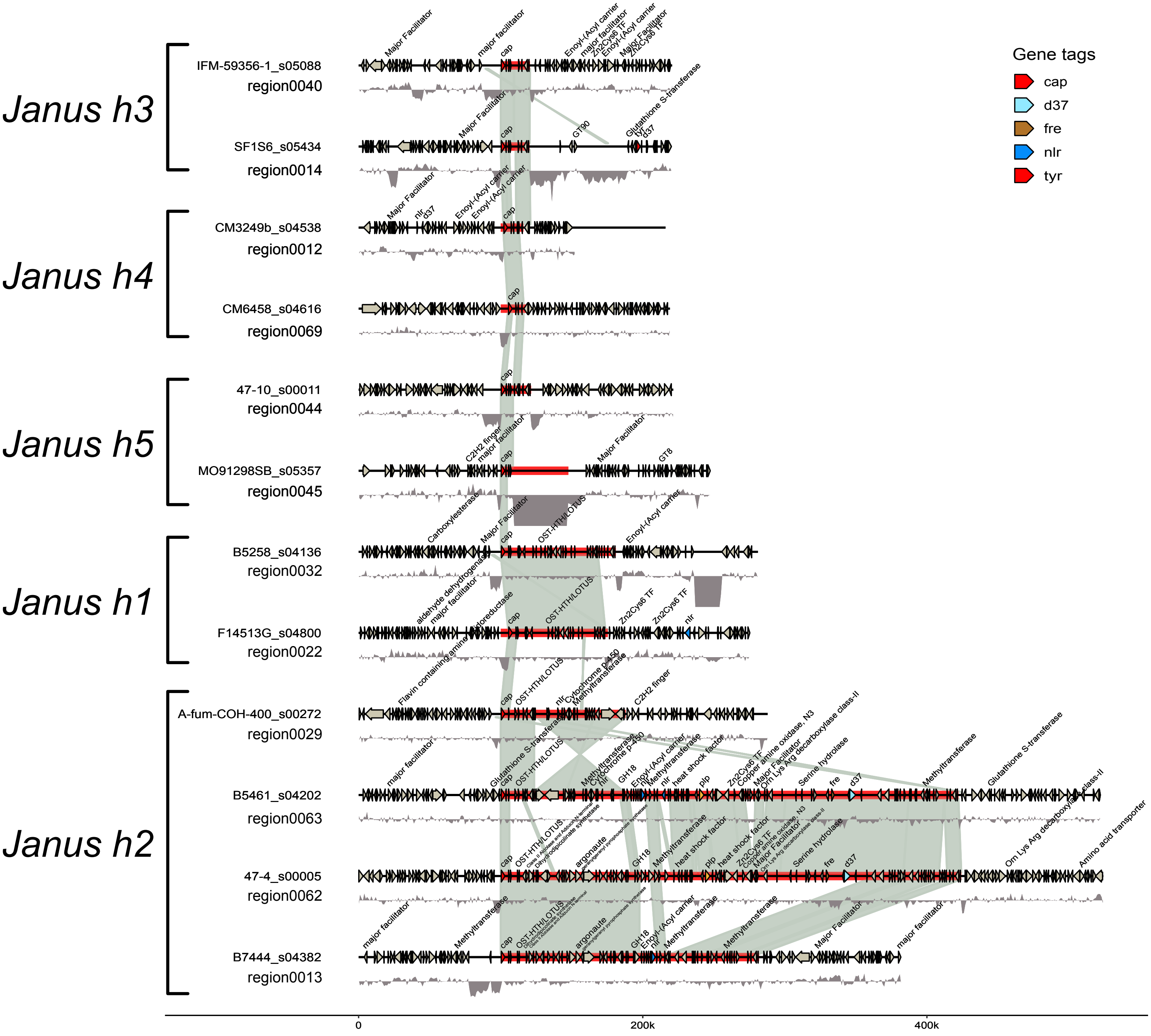
Janus h3
IFM-59356-1_s05088
region0040
SF1S6_s05434
region0014
Janus h4
CM3249b_s04538
region0012
CM6458_s04616
region0069
Janus h5
47-10_s00011
region0044
MO91298SB_s05357
region0045
Janus h1
B5258_s04136
region0032
F14513G_s04800
region0022
Janus h2
A-fum-COH-400_s00272
region0029
B5461_s04202
region0063
47-4_s00005
region0062
B7444_s04382
region0013
Gene tags
cap
d37
fre
nlr
tyr
0
200k
400k

### Figure S2D

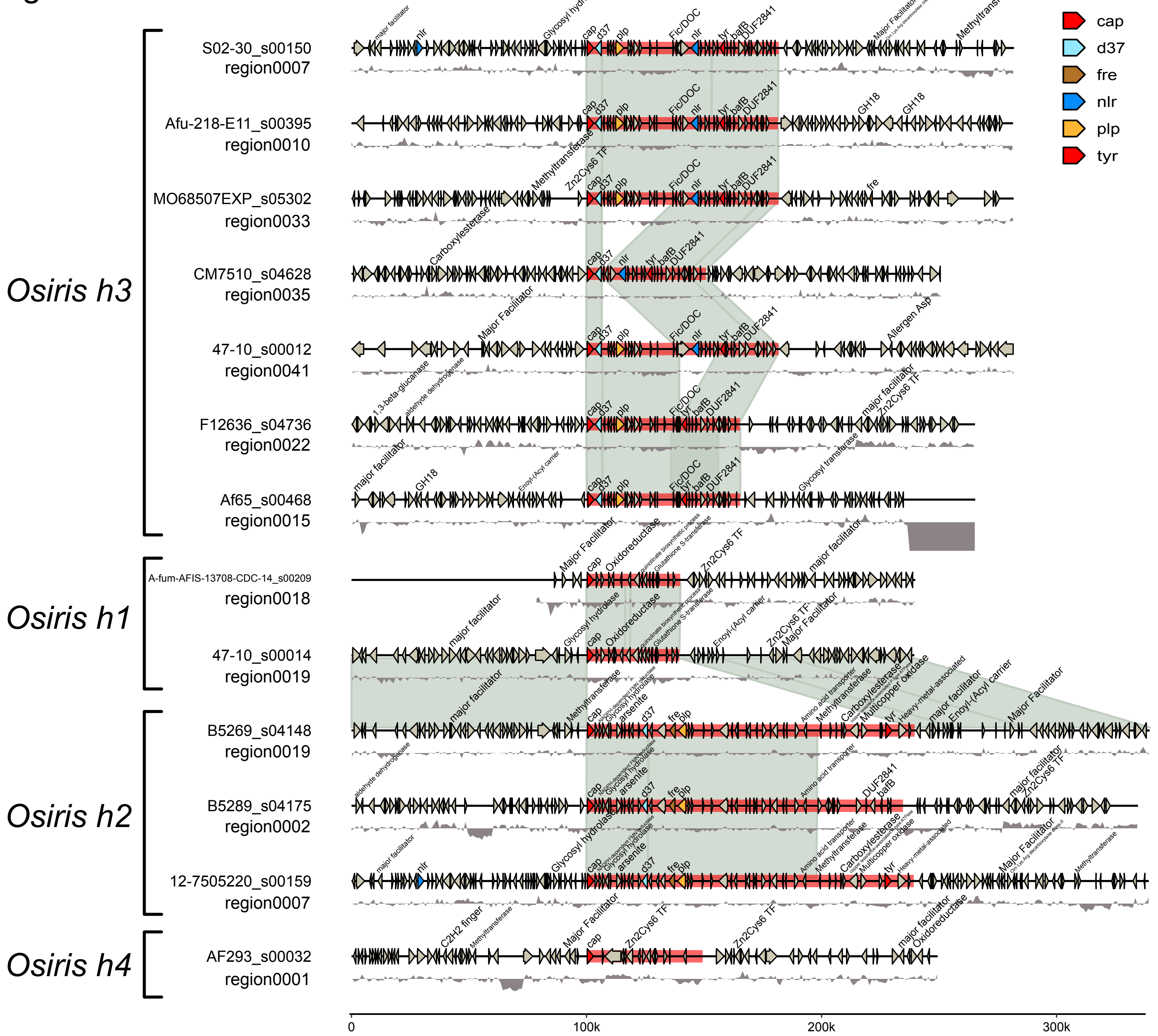

Gene tags
cap
d37
fre
nlr
plp
tyr
Osiris h3
S02-30_s00150
region0007
Afu-218-E11_s00395
region0010
MO68507EXP_s05302
region0033
CM7510_s04628
region0035
47-10_s00012
region0041
F12636_s04736
region0022
Af65_s00468
region0015
Osiris h1
A-fum-AFIS-13708-CDC-14_s00209
region0018
47-10_s00014
region0019
Osiris h2
B5269_s04148
region0019
B5289_s04175
region0002
12-7505220_s00159
region0007
Osiris h4
AF293_s00032
region0001
0
100k
200k
300k

### Figure S2E

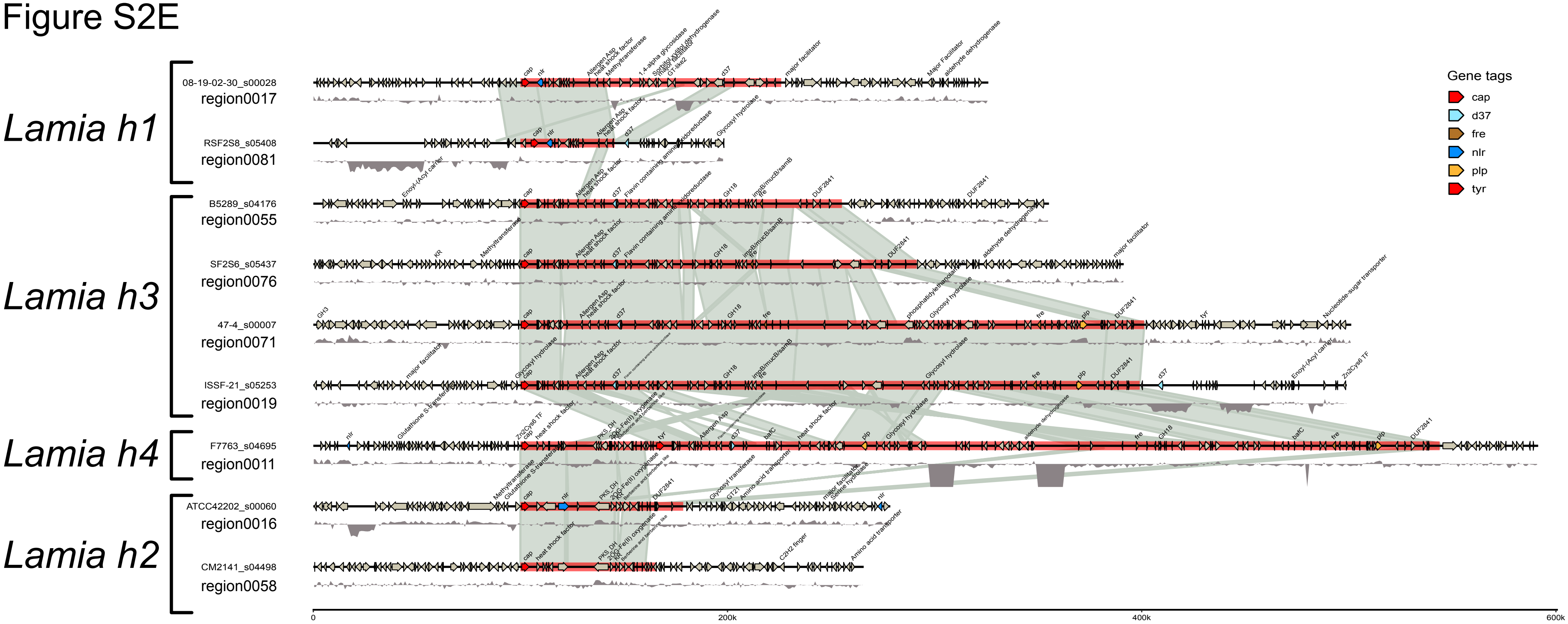

Lamia h1
08-19-02-30_s00028
region0017
RSF2S8_s05408
region0081
Lamia h3
B5289_s04176
region0055
SF2S6_s05437
region0076
47-4_s00007
region0071
ISSF-21_s05253
region0019
Lamia h4
F7763_s04695
region0011
Lamia h2
ATCC42202_s00060
region0016
CM2141_s04498
region0058
0
200k
400k
600k
Gene tags
cap
d37
fre
nlr
plp
tyr

### Figure S2F

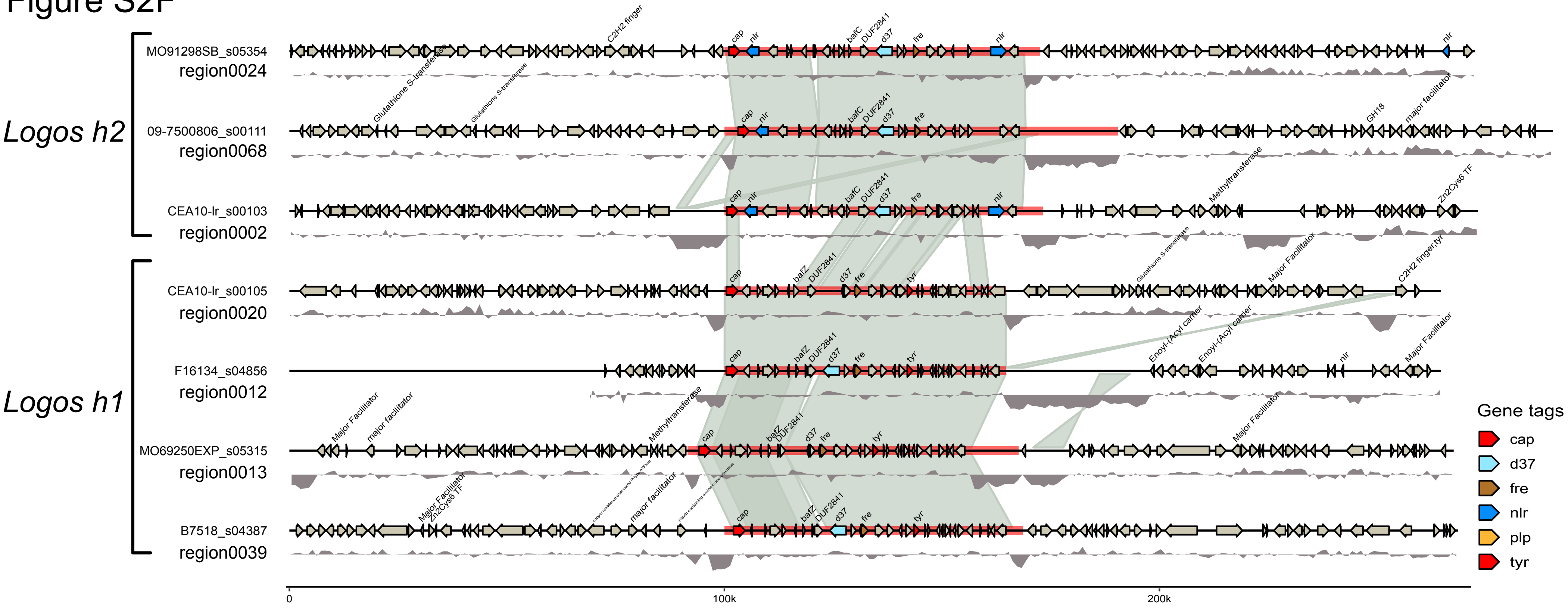

Logos h2
Logos h1
MO91298SB_s05354
region0024
09-7500806_s00111
region0068
CEA10-lr_s00103
region0002
CEA10-lr_s00105
region0020
F16134_s04856
region0012
MO69250EXP_s05315
region0013
B7518_s04387
region0039
cap
nlr
bafC
bafZ
DUF2841
d37
fre
tyr
C2H2 finger
Glutathione S-transferase
GH18
major facilitator
Major Facilitator
Methyltransferase
Zn2Cys6 TF
Enoyl-(Acyl carrier
C2H2 finger,tyr
Gene tags
cap
d37
fre
nlr
plp
tyr
0
100k
200k

### Figure S2G

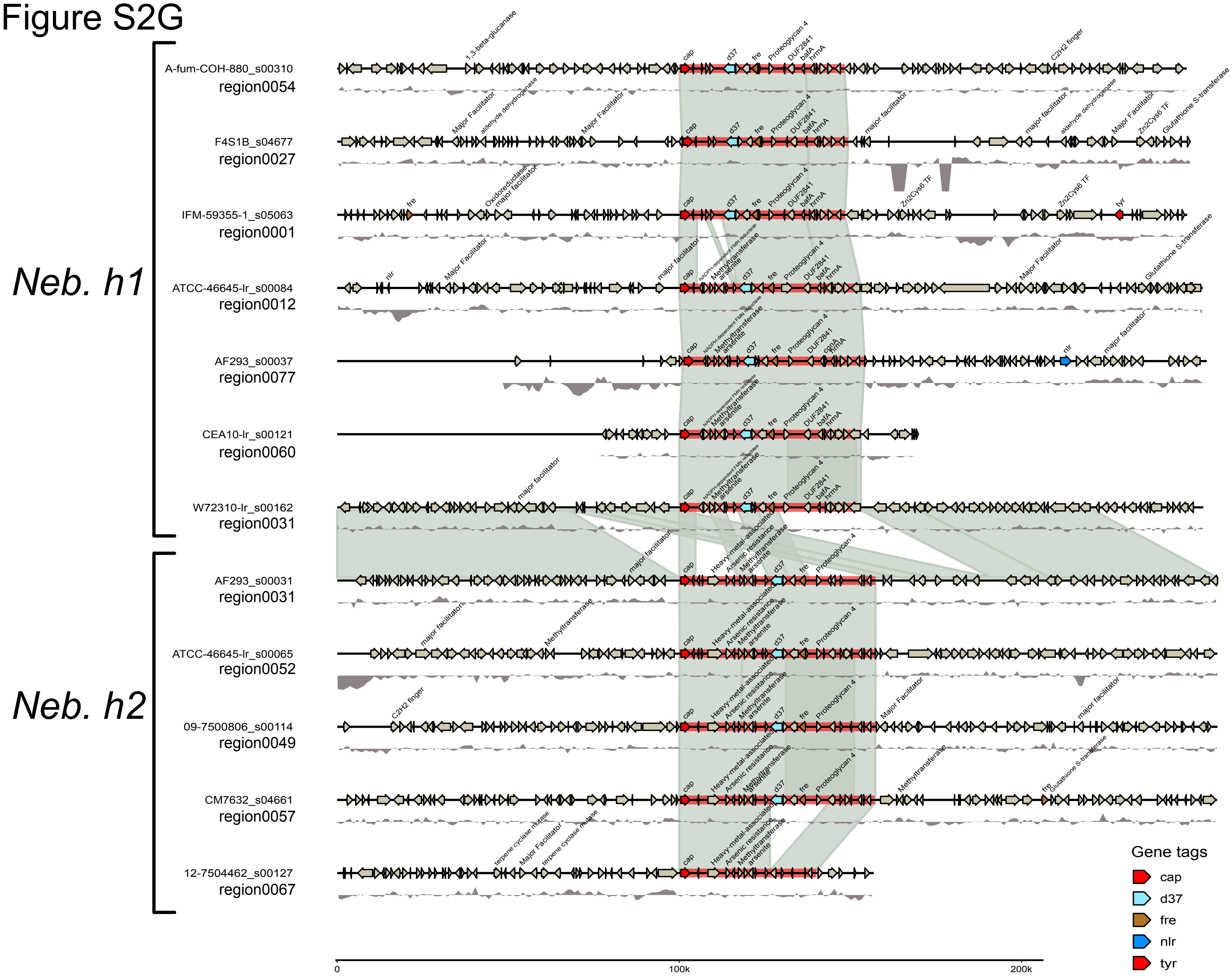
Neb. h1
Neb. h2
A-fum-COH-880_s00310
region0054
F4S1B_s04677
region0027
IFM-59355-1_s05063
region0001
ATCC-46645-lr_s00084
region0012
AF293_s00037
region0077
CEA10-lr_s00121
region0060
W72310-lr_s00162
region0031
AF293_s00031
region0031
ATCC-46645-lr_s00065
region0052
09-7500806_s00114
region0049
CM7632_s04661
region0057
12-7504462_s00127
region0067
Gene tags
cap
d37
fre
nlr
tyr
0
100k
200k

Figure S3

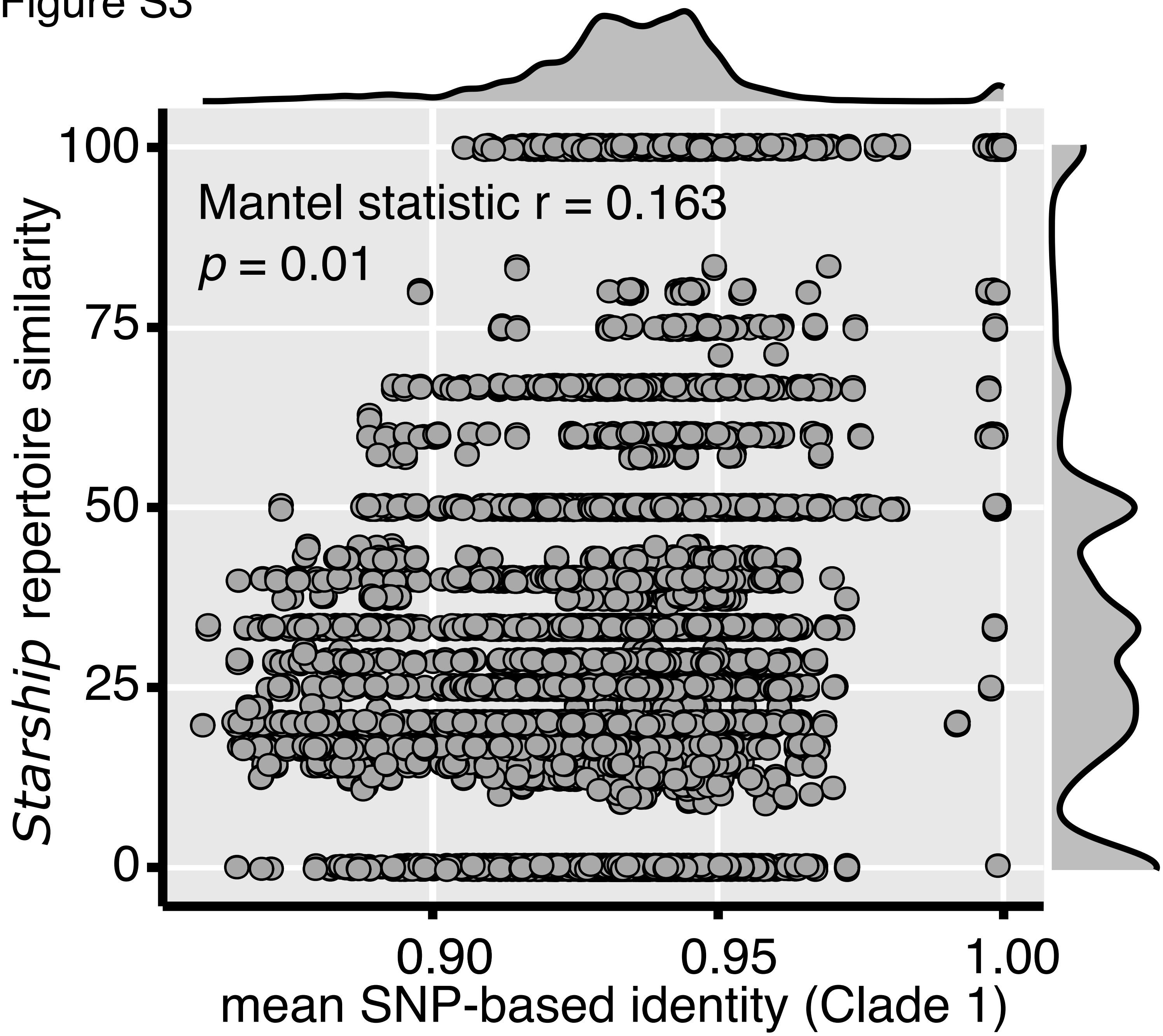
Mantel statistic r = 0.163
p = 0.01
Starship repertoire similarity
100
75
50
25
0
0.90
0.95
1.00
mean SNP-based identity (Clade 1)

### Figure S4

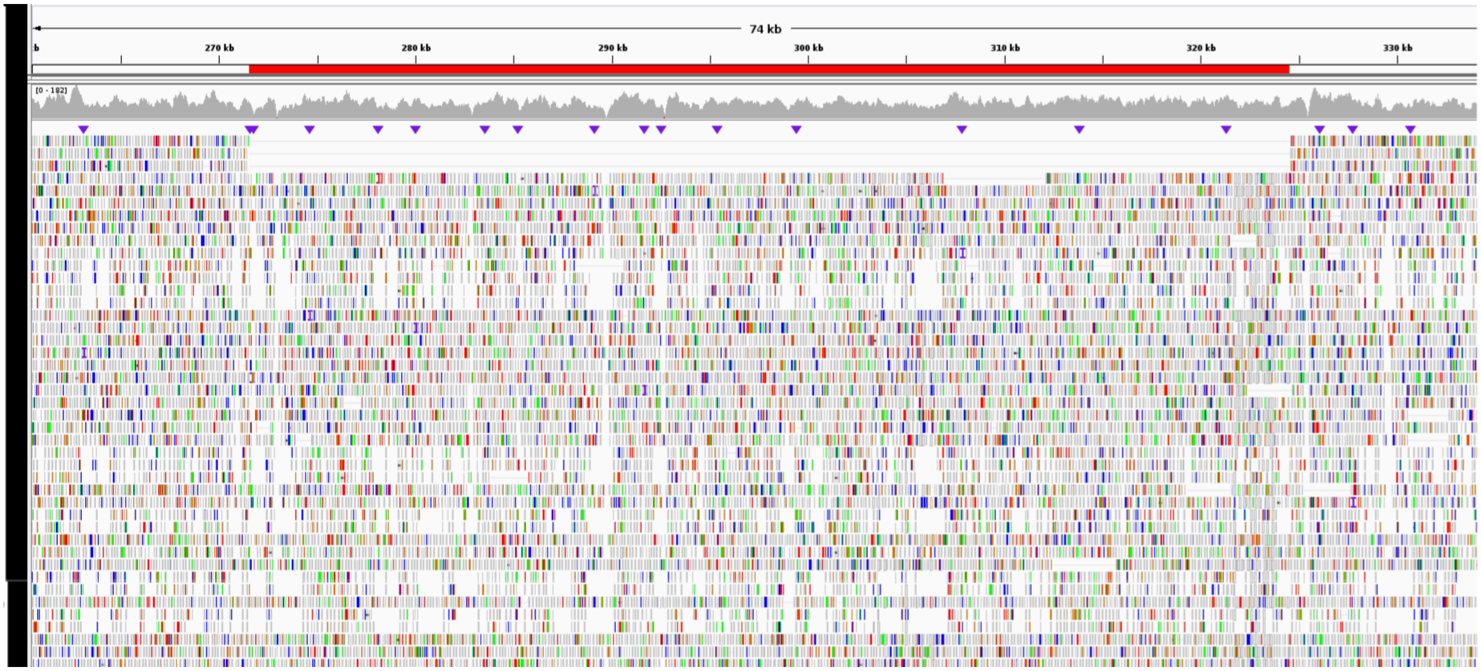

Figure S5

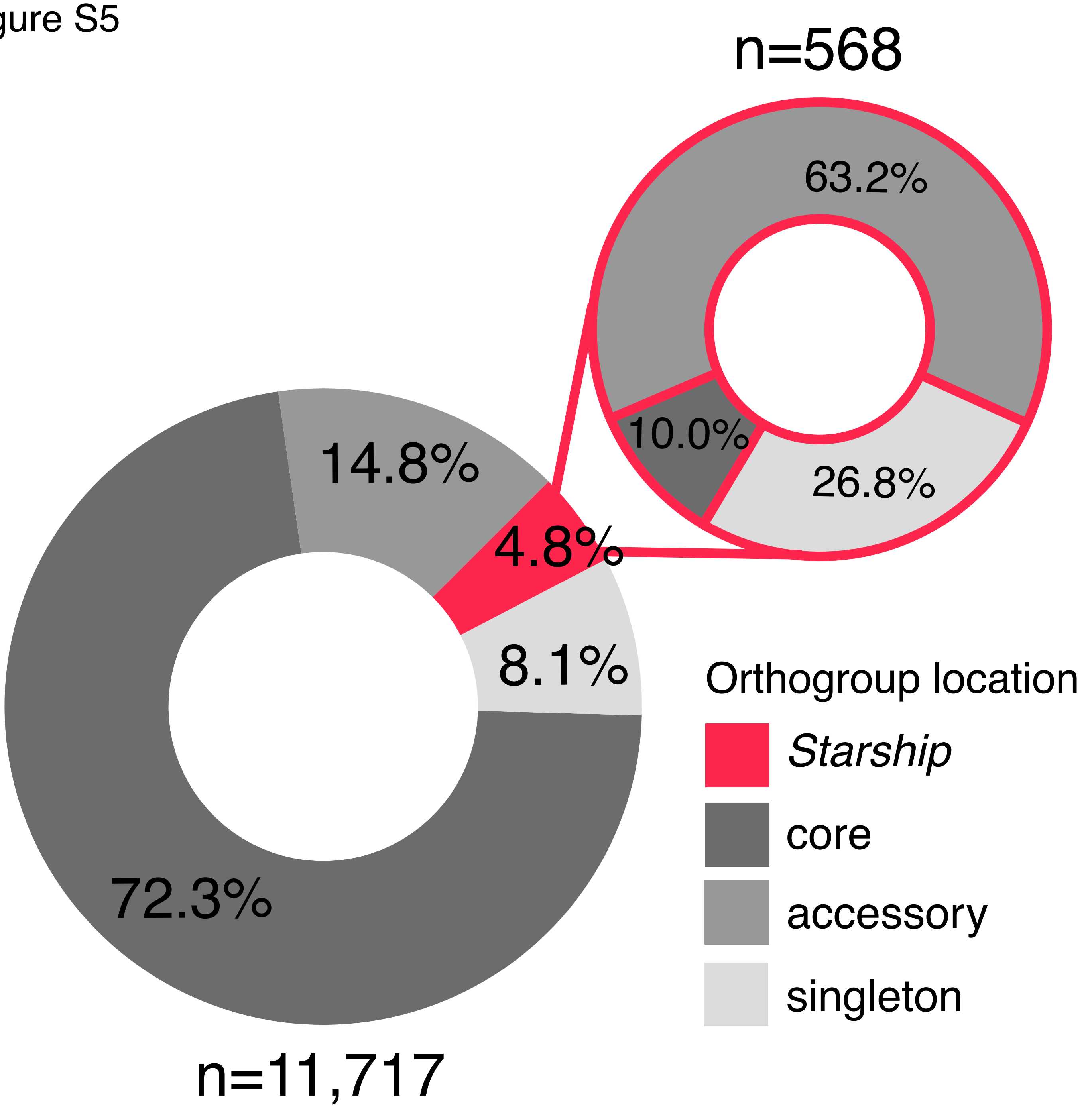
n=568
63.2%
10.0%
26.8%
14.8%
4.8%
8.1%
72.3%
n=11,717
Orthogroup location
Starship
core
accessory
singleton

### Figure S6

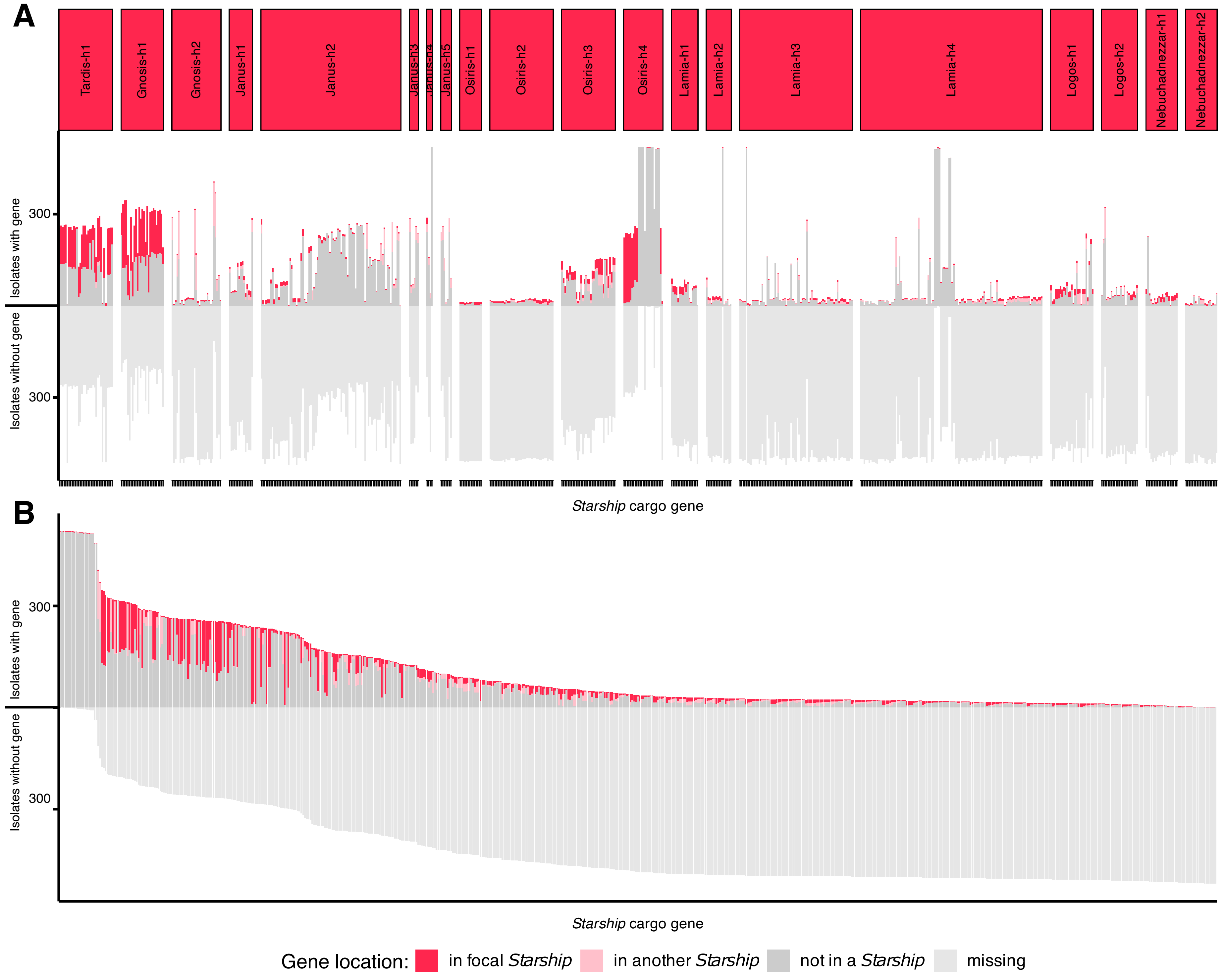

A
Tardis-h1
Gnosis-h1
Gnosis-h2
Janus-h1
Janus-h2
Janus-h3
Janus-h4
Janus-h5
Osiris-h1
Osiris-h2
Osiris-h3
Osiris-h4
Lamia-h1
Lamia-h2
Lamia-h3
Lamia-h4
Logos-h1
Logos-h2
Nebuchadnezzar-h1
Nebuchadnezzar-h2
Isolates with gene
Isolates without gene
300
300
*Starship* cargo gene
B
Isolates with gene
Isolates without gene
300
300
*Starship* cargo gene
Gene location:
in focal *Starship*
in another *Starship*
not in a *Starship*
missing

### Figure S7

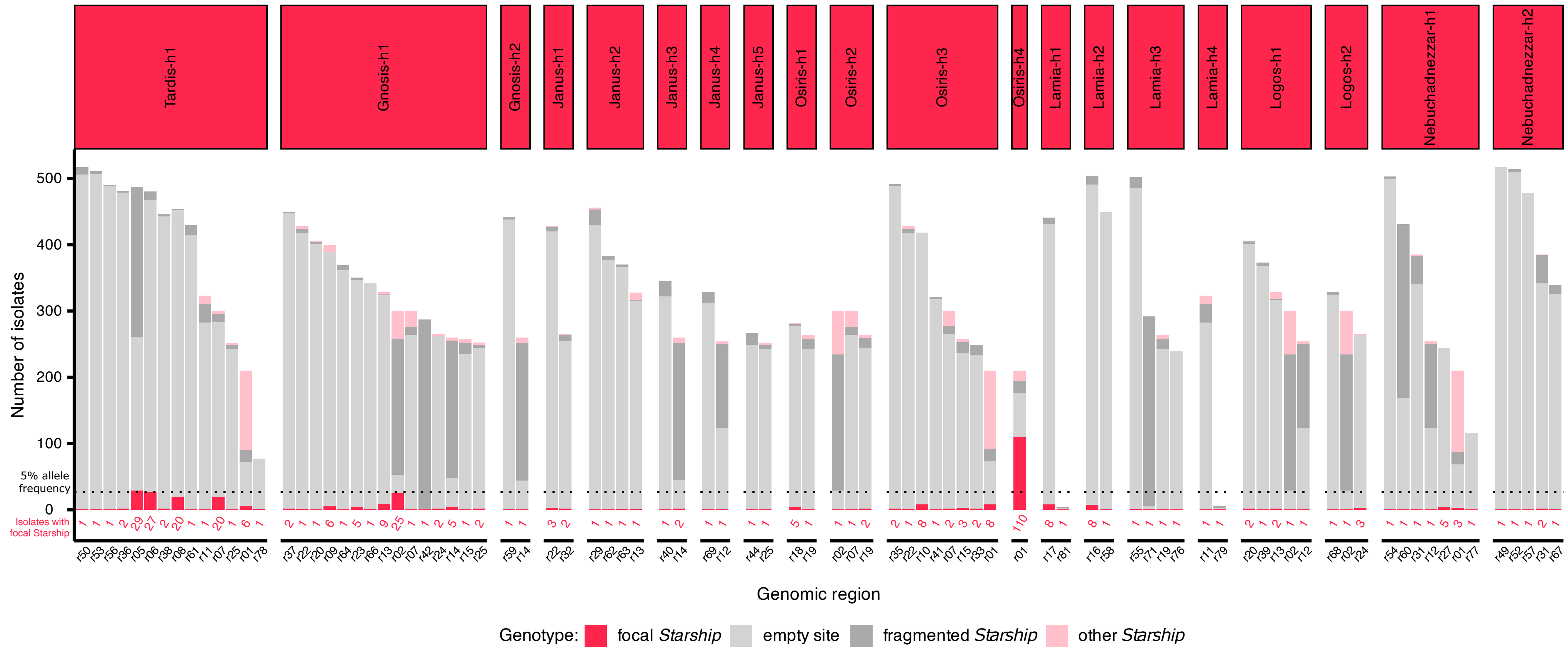
Tardis-h1
Gnosis-h1
Gnosis-h2
Janus-h1
Janus-h2
Janus-h3
Janus-h4
Janus-h5
Osiris-h1
Osiris-h2
Osiris-h3
Osiris-h4
Lamia-h1
Lamia-h2
Lamia-h3
Lamia-h4
Logos-h1
Logos-h2
Nebuchadnezzar-h1
Nebuchadnezzar-h2
Number of isolates
500
400
300
200
100
0
5% allele frequency
Isolates with focal Starship
Genomic region
Genotype: focal Starship empty site fragmented Starship other Starship

Figure S8

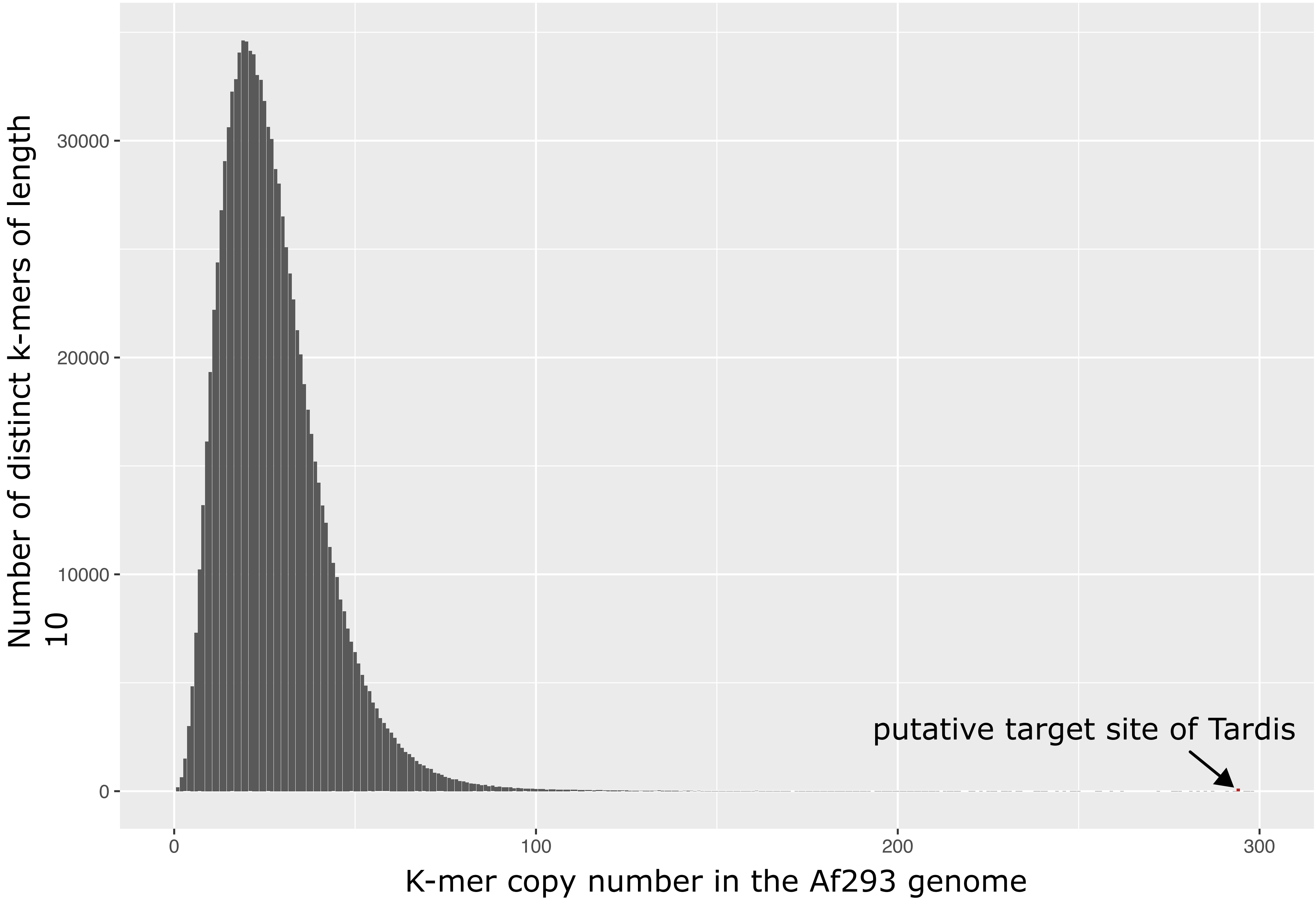

Number of distinct k-mers of length 10
30000
20000
10000
0
0
100
200
300
K-mer copy number in the Af293 genome
putative target site of Tardis

### Figure S9

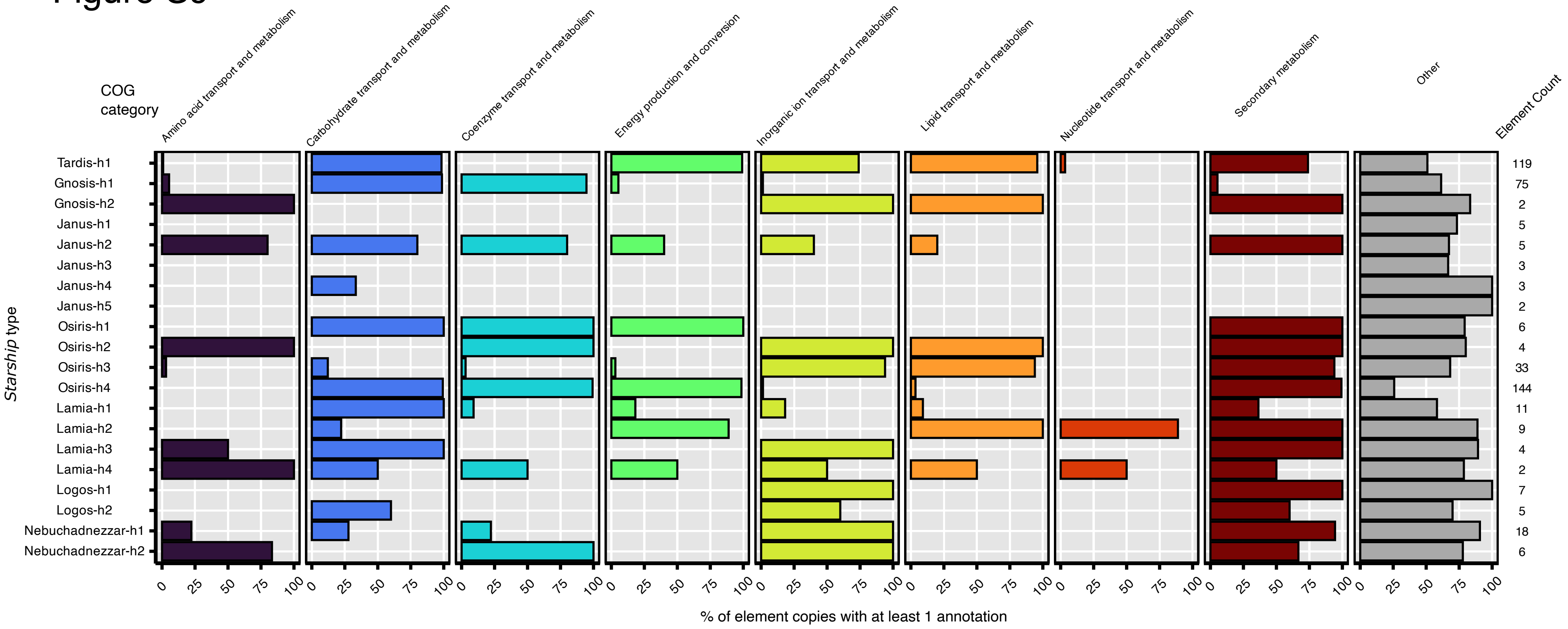
COG category
Amino acid transport and metabolism
Carbohydrate transport and metabolism
Coenzyme transport and metabolism
Energy production and conversion
Inorganic ion transport and metabolism
Lipid transport and metabolism
Nucleotide transport and metabolism
Secondary metabolism
Other
Element Count
Starship type
Tardis-h1
Gnosis-h1
Gnosis-h2
Janus-h1
Janus-h2
Janus-h3
Janus-h4
Janus-h5
Osiris-h1
Osiris-h2
Osiris-h3
Osiris-h4
Lamia-h1
Lamia-h2
Lamia-h3
Lamia-h4
Logos-h1
Logos-h2
Nebuchadnezzar-h1
Nebuchadnezzar-h2
119
75
2
5
5
3
3
2
6
4
33
144
11
9
4
2
7
5
18
6
0 25 50 75 100
% of element copies with at least 1 annotation

### Figure S10

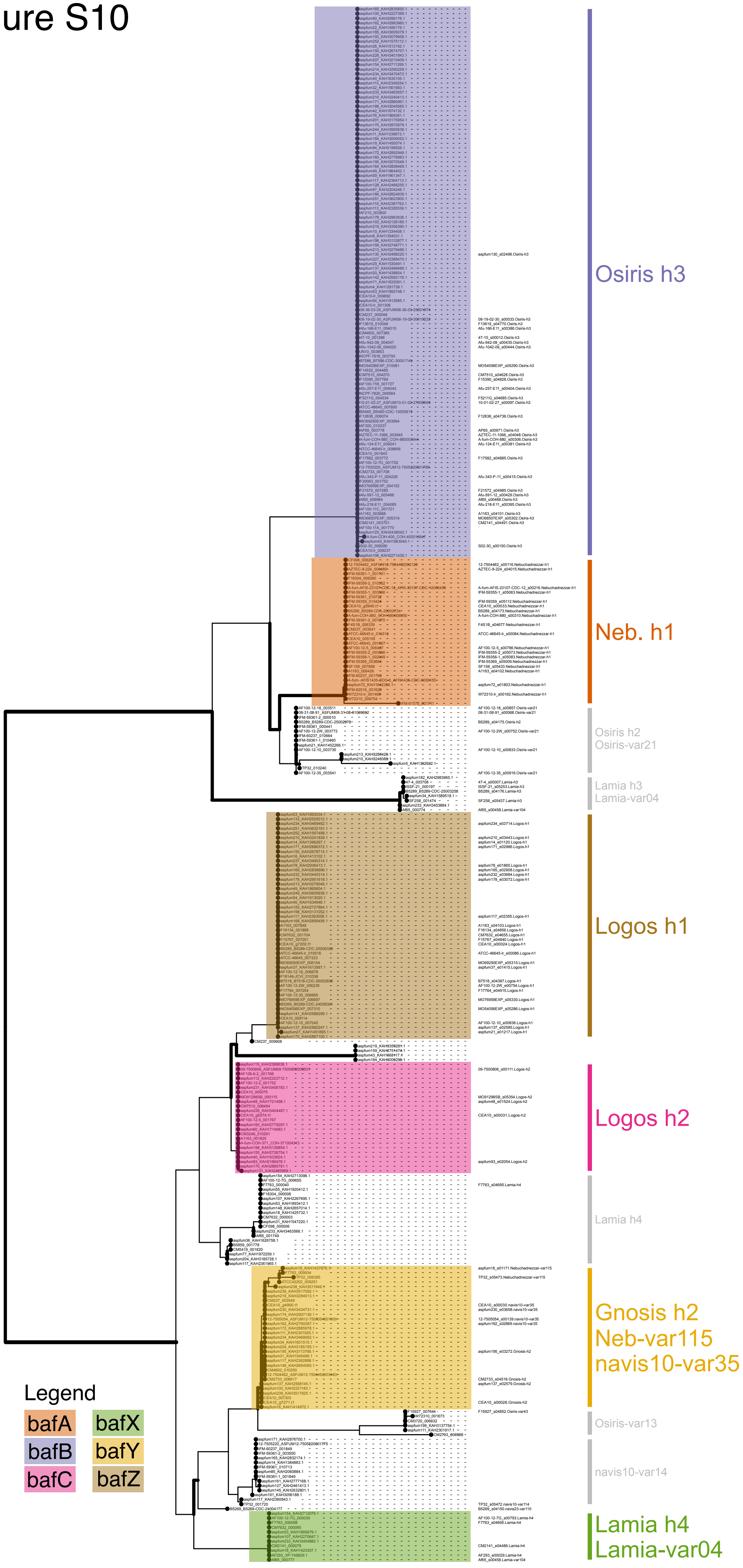
Osiris h3
Neb. h1
Osiris h2
Osiris-var21
Lamia h3
Lamia-var04
Logos h1
Logos h2
Lamia h4
Gnosis h2
Neb-var115
navis10-var35
Osiris-var13
navis10-var14
Lamia h4
Lamia-var04
Legend
bafA
bafX
bafB
bafY
bafC
bafZ

Figure S11

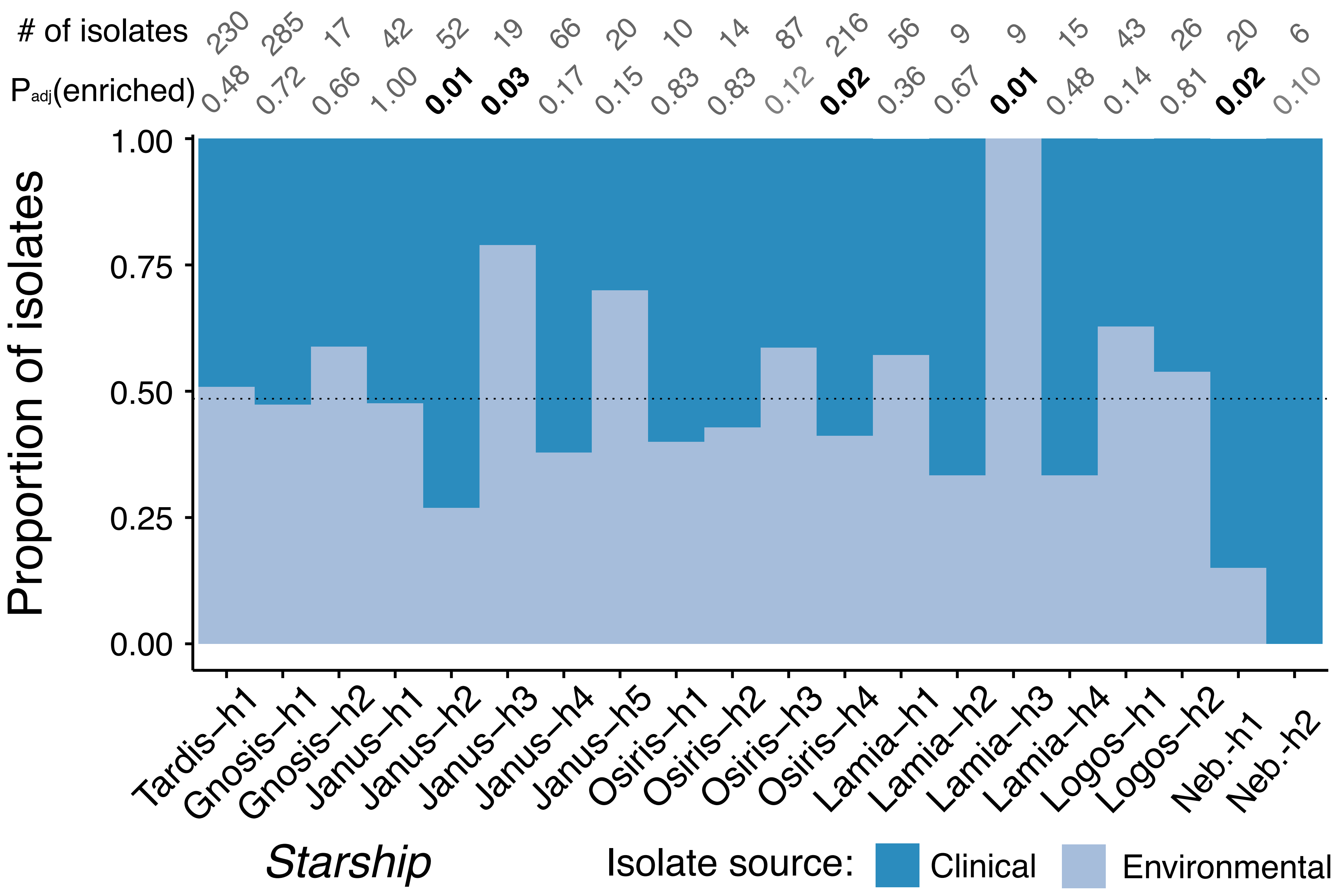

### of isolates
230
285
17
42
52
19
66
20
10
14
87
216
56
9
9
15
43
26
20
6
Padj(enriched)
0.48
0.72
0.66
1.00
0.01
0.03
0.17
0.15
0.83
0.83
0.12
0.02
0.36
0.67
0.01
0.48
0.14
0.81
0.02
0.10
Proportion of isolates
1.00
0.75
0.50
0.25
0.00
Tardis–h1
Gnosis–h1
Gnosis–h2
Janus–h1
Janus–h2
Janus–h3
Janus–h4
Janus–h5
Osiris–h1
Osiris–h2
Osiris–h3
Osiris–h4
Lamia–h1
Lamia–h2
Lamia–h3
Lamia–h4
Logos–h1
Logos–h2
Neb.-h1
Neb.-h2
Starship
Isolate source:
Clinical
Environmental

Figure S12

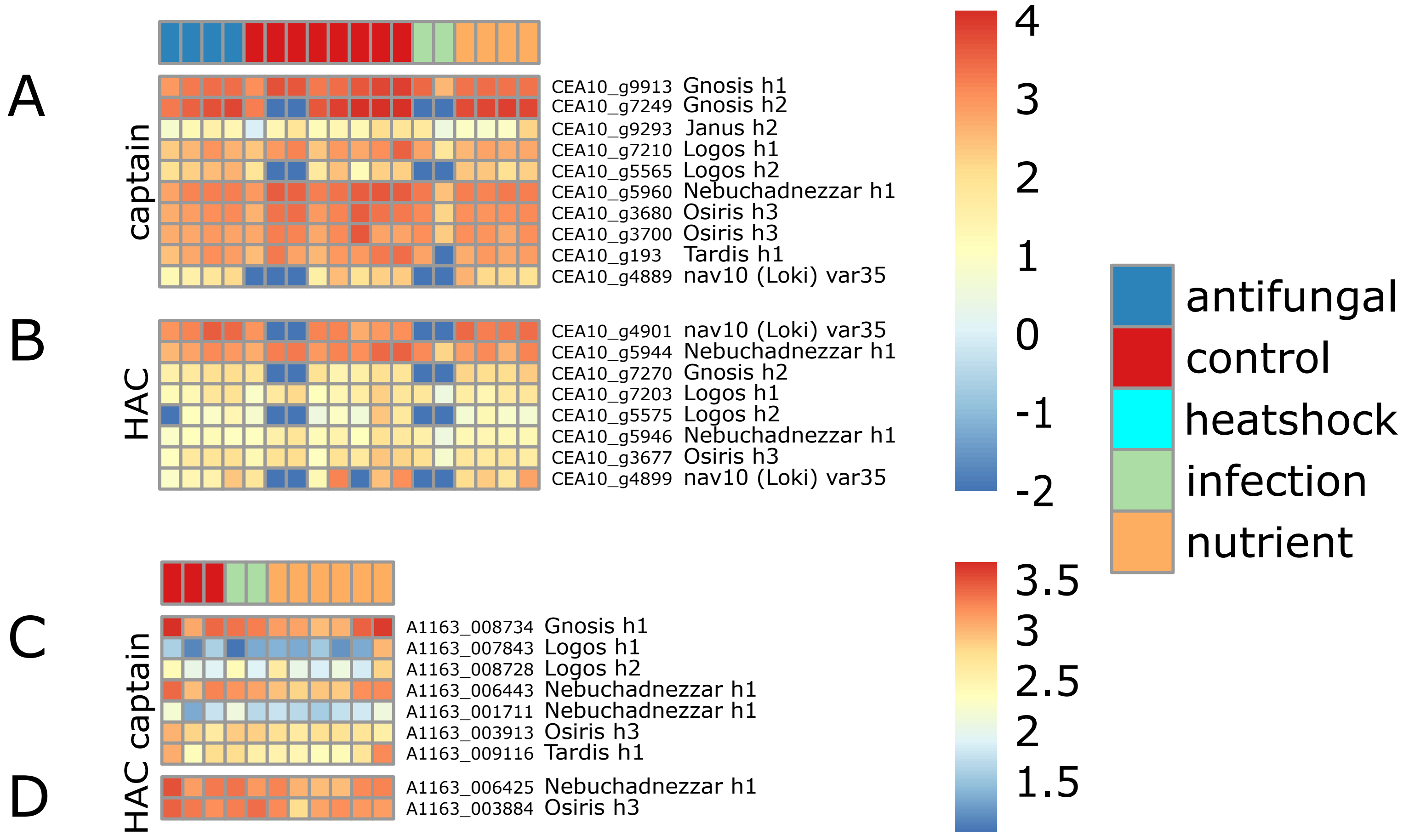
A
B
C
D
captain
HAC
HAC captain
CEA10_g9913 Gnosis h1
CEA10_g7249 Gnosis h2
CEA10_g9293 Janus h2
CEA10_g7210 Logos h1
CEA10_g5565 Logos h2
CEA10_g5960 Nebuchadnezzar h1
CEA10_g3680 Osiris h3
CEA10_g3700 Osiris h3
CEA10_g193 Tardis h1
CEA10_g4889 nav10 (Loki) var35
CEA10_g4901 nav10 (Loki) var35
CEA10_g5944 Nebuchadnezzar h1
CEA10_g7270 Gnosis h2
CEA10_g7203 Logos h1
CEA10_g5575 Logos h2
CEA10_g5946 Nebuchadnezzar h1
CEA10_g3677 Osiris h3
CEA10_g4899 nav10 (Loki) var35
A1163_008734 Gnosis h1
A1163_007843 Logos h1
A1163_008728 Logos h2
A1163_006443 Nebuchadnezzar h1
A1163_001711 Nebuchadnezzar h1
A1163_003913 Osiris h3
A1163_009116 Tardis h1
A1163_006425 Nebuchadnezzar h1
A1163_003884 Osiris h3
4
3
2
1
0
-1
-2
3.5
3
2.5
2
1.5
antifungal
control
heatshock
infection
nutrient

### Figure S13

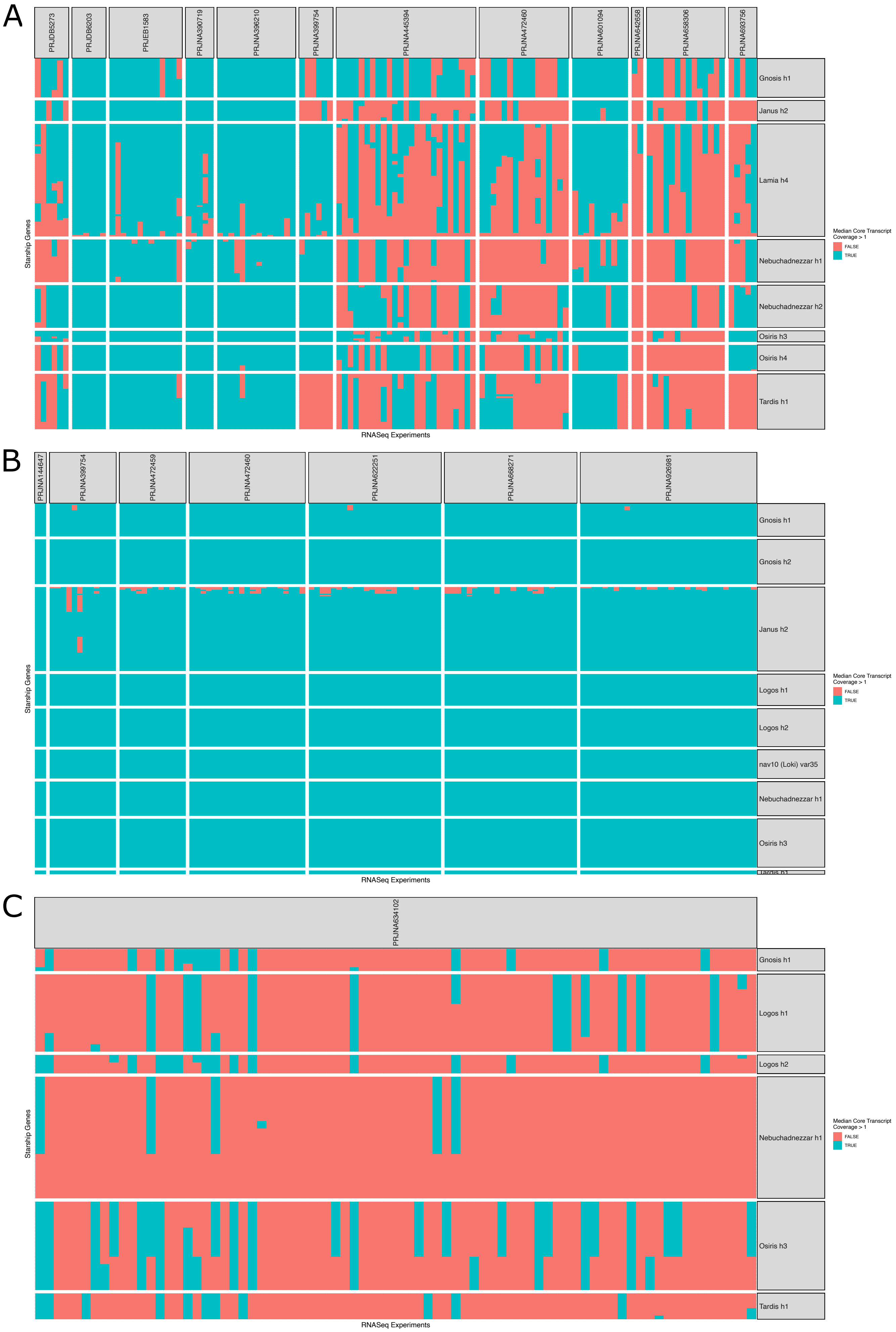
A
PRJDB5273
PRJDB6203
PRJEB1583
PRJNA390719
PRJNA396210
PRJNA399754
PRJNA445394
PRJNA472460
PRJNA601094
PRJNA642658
PRJNA658306
PRJNA693756
Gnosis h1
Janus h2
Lamia h4
Nebuchadnezzar h1
Nebuchadnezzar h2
Osiris h3
Osiris h4
Tardis h1
Starship Genes
RNASeq Experiments
Median Core Transcript Coverage > 1
FALSE
TRUE
B
PRJNA144647
PRJNA399754
PRJNA472459
PRJNA472460
PRJNA622251
PRJNA668271
PRJNA926981
Gnosis h1
Gnosis h2
Janus h2
Logos h1
Logos h2
nav10 (Loki) var35
Nebuchadnezzar h1
Osiris h3
Starship Genes
RNASeq Experiments
Median Core Transcript Coverage > 1
FALSE
TRUE
C
PRJNA634102
Gnosis h1
Logos h1
Logos h2
Nebuchadnezzar h1
Osiris h3
Tardis h1
Starship Genes
RNASeq Experiments
Median Core Transcript Coverage > 1
FALSE
TRUE

### Figure S14

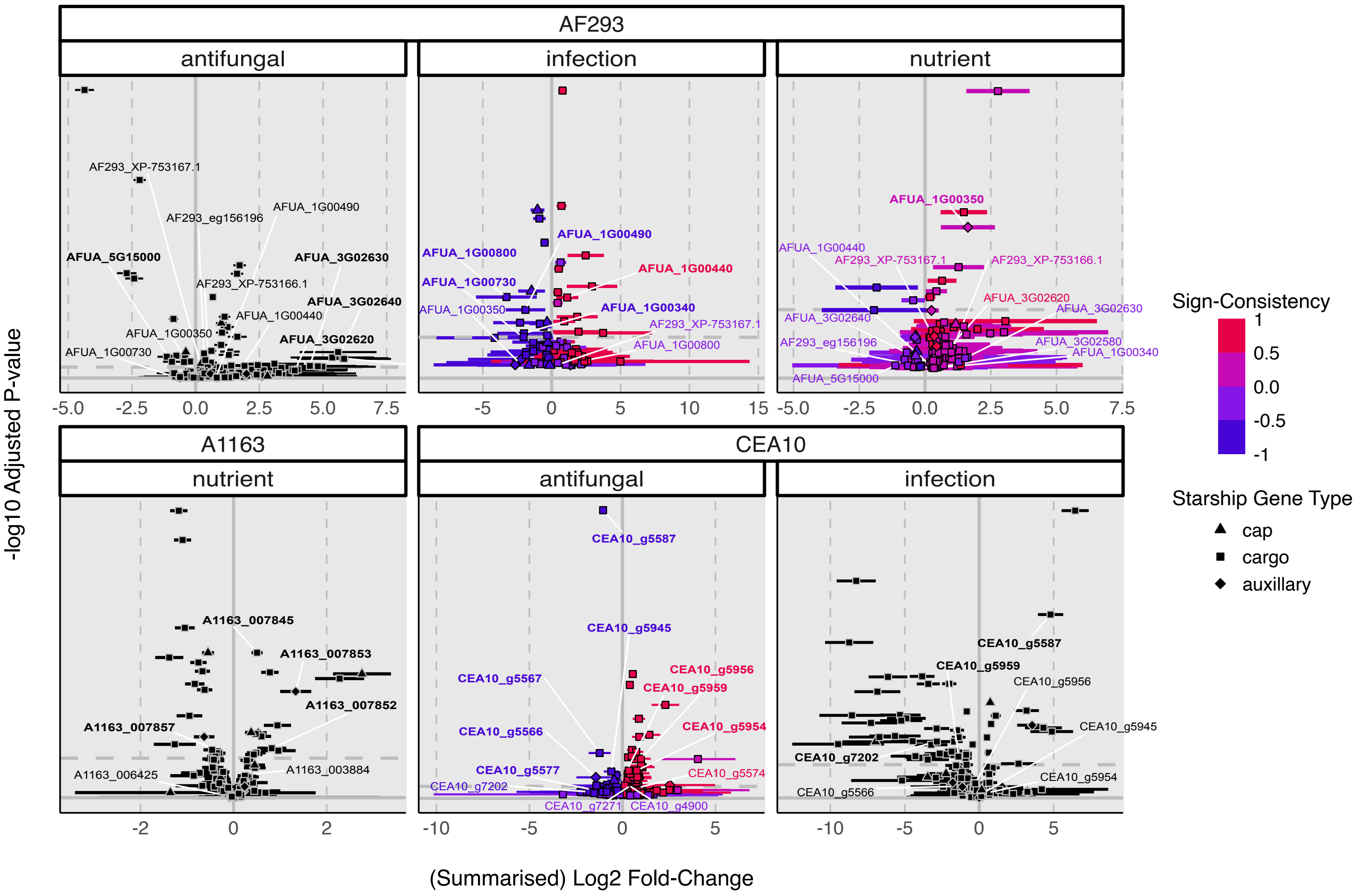

AF293
antifungal
infection
nutrient
A1163
nutrient
CEA10
antifungal
infection
-log10 Adjusted P-value
(Summarised) Log2 Fold-Change
Sign-Consistency
1
0.5
0.0
-0.5
-1
Starship Gene Type
cap
cargo
auxillary
AF293_XP-753167.1
AF293_eg156196
AFUA_1G00490
AFUA_5G15000
AFUA_3G02630
AF293_XP-753166.1
AFUA_3G02640
AFUA_1G00440
AFUA_1G00350
AFUA_3G02620
AFUA_1G00730
AFUA_1G00800
AFUA_1G00340
AFUA_3G02580
AFUA_5G15000
A1163_007845
A1163_007853
A1163_007852
A1163_007857
A1163_006425
A1163_003884
CEA10_g5587
CEA10_g5945
CEA10_g5956
CEA10_g5959
CEA10_g5567
CEA10_g5566
CEA10_g5954
CEA10_g5577
CEA10_g5574
CEA10_g7202
CEA10_g7271
CEA10_g4900

### Figure S15

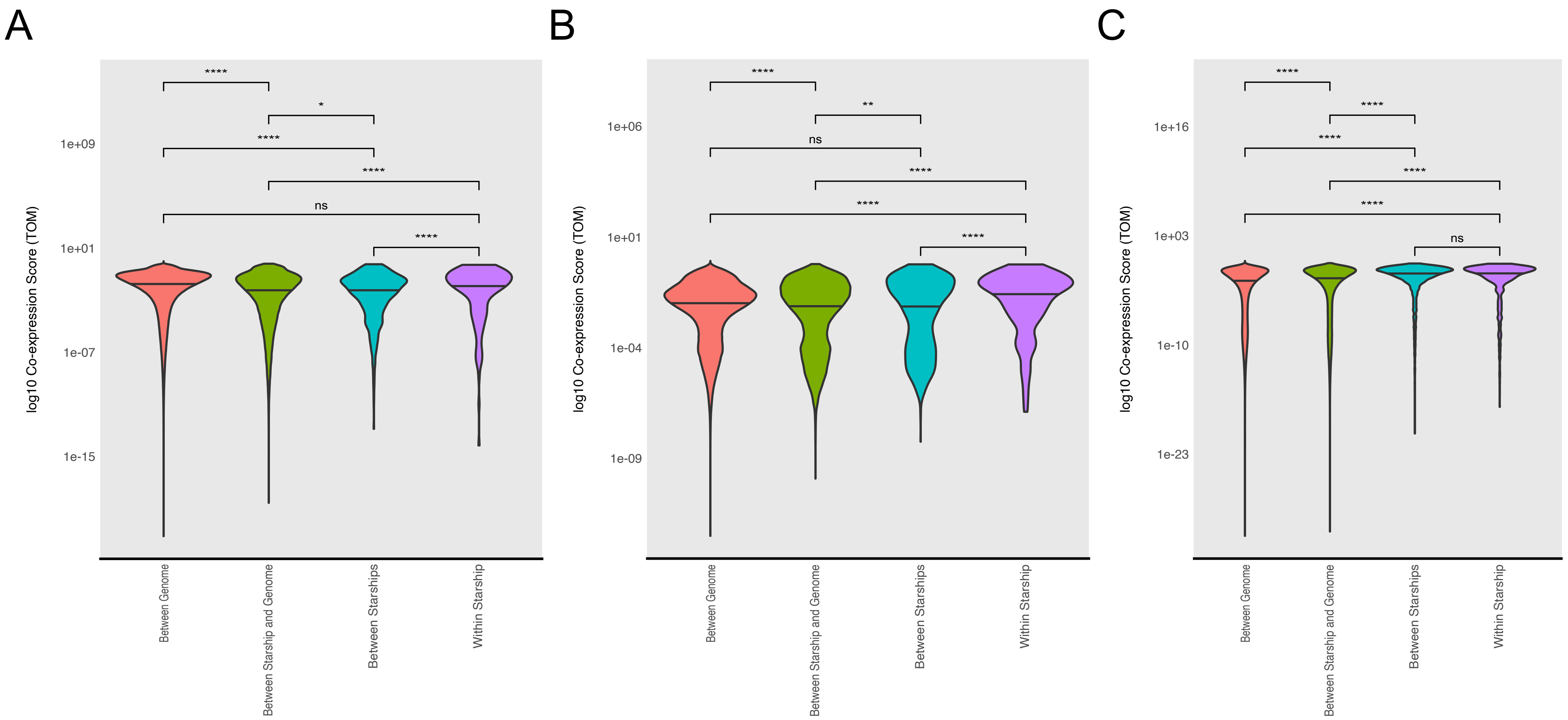
A
B
C
log10 Co-expression Score (TOM)
1e+09
1e+01
1e-07
1e-15
1e+06
1e-04
1e-09
1e+16
1e+03
1e-10
1e-23
****
*
**
ns
Between Genome
Between Starship and Genome
Between Starships
Within Starship

Figure S16

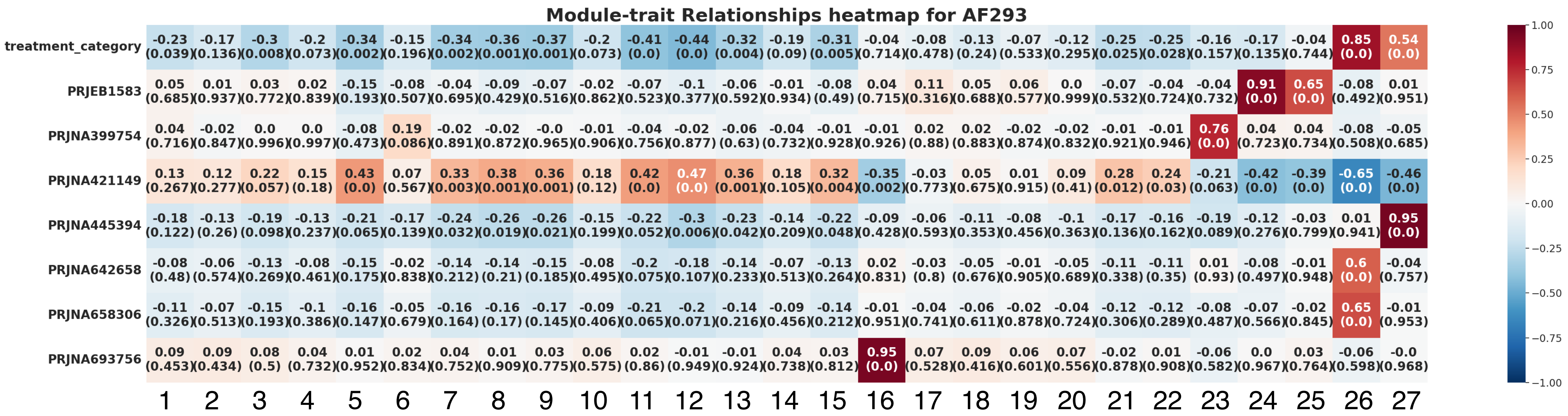
Module-trait Relationships heatmap for AF293
treatment_category -0.23 (0.039) -0.17 (0.136) -0.3 (0.008) -0.2 (0.073) -0.34 (0.002) -0.15 (0.196) -0.34 (0.002) -0.36 (0.001) -0.37 (0.001) -0.2 (0.073) -0.41 (0.0) -0.44 (0.0) -0.32 (0.004) -0.19 (0.09) -0.31 (0.005) -0.04 (0.714) -0.08 (0.478) -0.13 (0.24) -0.07 (0.533) -0.12 (0.295) -0.25 (0.025) -0.25 (0.028) -0.16 (0.157) -0.17 (0.135) -0.04 (0.744) 0.85 (0.0) 0.54 (0.0)
PRJEB1583 0.05 (0.685) 0.01 (0.937) 0.03 (0.772) 0.02 (0.839) -0.15 (0.193) -0.08 (0.507) -0.04 (0.695) -0.09 (0.429) -0.07 (0.516) -0.02 (0.862) -0.07 (0.523) -0.1 (0.377) -0.06 (0.592) -0.01 (0.934) -0.08 (0.49) 0.04 (0.715) 0.11 (0.316) 0.05 (0.688) 0.06 (0.577) 0.0 (0.999) -0.07 (0.532) -0.04 (0.724) -0.04 (0.732) 0.91 (0.0) 0.65 (0.0) -0.08 (0.492) 0.01 (0.951)
PRJNA399754 0.04 (0.716) -0.02 (0.847) 0.0 (0.996) 0.0 (0.997) -0.08 (0.473) 0.19 (0.086) -0.02 (0.891) -0.02 (0.872) -0.0 (0.965) -0.01 (0.906) -0.04 (0.756) -0.02 (0.877) -0.06 (0.63) -0.04 (0.732) -0.01 (0.928) -0.01 (0.926) 0.02 (0.88) 0.02 (0.883) -0.02 (0.874) -0.02 (0.832) -0.01 (0.921) -0.01 (0.946) 0.76 (0.0) 0.04 (0.723) 0.04 (0.734) -0.08 (0.508) -0.05 (0.685)
PRJNA421149 0.13 (0.267) 0.12 (0.277) 0.22 (0.057) 0.15 (0.18) 0.43 (0.0) 0.07 (0.567) 0.33 (0.003) 0.38 (0.001) 0.36 (0.001) 0.18 (0.12) 0.42 (0.0) 0.47 (0.0) 0.36 (0.001) 0.18 (0.105) 0.32 (0.004) -0.35 (0.002) -0.03 (0.773) 0.05 (0.675) 0.01 (0.915) 0.09 (0.41) 0.28 (0.012) 0.24 (0.03) -0.21 (0.063) -0.42 (0.0) -0.39 (0.0) -0.65 (0.0) -0.46 (0.0)
PRJNA445394 -0.18 (0.122) -0.13 (0.26) -0.19 (0.098) -0.13 (0.237) -0.21 (0.065) -0.17 (0.139) -0.24 (0.032) -0.26 (0.019) -0.26 (0.021) -0.15 (0.199) -0.22 (0.052) -0.3 (0.006) -0.23 (0.042) -0.14 (0.209) -0.22 (0.048) -0.09 (0.428) -0.06 (0.593) -0.11 (0.353) -0.08 (0.456) -0.1 (0.363) -0.17 (0.136) -0.16 (0.162) -0.19 (0.089) -0.12 (0.276) -0.03 (0.799) 0.01 (0.941) 0.95 (0.0)
PRJNA642658 -0.08 (0.48) -0.06 (0.574) -0.13 (0.269) -0.08 (0.461) -0.15 (0.175) -0.02 (0.838) -0.14 (0.212) -0.14 (0.21) -0.15 (0.185) -0.08 (0.495) -0.2 (0.075) -0.18 (0.107) -0.14 (0.233) -0.07 (0.513) -0.13 (0.264) 0.02 (0.831) -0.03 (0.8) -0.05 (0.676) -0.01 (0.905) -0.05 (0.689) -0.11 (0.338) -0.11 (0.35) 0.01 (0.93) -0.08 (0.497) -0.01 (0.948) 0.6 (0.0) -0.04 (0.757)
PRJNA658306 -0.11 (0.326) -0.07 (0.513) -0.15 (0.193) -0.1 (0.386) -0.16 (0.147) -0.05 (0.679) -0.16 (0.164) -0.16 (0.17) -0.17 (0.145) -0.09 (0.406) -0.21 (0.065) -0.2 (0.071) -0.14 (0.216) -0.09 (0.456) -0.14 (0.212) -0.01 (0.951) -0.04 (0.741) -0.06 (0.611) -0.02 (0.878) -0.04 (0.724) -0.12 (0.306) -0.12 (0.289) -0.08 (0.487) -0.07 (0.566) -0.02 (0.845) 0.65 (0.0) -0.01 (0.953)
PRJNA693756 0.09 (0.453) 0.09 (0.434) 0.08 (0.5) 0.04 (0.732) 0.01 (0.952) 0.02 (0.834) 0.04 (0.752) 0.01 (0.909) 0.03 (0.775) 0.06 (0.575) 0.02 (0.86) -0.01 (0.949) -0.01 (0.924) 0.04 (0.738) 0.03 (0.812) 0.95 (0.0) 0.07 (0.528) 0.09 (0.416) 0.06 (0.601) 0.07 (0.556) -0.02 (0.878) 0.01 (0.908) -0.06 (0.582) 0.0 (0.967) 0.03 (0.764) -0.06 (0.598) -0.0 (0.968)
1 2 3 4 5 6 7 8 9 10 11 12 13 14 15 16 17 18 19 20 21 22 23 24 25 26 27
1.00 0.75 0.50 0.25 0.00 −0.25 −0.50 −0.75 −1.00
